# Characterizing developmental changes in infant habituation using functional change point detection

**DOI:** 10.64898/2025.12.03.692095

**Authors:** Samuel Beaton, Samantha McCann, Sarah Lloyd-Fox, Clare E. Elwell, Ebrima Mbye, Anna Blasi Ribera, Sophie E. Moore

## Abstract

**Significance:** Habituation is an early-developing cognitive process linked to learning, typically reflected in functional near-infrared spectroscopy (fNIRS) studies as a reduction in evoked hemodynamic response amplitude. Conventional fNIRS analyses rely on linear time-invariance assumptions, potentially limiting insight into how responses evolve across repeated stimulus presentations.

**Aim:** To determine whether functional change point (FCPt) detection can characterize trial-specific changes in the infant hemodynamic response and reveal developmental differences in habituation timing.

**Approach:** Functional data analysis (FDA) treats entire response curves as statistical objects. Within this framework, FCPt detection identifies statistically significant structural shifts in the mean response across trials. FCPt detection with wild binary segmentation was applied to infant fNIRS data (*n* = 204) collected from infants living in rural Gambia at 5-, 8-, and 12-months of age during a habituation and novelty detection paradigm.

**Results:** Significant changes were identified within auditory cortical regions. A high proportion of detected change points corresponded to decreases in response magnitude at 8- and 12-months. Ordinal regression revealed an age-related shift toward earlier occurrence of decreasing change points, indicating that older infants completed habituation sooner within the trial sequence.

**Conclusion:** FCPt detection provides temporal information on the infant hemodynamic response unavailable to conventional analysis. This additional information has revealed a previously overlooked developmental change in the habituation response during the first year of infancy.

## 1 Introduction

### 1.1 Rationale

Habituation is an early-developing, low-level process of reducing attention to repeated, neutral stimuli [1], [2], conserving cognitive resources and potentially aiding survival as a consequence. It is linked to novelty response (‘dishabituation’), an increased response to a novel stimulus, which may be unpleasant or hazardous [3]. Habituation is thought to direct attention to support early learning, perhaps through the construction of internal representations of repeated stimuli [2], [4].

Habituation paradigms are suitable for infant neuroimaging research since they do not rely on overt behaviors and the outcome of interest is cognitive rather than a behavioral change [4]; see Nordt and colleagues [5] for a detailed discussion. Indeed, past infant functional near-infrared spectroscopy (fNIRS) research shows that the evoked infant cortical hemodynamic response can gradually decrease when stimuli are repeated [2], [6], [7], [8]. However, fNIRS data analyses for task-based experimental paradigms typically rely on averaging the signal across trials – time blocks concurrent with a task or stimulus – to reduce noise [9]. This is based on the linear time-invariance (LTI) assumption [10] that hemodynamic responses remain constant across trials and sum linearly with other physiological phenomena to give the recorded signal [11].

As an example, habituation in hemodynamic responses have previously been identified during an auditory fNIRS habituation task in the first year of life in infants recruited from the UK and The Gambia as part of the BRIGHT study [4], [8], [12]. The observed habituation of Gambian infants persisted even when introducing a novel stimulus intended to elicit a heightened response, indicating continued habituation. Given this lack of identifiable ‘novelty response’, the ability to distinguish between habituation trajectories of the hemodynamic response takes on increased importance. However, these previous results were generated using block averaging methods [13] to determine not only channels of activation but also condition contrasts between groups (‘epochs’) of 5 trials; see Lloyd-Fox and colleagues [8] for more information. Whilst informative, this approach overlooks the trial-specific and temporal information necessary to examine the gradual change in the hemodynamic response during the session. It may additionally overlook novelty detection if the associated increase in hemodynamic response occurs and subsides within the 5-trial epoch. Investigation of trial-by-trial change in the hemodynamic response may permit insight into different rates of habituation between ages: what appears to be an overall reduction in response across trial epochs at older ages may instead be the manifestation of a faster reduction in response in early epochs when averaged across a sufficiently large number of trials.

Functional data analysis (FDA) – a branch of statistics concerned with the analysis of mathematical functions as the core objects [14] – may enable such trial-by-trial analysis. This in turn could permit differentiation between the habituation trajectories of evoked hemodynamic responses: particularly useful in collected data where there is an absence of a clear novelty-response, but also useful in scenarios where the novelty response is exhibited, as an additional means to assess these two closely related phenomena. Most importantly, the ability to assess the timing of decreases in a repeated, evoked response – where “timing” refers to the trial in the experimental paradigm at which a change in the mean hemodynamic response occurs – would permit analysis into the rate at which infants process repeated, familiar stimuli more generally. This would thereby serve as an early measure of learning and providing insight into an important early-life cognitive process.

Full overviews of FDA can be found in works by Kokoszka and Reimherr [15], Ramsay and Silverman [16] and Müller [17], but relevant concepts are outlined briefly here for clarity, with additional detail found in Supplementary Materials 1. Within FDA, discrete data can represent a regular sampling of a continuous, smooth underlying process allowing the function itself to be treated as the object of interest [16], [18]. In practice, therefore, the interchangeable terms ‘functional objects’ or ‘functions’ often represent – and refer to – what we know as curves, in the intuitive sense. They can accordingly be used to study phenomena where data vary continuously and smoothly in time or space and can be represented by such curves. This representation reduces concerns about correlation between repeated measures and accounts for latent temporal structure [19], [20].

Observed or measured functions, *X*_*n*_, can be seen as realisations (samples) of a random function, 𝒳, leading to the concepts of mean and covariance functions [17]. When functions are observed repeatedly in an ordered manner, they may be modeled as a functional time series: a sequence of functional observations indexed by order (typically time) [21]. The group of approaches which can be used to determine points in functional time series where these structural properties change (see Figure 1) is known as functional change point (FCPt) detection [22].

**Figure 1:**
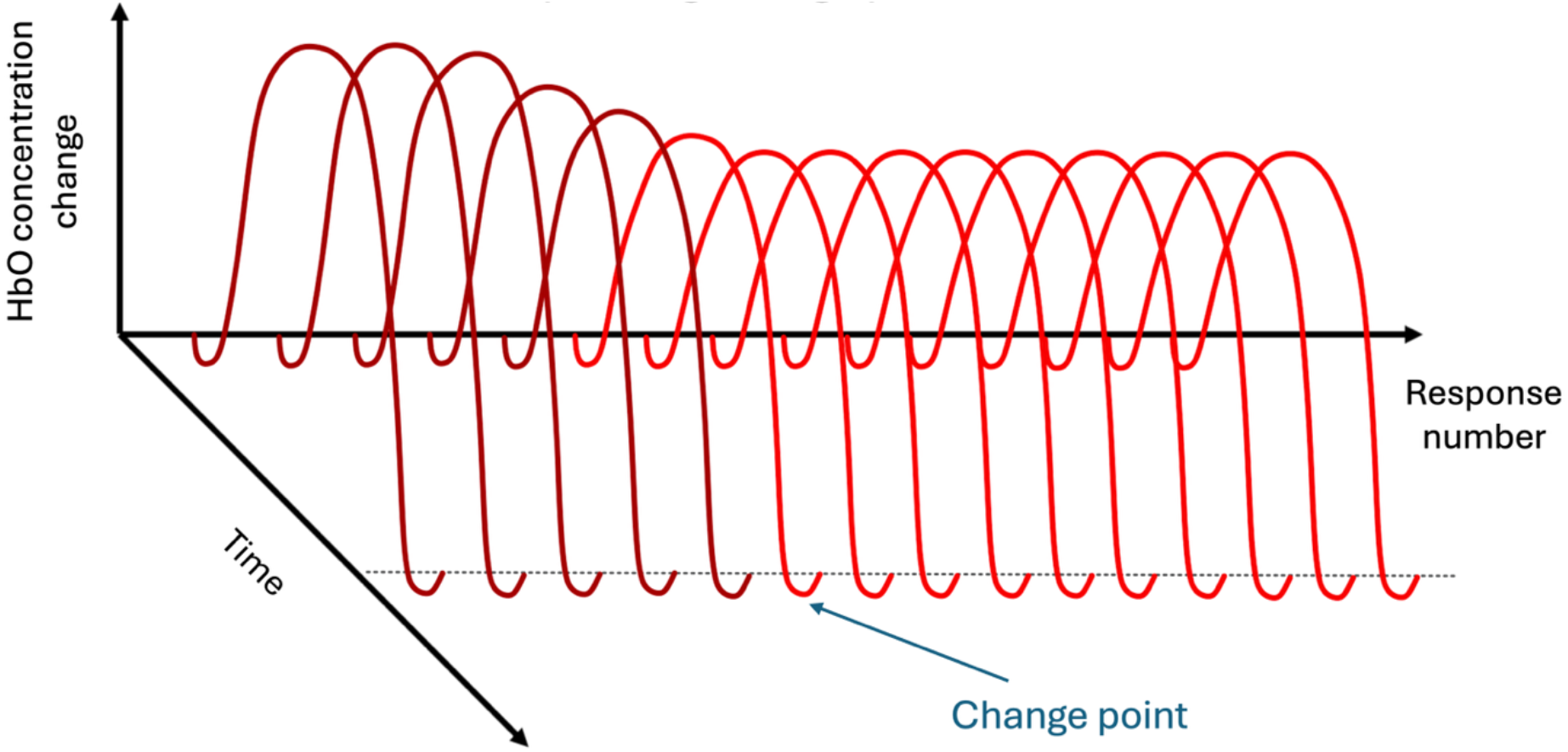
Hypothesized habituation of an evoked hemodynamic response and a corresponding example change point (FCPt). Hypothesized FCPts occur when significant structural change is present in the functional representation of the hemodynamic response; they are represented at the points by a change in color of the hemodynamic response functions. The set of consecutive evoked hemodynamic responses in this image forms a functional time series of length 15 i.e. a series of 15 functional objects.

During FCPt detection, a change point occurs when there is a significant difference in some structural property of interest – this property is often the mean function. A detected change point signifies that the collection of observed functions before and after the point differ enough to be seen as sampling two different underlying random functions 𝒳_*B*_ and 𝒳_*A*_, respectively. Detected changes may be global (i.e. across the whole mean function) or localized; indicate trends or shifts in the structure (shape) or mean level of the function.

In repeated-stimulus fNIRS paradigms, each stimulus presentation is designed to yield a hemodynamic response function, and the ordered collection of these response functions forms a functional time series. Furthermore, during habituation changes in the mean of the hemodynamic response are expected, alongside consistency in variance and shape (i.e. the response maintains a canonical hemodynamic profile). This is important because detection of functional change points requires a sufficiently stable underlying structure against which statistically significant deviations can be identified. This likely ensures that only change relating to the stimulus-locked hemodynamic response will be detected, since it is anticipated that this will be the one of the only data-generating processes which is consistent across trials.

Here, we propose to apply FCPt detection to investigate the change in functional representations of the evoked hemodynamic response, collected from infants using fNIRS as part of an auditory habituation paradigm. Previous applications of FCPt detection to identify changes in the mean function in neuroimaging data have examined epidemic changes – where the mean function returns to its original, pre-change state – in fMRI data [23], [24], [24]. However, to the authors’ knowledge, no work has utilized FCPt detection on either infant neuroimaging or fNIRS data as yet, making this the first such example of either case.

FCPt detection can be conducted using several methods. One common approach to FCPt detection is to first perform dimension reduction, using functional principal component analysis (FPCA), a decomposition of the data analogous to ordinary principal components analysis [15, p. 41]. However, structural change in modes orthogonal to the FPCs may be undetected using this approach [25]. In contrast, fully functional (FF) change point detection methods, which utilize the data in their entirety, may be more effective, though they require more computational resources.

This work aims to demonstrate that habituation can be modeled as a significant structural change in the hemodynamic response, as identified using FCPt detection applied to infant fNIRS data. It also aims to show that this approach permits the investigation of the timing of habituation and that this timing changes throughout infancy, providing valuable insight into an important early-life cognitive process.

### 1.2 Objectives

This work has two objectives:

a. To explore the ability of FCPt detection to obtain greater insight into the habituating infant hemodynamic response than has been previously possible using established approaches
b. Use FCPt detection to infer longitudinal change in evoked infant habituation responses

With respect to these objectives, the following hypotheses were formulated:

1. FCPt detection will provide information on the timing of changes in hemodynamic response structure related to habituation in a way which univariate approaches are unable to
2. If age-associated differences exist in the timing of habituation, older participants will exhibit earlier reductions in hemodynamic response and/or completion of the habituation process

## 2 Methods

### 2.1 Participants

Data described herein were collected as part of the Brain Imaging for Global HealTh (BRIGHT) Project, which aimed to expand diversity in neurodevelopmental studies, establish brain function-for-age curves in infants from high-(UK) and low-resource (The Gambia) settings and explore the effects that contextual factors related to living in a low-resource context may have on infant development [26].

204 infants were enrolled into the BRIGHT Project in The Gambia. Families were identified using the West Kiang Demographic Surveillance System [27], primarily from the village of Keneba and neighboring villages in the rural West Kiang District. In West Kiang, the predominant (79.9%) ethnicity is Mandinka [27] therefore recruitment was limited to this group to avoid confounds related to study protocol translation. Ethical approval was given by the joint Gambia Government-MRCG Ethics Committee (SCC 1451v2). Prior to participation, informed consent was obtained from all parents/carers in writing or via thumbprint if they were unable to write. Mothers were initially seen at a 34-36 week antenatal visit and assessment study visits took place at 7-14 days, 1-, 5-, 8-, 12-, 18- and 24mo of age thereafter. For the current analysis, data collected at 5-, 8- and 12mo of age (hereafter referred to as *x*mo for *x*-months of age) is used.

Infants born prior to 36 weeks gestational age were excluded from BRIGHT, however a minimum birth weight was not specified for Gambian infants. Anthropometric measures (head circumference, weight, and length/height) were collected and used to calculate z-scores. These included head circumference (HCZ), weight-for-height (WHZ), weight-for-age (WAZ) and height-for-age (HAZ). Low WHZ, WAZ and HAZ scores (< 2 standard deviations below the WHO reference) indicate that a child is wasted, underweight, or stunted, respectively.

### 2.2 Experimental procedures

#### 2.2.1 Data collection

Testing took place between June 2016 and July 2019 at the Medical Research Council Unit The Gambia at the London School of Hygiene and Tropical Medicine (MRCGLSHTM; www.mrc.gm).

Custom-made headgear was fitted after head measurements (head circumference, and ear-to-ear both around the forehead and over the top of the head) had been taken to aid with the alignment of fNIRS headgear with the 10/20 system anatomical landmarks. The headgear was designed to record responses bilaterally from auditory associative brain regions (see Figure 2), including the inferior frontal gyrus (IFG), middle and superior temporal regions, and the tempero-parietal junction [8], [28], [29]. The fNIRS array comprised 17 channels in each hemisphere with a source-detector (S-D) distance of 2cm, corresponding to a penetration depth of ∼1cm from the scalp surface and thus permitting measurement of the gyri and superficial sulci [8]. fNIRS data were collected with the NTS optical topography system (Gowerlabs Ltd., UK), using source wavelengths of 850 and 780nm).

**Figure 2:**
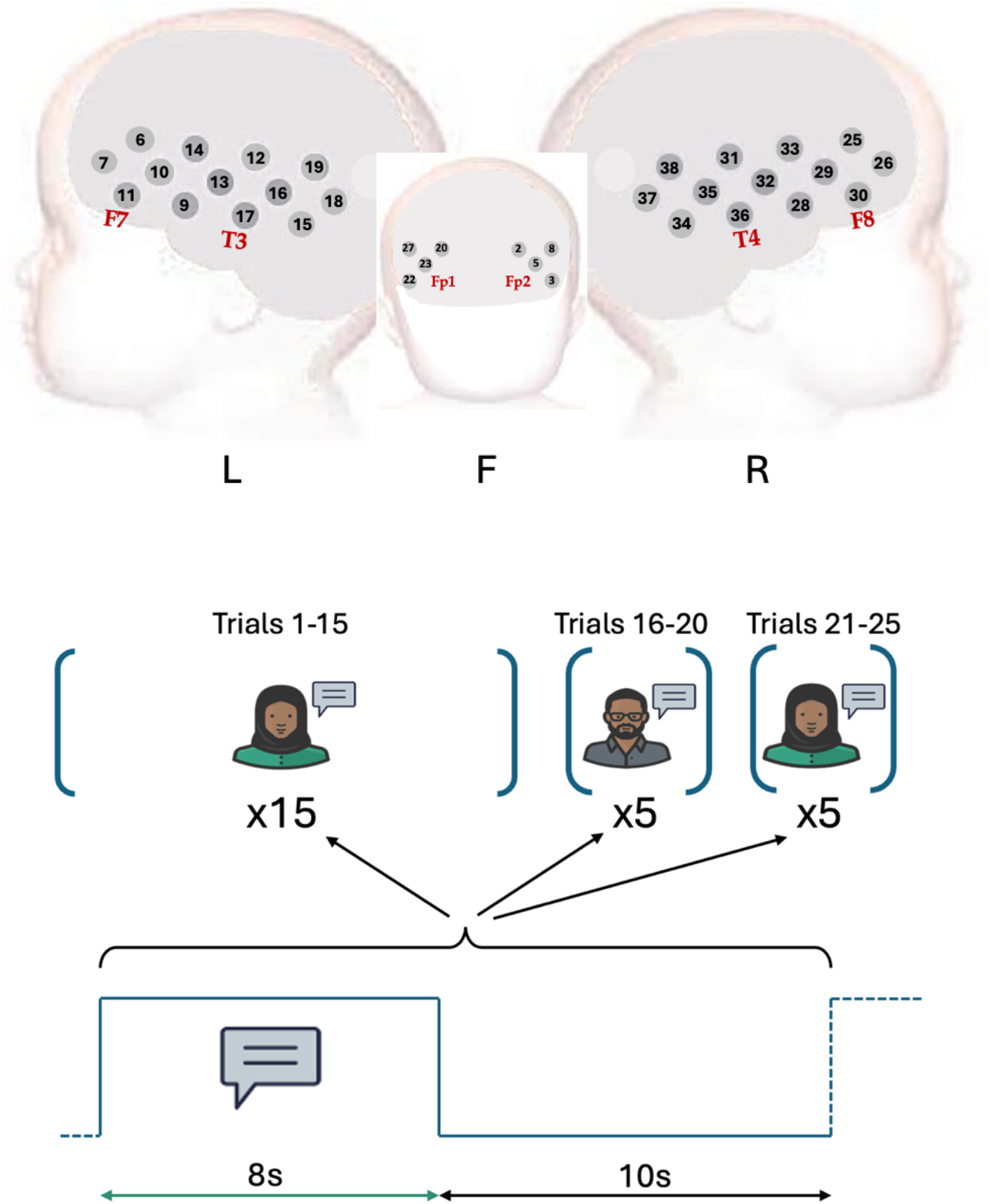
Array layout and experimental paradigm during data collection. Top: array layout during data acquisition. Bottom: The HaND paradigm in the BRIGHT Project, with 25 repetitions of an identical spoken 8s sentence, followed by a 10s inter-stimulus interval. Trials 1-15 and 21-25 have the same speaker; in trials 16-20 the identity and sex of the speaker is changed.

During data acquisition, infants sat on a carer’s lap. Carers were discouraged from interacting with the infant to minimise risk of confounding stimuli. However, infants’ attention was engaged by (non-social, non-auditory) bubble-blowing and silent demonstration of soft toys, which helped minimise infant head movement. The Habituation and Novelty Detection (HaND) task was part of a larger battery of fNIRS assessments with a total recording time of at least ∼21min 30sec (6min social task; 4min functional connectivity data acquisition; 7min 30sec HaND; 4min further functional connectivity) [8], [30], [31]. The protocol for 8- and 12mo infants was even longer, as it also included a working memory and deferred imitation tasks. Where possible, tasks were completed uninterrupted; sessions were paused and subsequently resumed in the event of infant discomfort.

#### 2.2.2 The HaND paradigm

The aim of this paradigm is to enable measurement of habituating hemodynamic responses related to a familiar stimulus and subsequent increase in response related to a novel stimulus [8].

The experimental paradigm consisted of 25 trials, each consisting of the same spoken 8sec sentence in Mandinka followed by 10sec of silence - traditionally considered the minimum amount of time necessary for the hemodynamic response to return to baseline (see Figure 2, bottom). Working with a battery of multiple tasks with infant participants meant time allocation for tasks was constrained, so the inter-stimulus interval (ISI) was chosen to be the bare minimum possible whilst still theoretically enabling a return to baseline after the hemodynamic response.

The same recording, with a female voice, was used for trials 1-15 (the Habituation trials); a different, male voice was used for trials 16-20 (the Novelty trials); finally, the original, female recording was again used for trials 21-25 (the Post-Test trials). The sentence in Mandinka: *“Denano a be nyadii. I be kongtan-rin? Abaraka bake ela naa kanan njibee bee, n kontanta bake le ke jeh”* roughly translates to: *“Hi baby! How are you? Are you having fun? Thank you for coming to see us today. We’re very happy to see you”*. Technical detail on the recording, processing and playback of the auditory stimulus can be found in previous work by Lloyd-Fox and colleagues [8] and Katus and colleagues [4].

### 2.3 Data processing

First, participants missing one or more of the familiarisation epochs due to insufficient compliance with the experimental task (based on offline coding of infant behaviour from video recording of the session) were excluded from the analysis. Following this, channels with poor optode coupling were pruned for remaining infant datasets. Initially channels were removed if the signal was weak or saturated, indicated by a mean signal amplitude less than 3 × 10^−4^ or greater than 1 × 10^7^, respectively based on the NTS system specifications. Subsequently, the coefficient of variation was calculated on a pre-selected motion-free section of the data for remaining channels as *CV* = *σ*/|*μ*|, where *σ* and *μ* are the standard deviation and mean of the light intensity signal in a channel, respectively [33]. Channels were pruned if this value was below 8% [7]. Infants were excluded if more than 40% of channels were pruned [8]. Channel signals were then converted from light attenuation to optical density, enabling motion artifact correction using the Homer2 software package [34].

Motion artifact correction was conducted using the spline interpolation approach [35] followed by wavelet denoising [36]. Spline interpolation corrects pre-identified motion artifacts channel-by-channel by estimating the true signal based on neighboring data, via cubic spline interpolation. The spline estimate is subtracted from the recorded signal to remove slow drift data components; after this step, signal before and after the artifact can be aligned, which maintains signal continuity to correct baseline shifts caused by motion [37]. Motion artifacts were identified using *hmrMotionArtifactByChannel* with parameters *tMotion = 1, tMask = 1, STDEVthresh = 15* and *AMPthresh = 0*.*4*. Following previous findings and the developers’ recommendations, a parameter of *p* = 0.99 was used for spline interpolation [35], [38]. Wavelet denoising decomposes the signal into constituent wavelets, removing those with characteristics corresponding to abrupt changes (e.g. spikes) caused by motion, before subsequent reconstruction. An interquartile range factor of 0.8 was implemented for the function *hmrMotionCorrectWavelet* [38]. Persistent motion artifacts were subsequently identified using *hmrMotionArtifactByChannel* as above; trials within 4 seconds after or 12 seconds before detected motion artifacts were removed. Channels with fewer than 60% (3/5) valid trials in trials 1-5, trials 6-10, or trials 11-15 were removed, to maintain a sufficient number of trials in order to accurately analyze the evoked hemodynamic response during the habituation stage of the paradigm. Participant exclusion was conducted again, with infants excluded if fewer than 60% channels remained after channel pruning and motion artifact removal.

Consistent with previous work with this data, a low-pass filter was then applied with a 0.8Hz frequency cutoff [12]. The low-pass filter removes high-frequency oscillations associated with physiological confounds such as the participant heart rate. The optical density data was then converted to oxy-(HbO) and deoxyhemoglobin (HbR) data using the modified Beer-Lambert law and an age-appropriate differential pathlength factor to account for age-related tissue composition changes [39], [40]. Finally, concentration data was linearly detrended [8] by fitting and subtracting a first-degree polynomial from the data for each trial, in each channel, for each participant. Linear detrending was applied to concentration change data on a trial-by-trial basis, with each trial consisting of 2 seconds of recorded data pre-stimulus onset (baseline) and 18 seconds post-stimulus onset.

Sample size, data retention, reasons for exclusion/withdrawal, and descriptive statistics for retained fNIRS data are provided in Supplementary Materials 4.

### 2.4 Detecting changes in the evoked hemodynamic response

A previous analysis of this dataset found evidence for continued habituation of the hemodynamic response even after the novel stimulus was introduced, as well as a lack of clear novelty response [8], [12]. The analysis outlined here aims to provide an approach to distinguish between trajectories of habituation to familiar auditory stimuli as part of the HaND paradigm, by identifying the timing of significant decreases in the associated evoked hemodynamic response. This would enable the identification of differences in habituation trajectories between groups, to investigate potential differences in this early-life cognitive process. To guide the reader, an overview of the approach to identifying these significant decreases is first provided, before a more detailed description of the key methods involved.

The data across the HaND paradigm was modeled as a functional time series of length 25, the number of trials in the experimental paradigm itself. The identification involved alternate application of the same two steps within a recursive framework – known as ‘wild binary segmentation’ – which enabled the detection of multiple, potentially closely-spaced functional change points within each functional time series. At each hierarchical step/level of wild binary segmentation, FCPt detection proceeded in two stages: first searching for potential functional change points using the FF approach, then testing their statistical significance.

FF FCPt detection is designed to detect a single FCPt [41]. However, in the context of the HaND paradigm, the aim was to identify potentially multiple structural changes in the hemodynamic response trajectory across repeated trials, corresponding to points in the experimental sequence at which the evoked response significantly altered in shape or magnitude. FCPt detection was therefore conducted within the wider recursive framework of wild binary segmentation to permit the identification of multiple candidate change points. This approach splits a functional time series into two sub-series either side of a statistically significant FCPt, when detected. The process is repeated at the next level, with these two subsequent sub-series split once again upon the detection of another significant FCPt, until no more significant FCPts are detected in any remaining sub-series (see Figure 3a).

**Figure 3:**
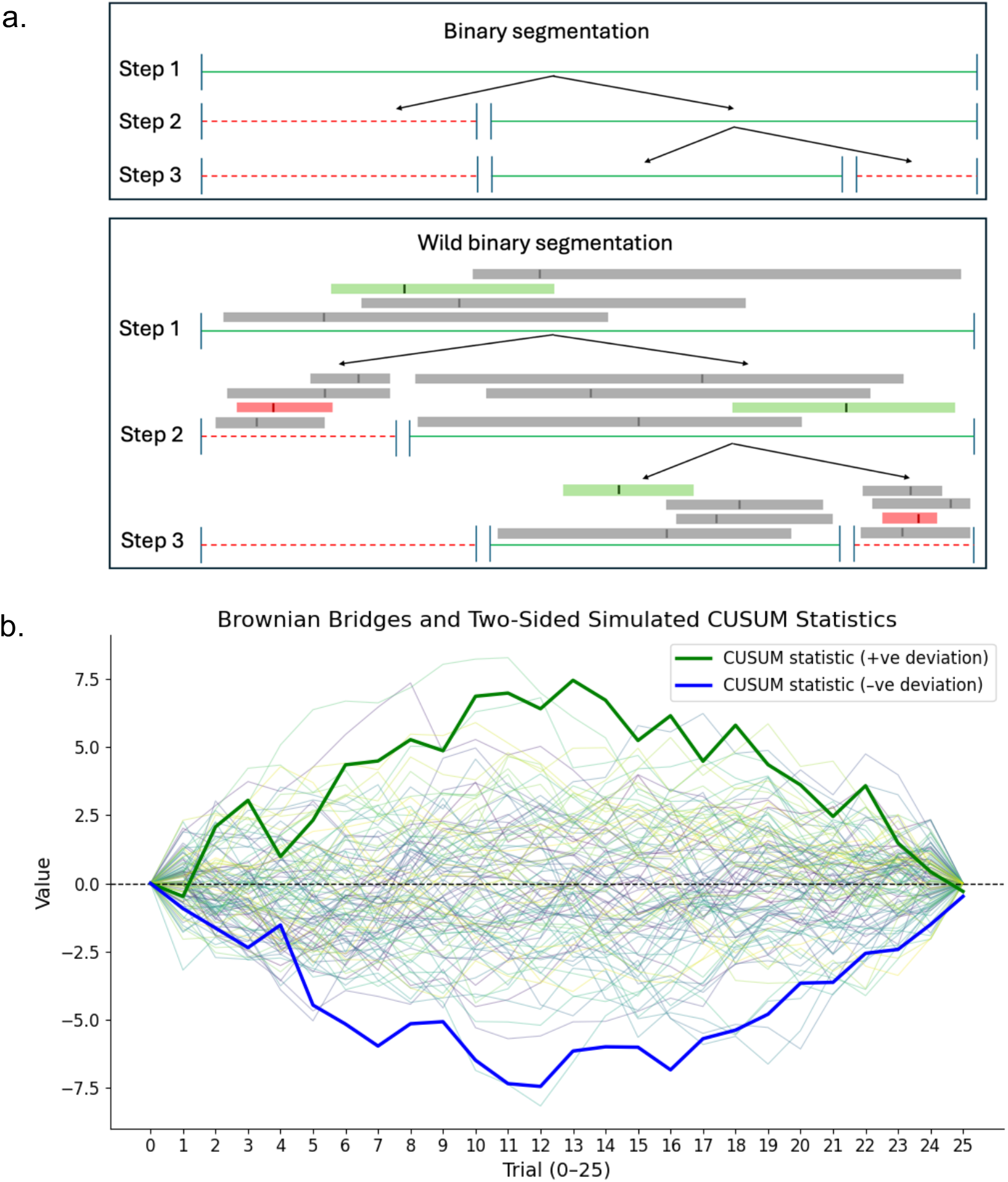
Binary segmentation, wild binary segmentation, and Brownian Bridges. (a) Top: Binary segmentation. Green horizontal lines at each step represent segments with significant changepoints; red dashed lines are segments with non-significant FCPts. Black vertical lines are used to mark trials where structural change occurs. FCPt detection is conducted on the entire range of trials (step 1) or ranges of trials between detected change points (step 2+). Bottom: Wild binary segmentation. In addition to the green, solid lines and red, dashed lines demonstrating segments with/without significant FCPts, respectively, sub-samples of the data used to robustly find the change point with the highest overall test statistic are shown. Grey/green/red bars represent sampled sub-segments, with potential change points marked. Green bars represent the sub-segment of data containing the point with the highest test statistic value. Red bars represent the sub-section with the highest test statistic value (marked) which did not meet statistical significance. (b) 100 simulated weighted sums of Brownian Bridges, anchored at the start and end points, with 25 steps. The maximal value of each generated summed bridge were recorded to create a ‘null’ distribution to compare against the maximal partial sum values across the 25 HaND trials. In this instance, the deviations of the positive (green) and negative (blue) CUSUM statistics would likely be significant, given the absolute magnitude of their largest values (circled).

At each step of wild binary segmentation, candidate change points were identified using the FF FCPt approach and the statistical significance of each candidate change point was then assessed. Multiple comparison correction was required to account for the potential multiple FCPts detected at each level, therefore the calculation of precise *p*-values for each FCPt was necessary. Monte Carlo simulation was employed to this end, by comparing observed test statistics to those derived from a generated set of Brownian Bridge processes.

The final set of FCPts across each channel’s functional time series is the collection of FCPts, identified at all levels of wild binary segmentation, which were significant after the appropriate multiple comparison correction had been conducted. Unless otherwise stated, functional change point detection and associated statistical significance testing was implemented in Python 3.9.16 [42] using in-house, publicly-available code.

#### 2.4.1 Functional change point detection

The aim of FCPt detection is to identify the largest structural breaks in a functional time series (see Figure 1) which are then subsequently tested for statistical significance. As applied to data collected using the HaND paradigm, this approach was used to identify trials within the paradigm where a potentially significant hemodynamic response change in amplitude and/or shape occurred. Full mathematical detail can be found in Appendix A; an intuitive overview is given here.

Each channel/chromophore combination contains a functional time series of 25 functions corresponding to the 25 trials in the HaND paradigm. For each of these channel/chromophore combinations, potential change points for each functional time series were identified via FF and FPCA-based FCPt detection, however, the FF approach yielded more consistent and interpretable results than the FPCA-based approach, with little difference in computational time. Accordingly, the FF approach and subsequent findings yielded by its implementation are outlined in the main body of this work; information on the use of FPCA-based FCPt can be found in Supplementary Materials 2, 3, and 5.

FF FCPt detection [25] uses a test statistic, 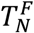, which, in intuitive terms, measures how much the average hemodynamic response across the first n trials differs from the overall average across all trials [43, Ch. 6], [44]. These so-called cumulative differences fluctuate randomly around zero in the case of a stable underlying hemodynamic response or when no consistent change is present in the data. Under the alternative, when the shape or magnitude of the mean response changes – such as when the hemodynamic response declines with habituation – these cumulative differences grow sharply due to deviation within each trial. The trial number at which this divergence reaches its largest value (i.e. the largest value of the test statistic) marks the most likely location of a structural change in the data and is a potential FCPt, which is then tested for statistical significance. In the case of two trials producing the same maximal test-statistic, the one which occurs first is chosen, i.e. the one corresponding to a smaller trial number in the experimental paradigm. In practice, this “cumulative-sum” (CUSUM) approach captures the trial at which the signal’s average behavior shifts most strongly over time.

#### 2.4.2 Wild binary segmentation for multiple change point detection

The FF FCPt detection approach and 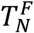 are designed to detect a single FCPt [41]. Wild binary segmentation was used to enable the detection of multiple, (potentially) closely-spaced candidate change points within a time series of functionally modeled hemodynamic responses.

Ordinary binary segmentation splits data at points where potential changes are detected, recursively partitioning the time series into subintervals and reapplying the detection procedure within each segment until no further significant change points are found (see Figure 3a, top). This is efficient but can overlook closely spaced change points as dominant changes may mask nearby ‘smaller’ ones [45], [46]. Wild binary segmentation [46] addresses this by generating multiple subintervals of different lengths in each data segment; detecting candidate change points in each; and recording the largest test statistic 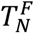. The candidate change point *n*^∗^ for the entire current data interval corresponds to the location of the largest test statistic across all subintervals (see Figure 3a, bottom).

#### 2.4.3 Monte Carlo simulation to assess functional change point significance

Multiple comparison correction was applied at each hierarchical level of wild binary segmentation to control the family-wise rate and ensure robustness of the findings. Monte Carlo simulation was therefore conducted to generate precise *p*-values and enable this correction. For full Mathematical detail of both the Monte Carlo simulation and multiple comparison correction, see Appendix B.

As the number of trials in the experimental paradigm, *N*, increases, the test statistic under the null hypothesis of no significant FCPts is capped by the largest value in a weighted sum of random fluctuations which are tethered to zero at each end, known as Brownian Bridges (see Figure 3b), with the weights corresponding to different components of data variability [22], [47].

To permit multiple comparison corrections it was necessary to calculate significance values for each potential FCPt. Statistical significance testing of potential change points was therefore conducted by comparing their associated test statistics to the maximum values of 10 000 generated summed Brownian Bridges of length *N*_*b*_, where *N*_*b*_ is the length of the functional time series interval being assessed at each stage of wild binary segmentation. The statistical significance of the potential FCPts’ associated test statistic was then given by the proportion of larger randomly generated maximum values.

Generating these significance values then permitted within-channel multiple comparison correction using the Bonferroni-Holm method [48] to correct the family-wise error rate (FWER), which was increased by conducting separate statistical tests for each potential FCPt within a channel. To address the hierarchical nature of the wild binary segmentation process, in which change points’ significance at ‘lower’ levels should only be tested if those at previous, ‘higher’ levels are statistically significant (see Figure 3a), a step-down multiple comparison procedure was conducted within each channel at every binary search level (full detail in Appendix B (ii)). The Bonferroni-Holm method was applied at each level using the *statsmodels* Python package [48], [49]. An alpha value of 0.01 was used to determine the statistical significance of change points.

### 2.5 Further analysis of statistically significant change points

As a first step, visual and qualitative assessment of the data before and after statistically significant FCPts was conducted, to assess the extent to which the detected FCPts corresponded to physiologically meaningful changes in the evoked hemodynamic response. To further verify the sensitivity of FCPt detection to the consistent evoked hemodynamic response structure, statistical testing was also performed to compare the channels containing FCPts with those in which statistically significant hemodynamic responses were identified. Further methodological details of this quantitative analysis can be found in Supplementary Materials 5.

Beyond these validation steps, it was important to determine the extent to which the detected changes captured a decrease in the hemodynamic response alongside the repeated stimuli. Furthermore, in line with Objective (b) of this work, statistical assessment of the longitudinal change in the timing of detected FCPts across the 5-, 8- and 12mo ages was conducted. These latter two analyses are described below.

#### 2.5.1 Assessing ‘types’ of change points

The motivation for using FCPt detection is to better understand the timing of changes in the hemodynamic response during the course of the experimental fNIRS session and, subsequently, assess the presence of habituation in the evoked response to a stimulus. Whilst this timing is valuable information, it is also important to understand the nature or type of structural change which occurs in the mean hemodynamic response – whether it corresponds to an increase or decrease in magnitude – since changes associated with habituation would likely be represented by decreasing FCPts. Accordingly, to determine whether the mean hemodynamic response increased or decreased following change points, the mean hemodynamic response was calculated before and after each statistically significant FCPt, then Simpson’s rule was used to estimate the definite integral of the function’s absolute values, using *scipy*.*integrate*.*simpson* [50]. The definite integral estimate after each change point was subtracted from that before it, as an estimate of area-under-the-curve (AUC) difference between functions; a positive value indicated a decrease in response. The proportion and statistical significance of the number of decreasing change points – suggesting habituation – was assessed using two-sided binomial tests, comparing the observed proportion of decreasing change points against a null hypothesis of random directionality. Analyses were conducted on all detected FCPts, as well as the first- and last-occurring FCPts in each channel in order to ascertain whether FCPt detection was sensitive to the start and end of the habituation process, respectively. Tests were conducted by age group, plus a combined set of FCPts across all three age groups of 5-, 8- and 12mo, with Bonferroni-Holm correction for multiple comparisons to control the FWER in all cases [48].

#### 2.5.2 Analyzing the effect of age

A key motivation for obtaining information regarding the timing of significant changes in the hemodynamic response, was the ability to distinguish between different groups’ habituation trajectories. Indeed, findings at 5-, 8- and 12mo from infants sampled for the dataset used in this work have previously found evidence for continued habituation even into the ‘novelty’ trials and additionally provided valuable evidence for spatial variation in habituation across ages [8], [12], yet the trial-specific timing of this habituation was not directly quantified.

The effect of age group on the rate of habituation was therefore examined in this longitudinal dataset by comparing the distributions, across the 5-, 8- and 12mo age groups and chromophores, of the trial numbers of the following sets of FCPts:

i. Decreasing change points
ii. Decreasing change points which were ***also*** the first to occur in a channel (‘First And Decreasing’)
iii. Decreasing change points which were ***also*** the last to occur in a channel (‘Final And Decreasing’)
iv. The earliest occurring decreasing change points in a channel (‘Earliest Decreasing’)
v. The latest occurring decreasing change points in a channel (‘Latest Decreasing’)

Set (i) was generated to investigate the timing of FCPts which likely correspond to habituation. Sets (ii) and (iii) investigate when habituation begins and ends, respectively. Sets (iv) and (v) were generated to account for decreasing FCPts in the hemodynamic response after or before an increase in the hemodynamic response, respectively, and which would therefore be discounted from sets (ii) and (iii).

Weighted ordinal logistic regression (OLR) was used to assess inter-age differences between trial numbers for each distribution (i)-(v), supplemented with post-hoc testing and multiple comparison corrections. OLR [51] preserves the ordinal nature of the outcome (FCPt Trial Number); and was conducted using the *OrderedModel* from the *statsmodels* package [49]. The predictors ‘Age’ and ‘Change Point Count Per Channel’ (the sum of age/chromophore combinations with identified change points, per channel, ranging from 0 to 6) were used to investigate associations with age and control for Channel differences. Weights were applied based on the number of participants whose data was used to construct each mean functional observation in question, to account for unequal numbers of valid trials across participants contributing to each functional observation. The model was fit using the logit distribution and optimized using the Broyden–Fletcher–Goldfarb–Shannon algorithm.

A post-hoc Wald test [52] confirmed any significant impacts of age on change point outcomes. Pairwise differences between age groups driving significant age effects were identified using an ANOVA model followed by Tukey’s Honestly Significant Difference (HSD) [53] to control the FWER.

## 3 Results

At all three ages, both HbO and HbR yielded at least 10 channels with statistically significant FCPts amongst the 25 paradigm trials (Table S1). Six channels at each age demonstrated significant FCPts in both chromophores, although these were not identical across ages. These channels were primarily localized to the superior and middle temporal gyri, as identified previously using channel registration [29]. Two channels exhibited significant change points in both chromophores at all ages, located bilaterally in homologous temporal regions (see Figure 4). Channels with FCPts in both chromophores typically exhibited canonical hemodynamic response morphology (Figure 5 and Figure 6), suggesting that FCPt detection was sensitive to task-evoked response structure.

**Figure 4:**
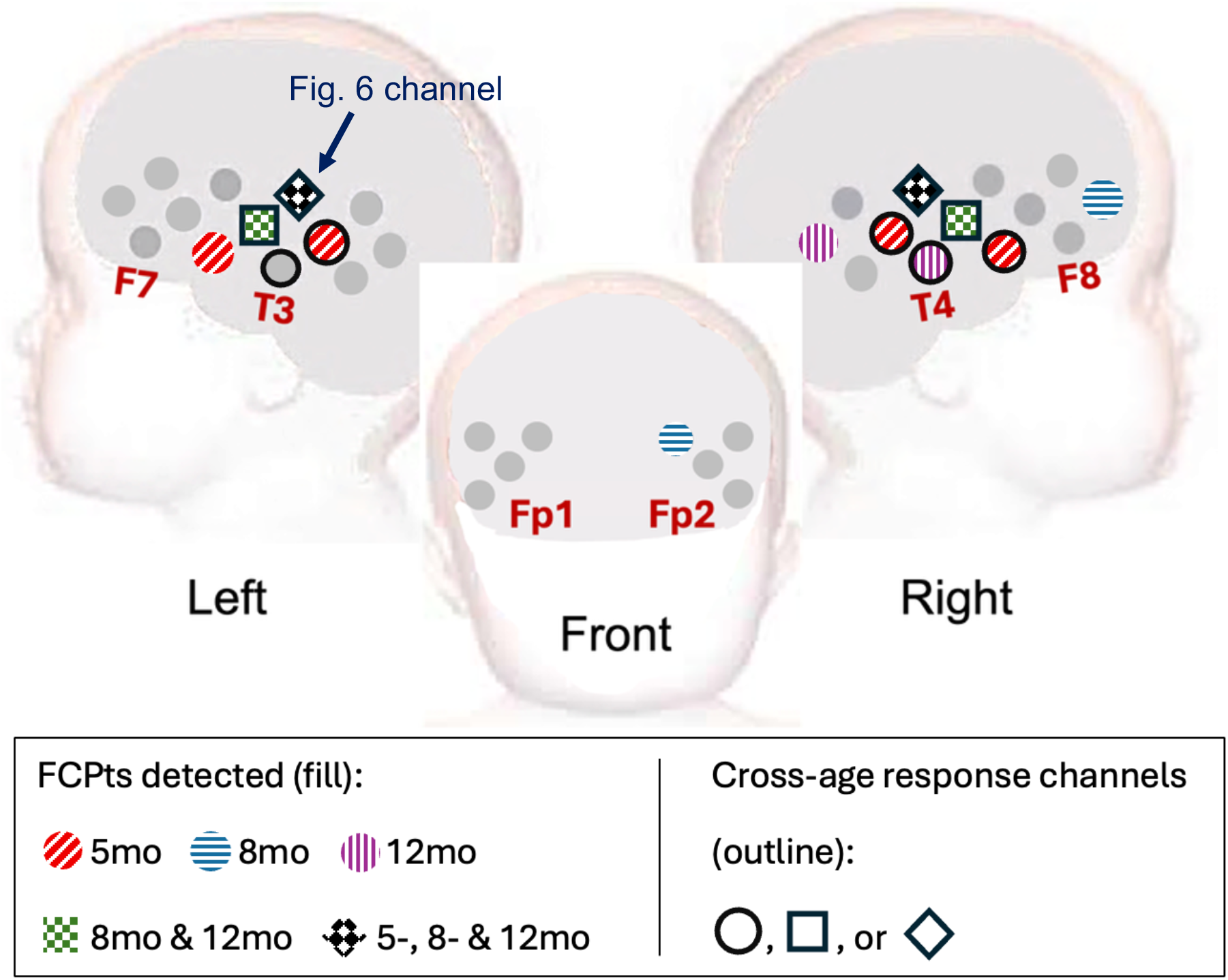
location of channels with detected FCPts in both chromophores for each of the 5-, 8- and 12mo cross-infant data groupings. Channels were typically located in auditory-associative temporal regions, with two of these channels – located in the same position in each hemisphere – containing FCPts in both chromophores at all three ages. A further two channels which, again, were located in the same position in both hemispheres, contained FCPts in both chromophores for the 8- and 12mo age groups. Channels for 5mo infants were less frequently shared by the other age groups, however all six channels at this age were in auditory-associative channels, in contrast to the two older age groups for whom the localization was more varied. The channel containing data in displayed in Figure 6 is indicated by an arrow. Cross-age channels with significant responses in either chromophore, as identified by conventional analyses approaches, are outlined as they are used during further analyses.

**Figure 5:**
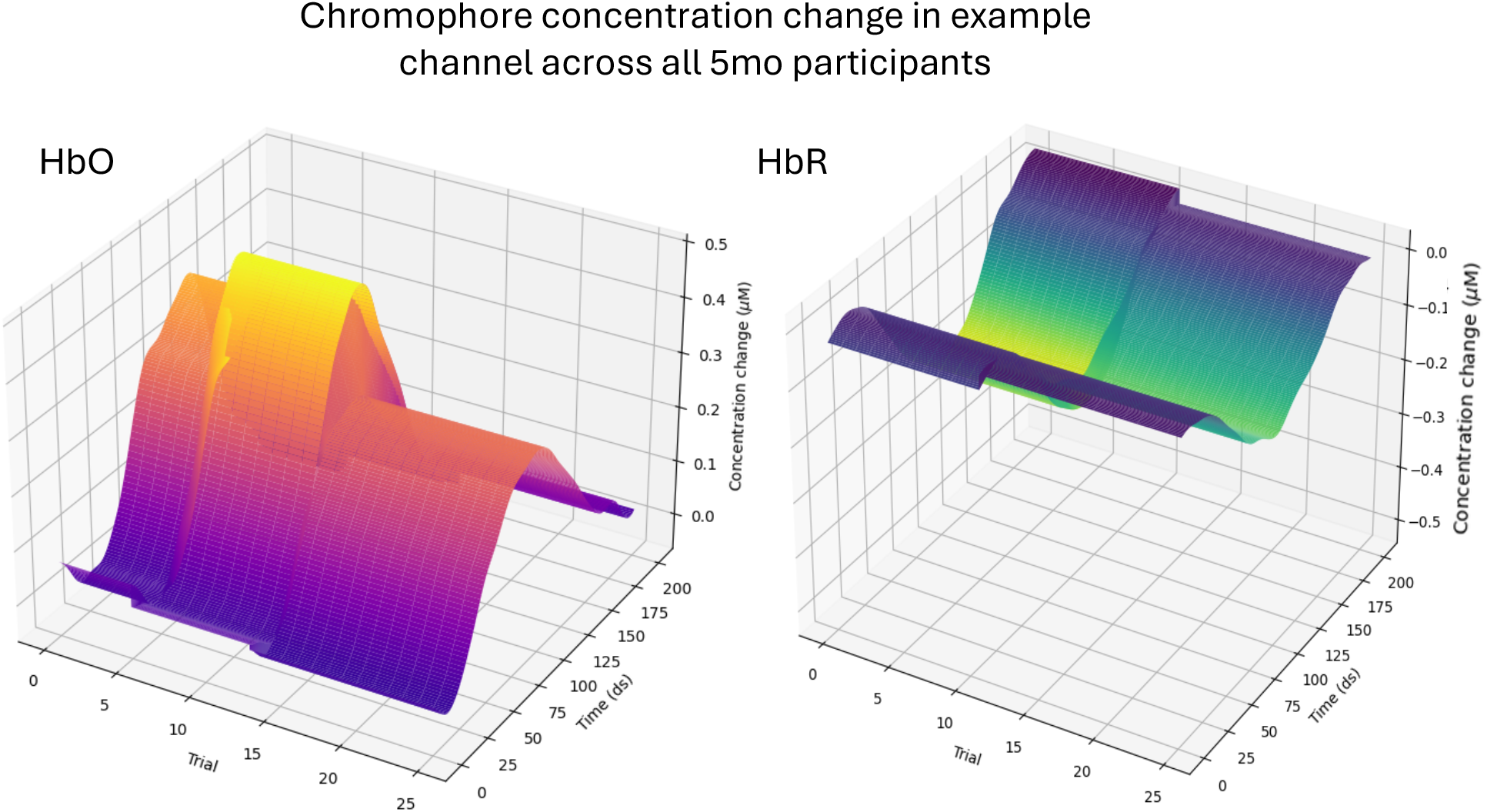
Change points and mean surface between them for a cross-age ROI channel using 5mo data. The channel shows clear hemodynamic response structure and habituation profile in both chromophores. It also exhibits an early apparent increase in the HbO response.

**Figure 6:**
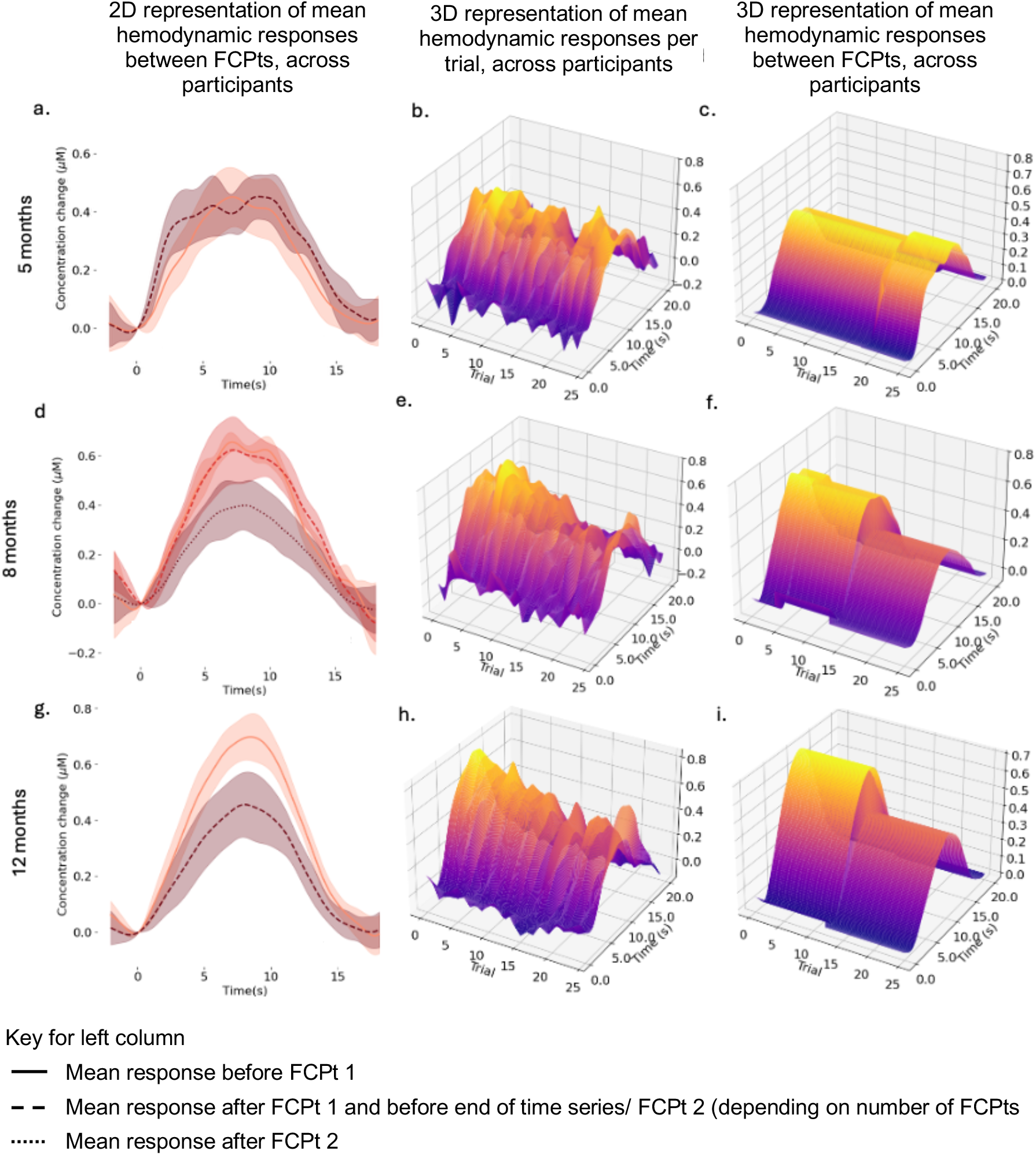
Hemodynamic responses, and their mean representations between FCPts, in a single example channel. Channel location is indicated in Figure 4. Left column: mean responses across infants before/after detected FCPts. Middle column: 3D surface representation of the trial-by-trial data across infants. Right column: 3D surface representation of the mean responses across infants before/after detected FCPts. This channel demonstrates a clear hemodynamic response structure at all three ages (in both chromophores – see Supplementary Materials 7). Univariate analyses did not detect a significant hemodynamic at 5mo. Figure (c) illustrates the capacity to distinguish between hemodynamic responses based on structural information rather than solely the magnitude of the sampled fNIRS values. Figures (a), (d) and (g) illustrate an increasing difference pre- and post-FCPt with age.

To assess spatial plausibility quantitatively, overlap was first assessed between (i) channels containing significant FCPts at each age in either chromophore and (ii) a cross-age set of channels defined as those exhibiting a significant evoked hemodynamic response at all of the three ages in prior block-averaged analyses, in either chromophore [12] (see Figure 4). This cross-age set of channels reflects those plausibly responsive to the auditory paradigm across development, without requiring age-specific detection at each time point. Statistically significant overlap was observed, indicating that FCPt detection was sensitive to the morphology and changes of time-locked, task-evoked responses (Supplementary Materials 5a). As a less spatially restrictive but age-matched validation, overlap was also examined at each age separately, comparing channels containing significant FCPts with those exhibiting significant evoked hemodynamic responses within that age (Supplementary Materials 5b), matched by chromophore. This analysis yielded consistent results, indicating that FCPt detection remained preferentially localized to task-relevant regions under more specific spatial criteria.

Whilst this overlap is encouraging, it is important to emphasise that the intended use of FCPt detection is to complement conventional (block-averaging based) analyses by providing additional insight into the temporal characteristics of habituation. Therefore, to assess the extent to which detected FCPts corresponded to physiologically meaningful changes in the evoked hemodynamic response, visual and qualitative assessment of the data before and after statistically significant FCPts was also carried out.

FDA representations of the evoked hemodynamic responses were faithful to their discrete counterparts, preserving the canonical hemodynamic shape while providing a smooth functional representation. Within this framework, FCPt detection identified statistically significant structural breaks in the mean response. This enabled representations of consistent “before” and “after” response forms within individual channels in both chromophores (Figure 6 and Figure 4).

In many channels, the change corresponded to a clear alteration in response magnitude, visible in the mean functions before and after the detected FCPt (Figure 6a, d, g). However, the method additionally provided the trial number at which this structural shift occurred, enabling examination of when habituation-related changes emerged across the experimental session (Figure 6c, f, i).

Trial numbers of significant FCPts ranged from 3 to 20, indicating substantial variation in the timing of structural changes across channels and ages. This variability highlights the limitation of block-averaging approaches, which implicitly assume stable responses within predefined trial blocks and cannot identify discrete transition points in the response trajectory. It also therefore reinforces the need for approaches with the capacity to assess habituation tasks on a trial-by-trial basis.

Crucially, FCPt detection identified the specific trial at which significant structural change emerged within the sequence of repeated stimulus presentations. This trial-specific temporal information provides a level of specificity unavailable to block-averaged, magnitude-based summaries alone.

### 3.1.1 Assessment of change point ‘types’

The proportion of ‘decreasing’ change points was statistically analyzed using two-sided binomial tests, to assess the sensitivity of FCPt detection to decreases in the hemodynamic response as a measure of habituation. Results (Table 1) are reported for channels previously identified as task-responsive across all three ages (5-, 8-, and 12-months) using conventional block-averaged methods (‘cross-age channels’) [12]. This approach balances restriction of analyses to plausible task-responsive regions while accounting for age-related differences in response strength or detectability. Analyses on age- and chromophore-specific response channels were also conducted; results were consistent and can be found in Supplementary Materials 6.

**Table 1:** Proportion of decreasing FCPts which occurred first, last, or anywhere during the channels’ time functional series.

| Age | Change points | Total | Decreasing | Proportion (%) | *p*-value (adj.) |
| --- | --- | --- | --- | --- | --- |
| 5mo | ALL | 13 | 8 | 61.5 | 0.581 |
|  | First | 6 | 4 | 66.7 | 0.688 |
|  | Last | 6 | 2 | 33.3 | 0.688 |
| 8mo | ALL | 16 | 16 | 100 | <0.001*** |
|  | First | 8 | 8 | 100 | 0.013* |
|  | Last | 8 | 8 | 100 | 0.013* |
| 12mo | ALL | 14 | 13 | 92.3 | 0.005** |
|  | First | 7 | 6 | 85.7 | 0.167 |
|  | Last | 7 | 7 | 100 | 0.023* |
| All | ALL | 43 | 37 | 86.0 | <0.001*** |
|  | First | 21 | 18 | 85.7 | 0.005** |
|  | Last | 21 | 17 | 80.1 | 0.013* |

Significance reported after Bonferroni-Holm correction ( $p^* < 0.05$ ; $p^{**} < 0.01$ , $p^{***} < 0.001$ ). Proportions for the 5mo data were varied, with no FCPT category containing a significant number of decreasing changes. At 8mo all detected FCPTs were decreasing, with all but one FCPT decreasing at 12mo. Results at these latter two ages likely drove the results when analyzing across all three ages, in which the first, last, and overall set of FCPTs all had a significant proportion of decreasing FCPTs.

There were no statistically significant differences in the proportion of decreasing change points at 5mo across all categories of change points (*p*-values > 0.5). At 8mo, in contrast, all mean hemodynamic response functions decreased after detected FCPts, so taking these FCPts together and examining both first and last FCPts produced significant proportions (*p* < 0.01, *p* < 0.05, and *p* < 0.05 respectively) in all three cases. All but one of the first occurring FCPts detected across at 12mo corresponded to a decrease in the mean hemodynamic function, though the proportion of decreasing FCPts in this category was not significant (*p* > 0.05), likely due to small sample size. However, all other detected FCPts at this age were decreasing ones, meaning the proportions of final (*p* < 0.05) and overall (*p* < 0.01) decreasing FCPts were significant.

Combining change points across all ages, the proportion of all detected change points, first change points, and last change points which decreased were all significantly higher than expected by chance (*p*-values < 0.001, *p* = 0.006, and *p* = 0.017, respectively). This is likely driven by the results from 8- and 12mo infants, considering the age-specific results reported above.

### 3.1.2 Effect of Age

Investigation of longitudinal change in the timing of habituation was conducted using weighted ordinal logistic regression, to examine the effect of infant age on the timing of decreases in the hemodynamic response and therefore investigate longitudinal change in habituation trajectories. As with the analyses of decreasing change points, results are reported for FCPts detected within the cross-age task-responsive channels defined above. Analyses on age- and chromophore-specific response channels were also conducted; results were consistent and can be found in Supplementary Materials 7.

Results are reported fully in Table 2 and Table 3. The Age coefficient was significant for the Decreasing (*β* = −0.247, *SE* = 0.114, *z* = −2.17, *p* = 0.030), Final And Decreasing (*β* = −0.497, *SE* = 0.218, *z* = −2.28, *p* = 0.022), and Latest Decreasing change points (*β* = −0.359, *SE* = 0.175, *z* = −2.06, *p* = 0.040). Wald tests confirmed the significance of these differences (see Table 3).

**Table 2:** Weighted ordinal logistic regression results for FCPts.

| Change points | $\beta$ | S.E. | $z$ | $p$ (adj.) |
| --- | --- | --- | --- | --- |
| Decreasing | **-0.247** | **0.114** | **-2.17** | **0.030\*** |
| First And Decreasing | -0.216 | 0.165 | -1.30 | 0.192 |
| Final And decreasing | **-0.497** | **0.218** | **-2.28** | **0.022\*** |
| Earliest Decreasing | -0.195 | 0.145 | -1.34 | 0.180 |
| Latest Decreasing | **-0.359** | **0.175** | **-2.06** | **0.040\*** |

*Detected across both chromophores, after Bonferroni-Holm correction. Significant effects for age were found in the change points which decreased, particularly those occurring later in the series of 25 trials. Significance: $p^* < 0.05$ ; $p^{**} < 0.01$ , $p^{***} < 0.00$ .*

**Table 3:**
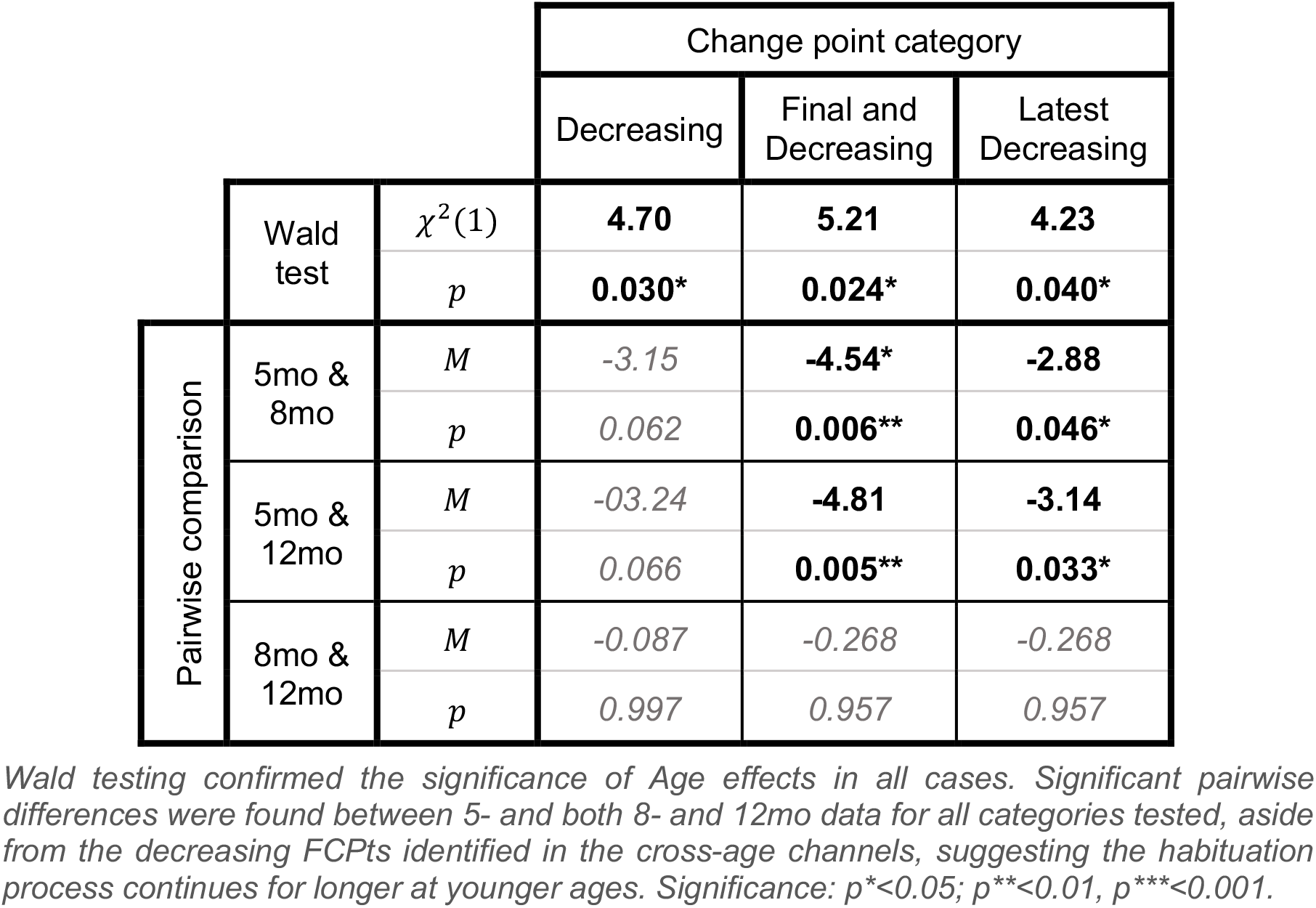
Post-hoc testing of change point categories for which significant channel effects were identified using weighted ordinal logistic regression.

In all three cases, Tukey’s HSD results showed that differences between age groups were driven by later change point occurrences at 5mo than the two later time points. Decreasing change points overall occurred a mean of 3.58 trials later than at 8mo (*p* = 0.026) and 3.66 trials later than at 12mo (*p* = 0.029). More specific analysis of late-occurring, decreasing change points showed that change points which were both Final And Decreasing had mean trial occurrences 4.54 later at 5mo when compared to 8mo (*p* = 0.006) and 4.81 trials later when compared to 12mo (*p* = 0.005). Similarly, the Latest Decreasing change points occurred an average of 2.88 trials later at 5mo than at 8mo (*p* = 0.046) and 3.14 trials later than at 12mo (*p* = 0.033). Interestingly, the earliest occurring FCPts were relatively consistent across age groups, with no other significant results found. No significant pairwise effects were found for Decreasing change points. No significant effects of Change Point Count Per Channel were found for any of the change point types.

These age-related differences across ages in the later change point occurrences are best exemplified by the differences between the Latest Decreasing change points (see Figure 7; similar figures for other change point types can be found in Supplementary Materials 7), which shows both the smaller mean trial number for the two older age groups and the greater distribution breadth of responses found in younger infants.

**Figure 7:**
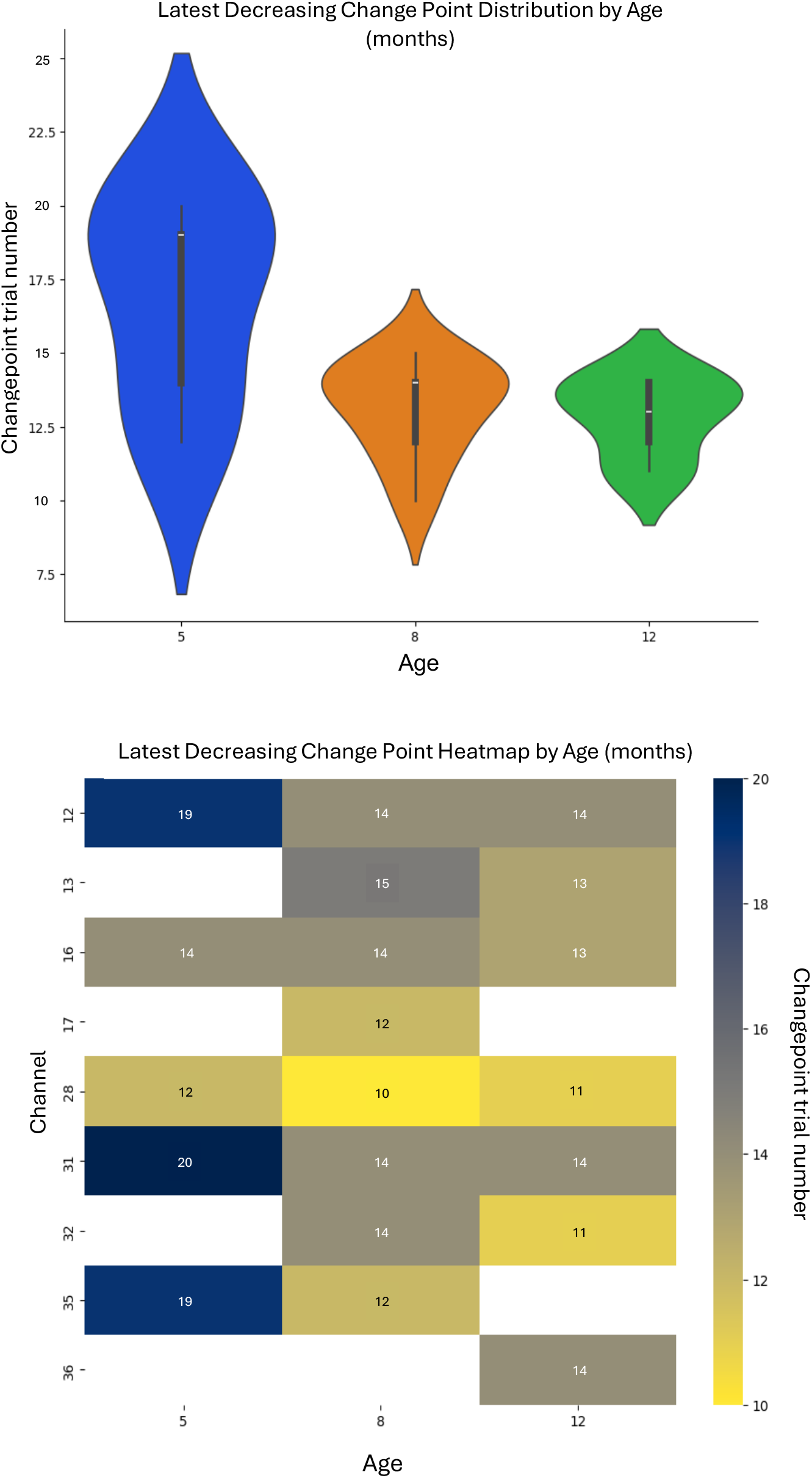
Age effects for latest decreasing FCPts across both chromophores. Top: Violin plots illustrating the difference in distribution of latest decreasing change points at 5-, 8- and 12mo. Distribution differences can be observed between 5- and both 8- and 12mo data, with similar distributions for the older two ages. Bottom: precise trial numbers of the latest occurring decreasing FCPts in each channel.

Visualization of other aspects of habituation are noted when visualizing the change in mean hemodynamic response AUC across channels for the 25 HaND paradigm trials, calculated using Simpson’s rule for integration and presented as a proportion of the first trial (Figure 8). The figure illustrates a greater proportional magnitude of decrease in size for the two older age groups, as well as the trajectory of decrease in which a sharp, step-like drop occurs after a stable initial phase and before stabilising again at a steady lower level after around 15 trials. These phenomena are exhibited to a lesser extent at 5mo, though the decrease in change is smaller and reaches a more unstable plateau towards the end of the series of paradigm trials, pointing to habituation which is smaller in size and less consistent at this age.

**Figure 8:**
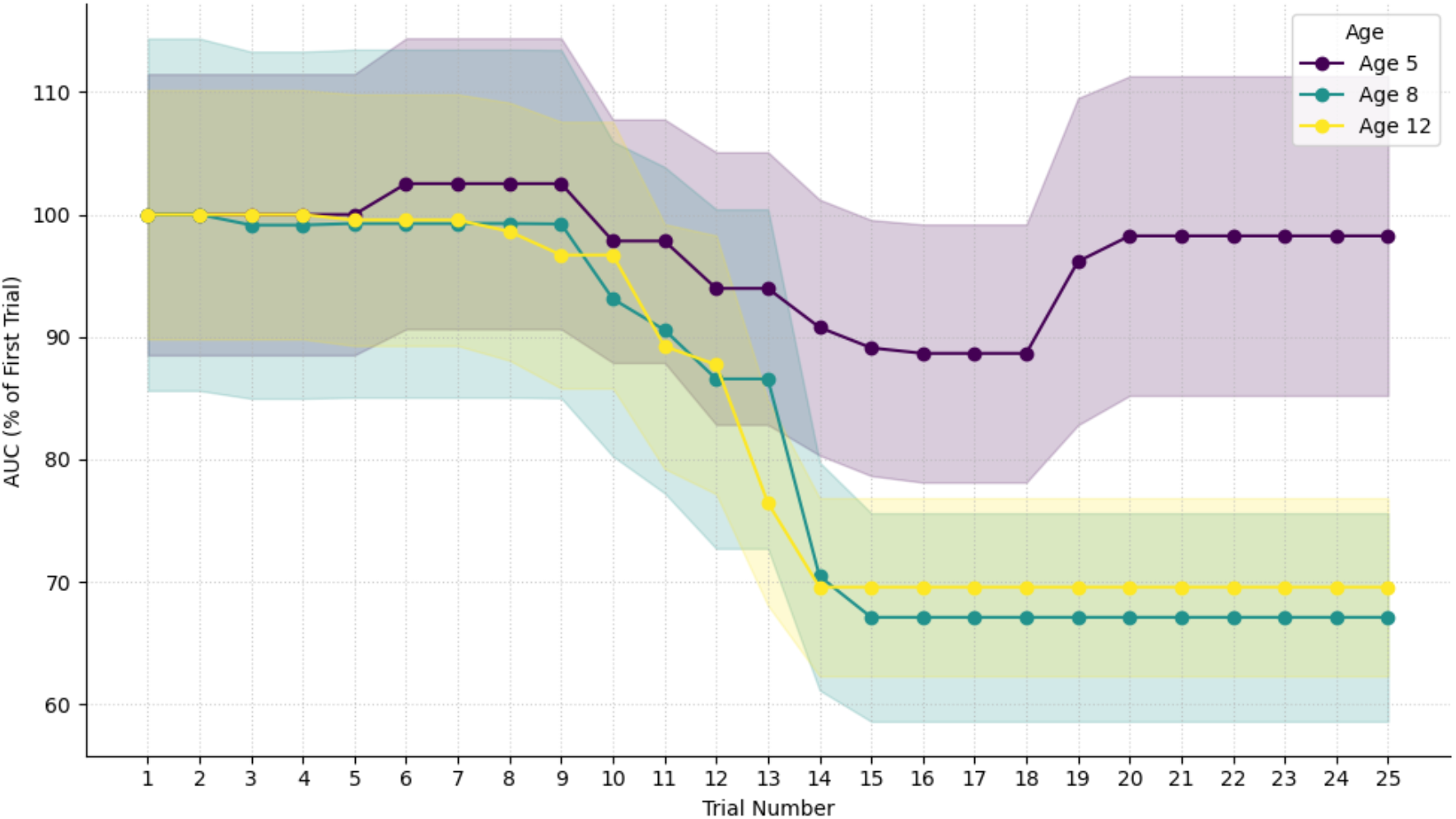
Mean area under the hemodynamic response function across channels with FCPts, expressed as a percentage of the 1st of the 25 HaND trials. Area-under-the-curve (AUC) was calculated in each ‘cross-age’ channel exhibiting significant hemodynamic responses, as identified using conventional analyses. Line graphs plot the mean calculation of this AUC across channels for each age. Greater decreases which reach a more consistent and ‘settled’ plateau can be observed at 8- and 12mo, when compared to those at 5mo. Shaded regions represent ±1SEM. Trial-by-trial investigation of the data in this manner is a key advantage of using FDA, and FCPt detection more specifically, to examine the changing evoked hemodynamic response function in during habituation.

## 4 Discussion

### 4.1 Assessing the timing of habituation

The present findings support the view that FCPt detection provides insight into trial-specific timing information of fNIRS task-based infant data unavailable to conventional, magnitude-based approaches. Specifically, FCPt detection using a fully functional approach was able to characterize change in the infant hemodynamic response, evoked by repeated auditory stimuli, and measured using fNIRS.

FCPt detection appears to accurately capture structure in fNIRS signals corresponding to the evoked hemodynamic response, indicating sensitivity to the consistent morphology of time-locked evoked responses. Examination of modeled mean hemodynamic responses before and after detected FCPts implied that the change points themselves were qualitatively representative of change in the in the evoked hemodynamic response (Figure 5 and Figure 6, and Supplementary Materials 8). Moreover, the significant proportion of decreasing change points – identified in a dataset in which evidence for significant habituation without novelty response already exists at these ages [8], [12] – points to the sensitivity of FCPt detection to changes in the hemodynamic response specifically due to habituation. Further, this is consistent with other studies which have previously reported repetition suppression in response to repeated stimuli in developmental populations [2], [54].

FCPt detection revealed that these significant changes occurred at a range of trial indices covering over 70% of the experimental paradigm, with FCPts detected at all of the trial numbers within this range. This underscores the importance of trial-wise analytical approaches capable of detecting variability in habituation timing. It also provides evidence for a gradual decrease in response during habituation [3], whilst the variation of change point trial numbers across channels also suggests that the timing of habituation may be dependent on cortical location.

Additionally, these change points were detected using information across the entire temporal window of the evoked hemodynamic response by modeling each hemodynamic response as a holistic statistical object. This approach is possible since modeling of functions within FDA exploits the smooth behavior of functional objects and naturally incorporates the temporal structure of functions [19], [20]. Nevertheless, analyses outlined here point to the possibility and advantages of considering evoked hemodynamic responses holistically, rather than relying on peak amplitude or predefined time windows.

Together, these findings indicate that FCPt detection provides characterization of the timing of structural change in the hemodynamic response in a manner not achievable using block-averaged summaries.

### 4.2 Developmental change in habituation

This characterization of timing enabled assessment of longitudinal changes in evoked infant habituation responses, demonstrating that older infants complete habituation earlier within the experimental sequence.

Development of habituation in the evoked hemodynamic response is evident in the assessment of age effects on the timing of various types of change points. The later decreasing changes in the hemodynamic response typically occur earlier at 8- and 12mo than at 5mo, where the FCPt timing distribution is more varied. This suggests that infants’ responses and associated hemodynamic responses may attenuate more quickly to repeated spoken stimuli at older ages, despite appearing to begin at approximately the same time (see Figure 7). The lack of significant differences in timing of FCPts between 8- and 12mo does, however, suggest that this aspect of habituation development is fully realised by 8mo.

Whilst the proportion of decreasing FCPts detected at 8- and 12mo was marked, results for 5mo infants were more ambiguous. No statistical significance was found at this age for the proportion of decreasing changes when considering the first, last, or overall set of detected change points. This is consistent with previous analysis using conventional methods [8], [12]. In part, this may be driven by some channels which exhibited early increases in hemodynamic response (see Figure 5), possibly reflecting an initial increase in responsiveness (‘sensitisation’) before habituation [55]. Conversely, some channels contained increasing change points towards the end of the experimental paradigm (see Figure 8). Given that the peak of mean hemodynamic responses before and after structural changes in several of these channels is similar in magnitude, it is possible that the higher percentage of increasing FCPts reflects detection of structural change which is overlooked by analysis using summary scalar values (see Figure 6c). However, this wouldn’t explain why these types of detected changes seem to be unique to the 5mo data beyond the possibility that the data collected at younger ages contains more noise due to motion and/or poor scalp-optode coupling [56]. Alternatively, it is possible that the variation in the nature of structural changes at this age corresponds to the ongoing development of important auditory and speech perception processes [57], [58]; anomalous results caused by variation due to a smaller (channel-wise) sample size; or possibly even a true novelty response, or else a combination of these factors.

### 4.3 Complementary roles of FCPt detection and conventional approaches

FCPt detection was sensitive to trial-wise structural changes in the hemodynamic response, capturing nuanced temporal information about variation in magnitude. Conversely, outside of auditory-associative channels, FCPts were detected in response to changes in signal properties not clearly attributable to the evoked hemodynamic response, such as a greater prevalence of higher frequency data components or temporally-localized changes in shape rather than magnitude. These detected FCPts possibly corresponded to physiological phenomena unrelated to the evoked hemodynamic response and to persistent motion artifacts, respectively, and would be mitigated by the averaging across different paradigm trials which is central to conventional analysis approaches. In short, both FCPt detection and conventional analyses methods ask and answer fundamentally different questions. Used in combination with conventional analyses (e.g. block averaging, GLM-based approaches), FCPt detection enables comprehensive, multifaceted investigation of functional NIRS data.

### 4.4 Strengths, limitations, and future avenues for research

Several major strengths in this work are methodological, some of which are introduced by FCPt detection itself and are described above. Aside from these, work presented here benefits from the use of hierarchical wild binary segmentation. Firstly, the inclusion of this method enables the detection of multiple change points – important in habituation paradigms, where changes may unfold gradually rather than as a single discrete shift – whilst countering the potential masking of significant FCPts by other dominant changes located closely in the series of trials [45], [46]. The hierarchical testing framework also ensured statistical control of the FWER while allowing identification of multiple change points.

Another key strength is the interpretability of the results. Mean functions before and after FCPts are readily understandable and clear when presented in two dimensions (see Figure 6a, d, and g). When used to construct three dimensional representations they are faithful enough to the original data that correspondence to it can be observed, yet still clearly indicate the timing of significant change within the functional time series (Figure 6b, c, e, f, h, and i). Moreover, whilst change point detection can be conducted on scalar data [59], [60], [61], [62], modeling each trial as a functional object means that any change points detected are necessarily discrete values and correspond to a particular trial index in the experimental paradigm. This imbues results with simplicity and intuitive meaning, respectively.

Limitations of this work include the use of CUSUM-based statistics, which are biased towards detection in the middle of the observation series. This bias stems from the fact they are designed to accumulate deviations from a reference value over time; as such, near the start/end of the series, there are limited observations with which to establish a baseline or confirm a persistent shift [63]. This bias persists even with wild binary segmentation and is particularly important to consider when interpreting the apparent shape of the habituation trajectory. Alternative approaches altogether may improve FCPt detection at series boundaries and mitigate this bias [64].

Persistent physiological signal components in individual data did not permit meaningful change point detection or subsequent analysis at this level; FCPt detection was therefore conducted on group data. Consequently, small (group-wise) sample sizes for change points at each age increased volatility in results, making them sensitive to minor variations in ROI channels, meaning caution is warranted when interpreting results concerning age-related differences. Future work may improve individual-level robustness by exploiting the greater spatial resolution afforded by HD systems [65], [66], either to analyze spatially averaged individual data e.g. using parcellations [67] or by incorporating larger channel numbers in group analysis.

A further limitation is that the approach used here assumes that the residual functions *ϵ*_*n*_(*t*) are independent, whereas in reality temporal dependence between observations likely exists due to the design of the HaND paradigm. The long-run (LR) covariance function can account for the cumulative effect of temporal dependencies [68] and can be used during FCPt detection [44]. However, it was found that the length (*N* = 25) of the functional datasets was not sufficient to generate reliable LR covariance estimates (see Supplementary Materials 11), limiting detection accuracy. Using the ordinary covariance function introduces potential standard error underestimation and consequent false positive inflation, though a stable yet biased estimate was preferred to a variable LR estimate. Moving forward, it may be interesting to look at the relationship between the number of trials and the stability of the LR covariance function, since the extent to which the error introduced by using the ordinary covariance function is currently unknown.

Finally, applying change point detection may provide further insight into the temporal dynamics of habituation at different scales, via application to other developmental paradigms or imaging modalities [4]. More broadly, it may also allow researchers to evaluate the fidelity of the assumptions they make about data prior to processing or analysis [69].

## 5 Conclusion

This work introduces an approach to identify significant changes in the evoked hemodynamic response in fNIRS data, on a trial-by-trial basis. Using infant data from a rural Gambian population recorded using a HaND paradigm, the use of functional change point detection with wild binary segmentation has been tested and demonstrated to provide trial-specific insight into the timing of structural hemodynamic change. Group-level analysis from participants assessed at 5-, 8- and 12mo identified structural change corresponding to the evoked hemodynamic response. Further analyses indicated a high proportion of decreasing responses after FCPts at 8- and 12mo, and an age-related shift in the timing of decreasing hemodynamic responses, suggesting older infants in the first year of life habituate to auditory stimuli both sooner and with greater consistency. These findings demonstrate the utility of analyzing fNIRS data using FCPt detection and provide valuable insight into the development of habituation in the first year of infancy.

## Author Contributions (CRediT)

SB: conceptualization, data curation, formal analysis, methodology, software, visualization, writing – original draft, writing – review and editing; CE: funding acquisition, project administration, resources, writing – review and editing; SLF: funding acquisition, project administration, resources, writing – review and editing; EM: data curation, investigation, project administration; SMc: project administration, supervision, writing – review and editing; AB: data curation, supervision, writing – review and editing; SEM: funding acquisition, project administration, resources, supervision, writing – review and editing.

## Biographies

**Samuel Beaton** is a researcher in the Department of Women and Children’s Health at King’s College London. His research involves developing analytical pipelines for functional near-infrared spectroscopy (fNIRS) and high-density diffuse optical tomography (HD-DOT) data, using these to investigate brain development in infancy.

**Samantha McCann** is a Public Health Registrar and formerly Postdoctoral Research Associate in the Department of Women and Children’s Health at King’s College London. Her main research interest is supporting early child development, with a strong focus on the impact of undernutrition in infancy on long-term neurodevelopmental outcomes.

**Sarah Lloyd-Fox** is a Principal Research Associate in the Department of Psychology, University of Cambridge. She leads several multi-disciplinary projects focusing on developmental trajectories of early cognitive and brain development during pregnancy, infancy and early childhood. Her research focuses on understanding how family and environmental context - i.e. contextual factors such as poverty associated challenges and enriched multigenerational family support - shape early life.

**Clare Elwell** is a professor of medical physics leading projects to investigate brain function using functional near infrared spectroscopy in high and low resource settings in adult and infants.

**Ebrima Mbye** is a Field Coordinator at MRCG@LSHTM, formerly employed on the BRIGHT Project and currently engaged in the INDiGO Trial working under Professor Sophie Moore.

**Anna Blasi** is a Postdoctoral Research Fellow at UCL. Her research interests are centered on functional aspects of human physiology. Her research career started with models of the cardiovascular system and the effects of disease. Through her work at UCL, KCL, and Birkbeck, her research interests have shifted toward the use of functional imaging (fNIRS, fMRI) to study brain function and neurocognitive development in early infancy.

**Sophie Moore** is Professor of Global Women and Children’s Health in the Department of Women & Children’s Health at King’s College London and an Honorary Associate Professor at the London School of Hygiene and Tropical Medicine (LSHTM). Her research focuses on the nutritional regulation of ‘healthy’ fetal and infant growth, incorporating infant immune and brain development outcomes, and on the mechanisms through which maternal, infant and childhood nutrition may influence development and later health.

## Acknowledgments

We would like to place on record our thanks to the children, mothers and wider families who took part in this study. We also thank the BRIGHT Project study team in Keneba for their invaluable contribution to data collection and management.

## Code and Data Availability

The code used to conduct the analyses and generate the figures presented in this paper are available at https://github.com/sam-beaton/fcpt and rely on the following open source Python packages: *h5py* [70], *numpy* [71], *matplotlib* [72], *os* [73], *pandas* [74], *pickle* [75], *scikit-fda* [76], *scipy* [50], *statistics* [77], *sys* [78].

The data used to support this study are stored in the Brain Imaging for Global Health Data Repository. The conditions of ethics approval do not allow public archiving of pseudo-anonymized study data. The data cannot be fully anonymized due to the nature of combined sources of information, such as neuroimaging, sociodemographic, geographic and health measures, making it possible to attribute data to specific individuals. Hence, the release of data would not be compliant with GDPR guidelines unless additional participant consent forms are completed [12]. Access to any data collected during or generated by the BRIGHT project is fully audited and, to ensure data security, is overseen by the data management team in the UK and The Gambia. Relevant data sharing procedures were created in consultation with stakeholders and external consultation [79]. To access the data, interested readers should use the Contact page of the BRIGHT website. Access will be granted to named individuals following ethical procedures governing the reuse of sensitive data. Specifically, requestors must pre-register their proposal and clearly explain the planned analysis to ensure that the purpose and nature of the research is consistent with that to which participating families originally consented. Additionally, requestors must complete and sign a data sharing agreement to ensure data is stored securely. Approved projects would need to adhere to the BRIGHT project’s policies on Ethics, Data Sharing, Authorship and Publication.

## Appendices

### Appendix A Mathematical detail of FCPt detection

In a framework where functions are treated as the fundamental objects of analysis, mathematical and statistical computations must act on whole functions rather than on individual values. An important concept in FDA is therefore the operator: a mathematical mapping that takes one function as an input and produces either another function or a scalar as output [80]. A particularly important type of operator is the integral operator:

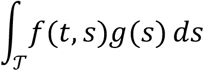

in which a function, *f*, ‘acts on’ another function, *g*, across the entire domain, 𝒯. This type of operator is crucial in defining the test statistic for FF FCPt detection, the mathematical detail of which constitutes the rest of this Appendix.

The fNIRS time series data represents a discretely sampled measurement of the underlying functional (hemodynamic response) process across some time range 𝒯. For each HaND paradigm trial (*n* = 1, …, *N*, where *N* = 25), we observe a function *Y*_*n*_(*t*), *t* ∈ 𝒯, sampled at discrete time points *t* = {𝒯_1_, …, 𝒯_*M*_}. We model the data as:

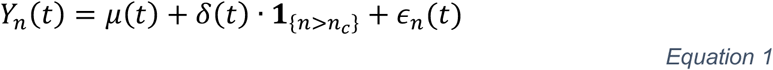

where, *μ*(*t*) is the mean function, *δ*(*t*) is a functional change activated after trial *n*_6_, and *ϵ*_*n*_(*t*) is a zero-mean stochastic error process representative of the measurement or systemic error. The null hypothesis:

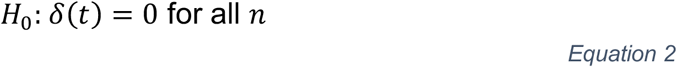

is tested. If there are no structural breaks, *H*_0_ holds. Under the alternate hypothesis, *H*_*A*_, *δ*(*t*) ≠ 0 for all *n* > *n*_*c*_ if there is a change point at some trial *n* = *n*_*c*_.

In each channel, age-grouped data was obtained by averaging across participants for each trial, yielding a single group-averaged function per trial, per chromophore in each channel, represented by Equation 1. The error term in Equation 1 is typically modeled with zero mean and covariance function: *c*(*t, s*) = *E*{*ϵ*_*n*_(*t*)*ϵ*_*n*_(*s*)}. This describes the linear association between functional observations *ϵ*_*n*_ at two different time points, *t* and *s*, during fNIRS recording. The covariance function is a key component of FCPt detection. In practice, *c*(*t, s*) is estimated using the sample covariance function:

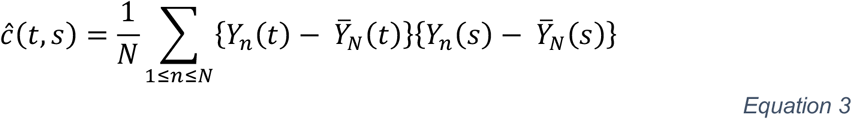

where 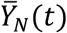 is the sample mean hemodynamic function:

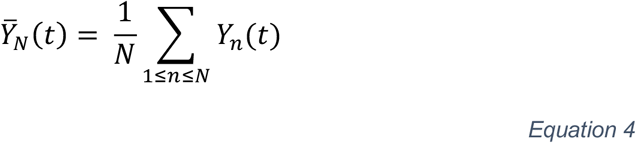

Potential change in the mean hemodynamic response function at all trials *n*_6_ was tested using the FF FCPt detection method [25], during which structural breaks in the mean function are examined using the covariance structure. This exploits the way deviations in the cumulative mean behavior of functional data are expected to behave under the null hypothesis (Equation 2).

For a particular trial *n* in the HaND paradigm (1 ≤ *n* ≤ *N*) it is possible to adapt the sample mean in Equation 4 to give the sample mean hemodynamic response function of the first *n* trials (i), and the final (*N* − *n*) trials (ii):

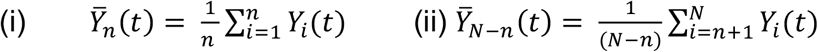

This can be used to give the weighted cumulative mean difference up to trial *n*:

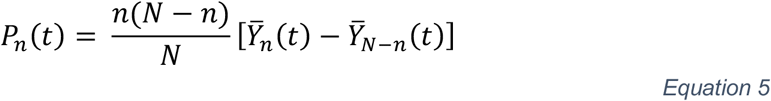

If the mean is constant, *P*_*n*_(*t*) will be small for all 1 ≤ *n* ≤ *N*; if a structural change occurs it will be large. The parabolic weight function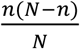 stabilises random variability which can occur when *n* is large (≈ *N*) or small [81, p. 84]. Substituting the full definitions of the two partial sums 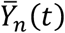 and 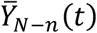 into Equation 4 and rearranging gives

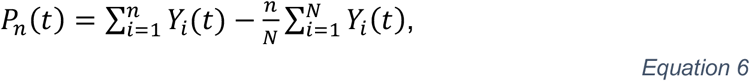

a cumulative sum (CUSUM) statistic, so named because it involves the sums of the weighted cumulative mean differences in Equation 5. *P*_n_(*t*) represents the deviation of the cumulative mean up to trial *n* from the overall mean across all *N* trials. This motivates the test statistic 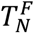:

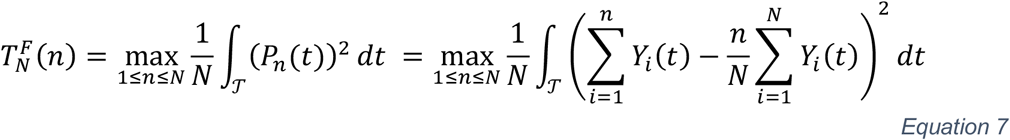

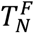 represents the trial number *n* which produces the largest absolute difference between the cumulative sum of *Y*_*n*_ and all its antecedents, and the mean function across all functional observations weighted by 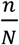 evaluated across the entire (time) domain. Normalising by *N* ensures 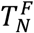 is well scaled.

In intuitive terms, 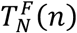 measures how much the average hemodynamic response across the first n trials differs from the overall average across all trials. These so-called cumulative differences fluctuate randomly around zero in the case of a stable underlying hemodynamic response or when no consistent pattern is present in the data. Under the alternative hypothesis, *H*_*A*_, when the shape or magnitude of the mean response changes – such as when the hemodynamic response declines with habituation – the cumulative difference grows sharply due to deviation within each trial caused by *δ*(*t*) in Equation 1. The trial number, *n*_*c*_, at which this divergence reaches its largest value marks the most likely location of a structural change in the data. In practice, this “cumulative-sum” (CUSUM) approach captures the trial at which the signal’s average behavior shifts most strongly over time (full mathematical formulation in Appendix A, Equation 7).

Each channel/chromophore combination contains a functional time series of 25 functions corresponding to the 25 trials in the HaND paradigm. For each of these channel/chromophore combinations, candidate change points, *n*^∗^, for each functional time series were identified, corresponding to trial numbers where the largest structural change occurred. Candidate FCPts were identified as the smallest *n*^∗^ such that 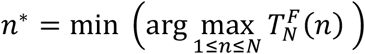: the smallest (earliest occurring) trial number which produces the largest cumulative difference from the overall functional mean in each channel/chromophore combination.

Following the recommendations of the developers of the FF method the group-averaged fNIRS data functional object in the trial window was estimated using a set of 21 basis spline functions [16, pp. 46–47], [17], [25].

### Appendix B FCPt significance testing with Brownian Bridges

(i) To assess the size of 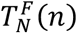 values for the data and the significance of the associated change points, the distribution of the test statistic under *H*_0_ is required, which relies on the estimated covariance, *ĉ* (*t, s*) function, as defined in Appendix A. An eigendecomposition of the associated estimated covariance integral operator ∫*ĉ* (*t, s*) *ds* provides its estimated eigenfunctions 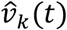 and eigenvalues 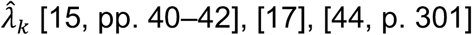:

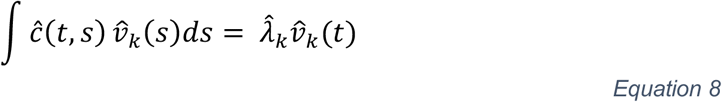

Eigenfunctions characterize the dominant modes/shapes of variation in functional data. The *k*^*th*^ eigenvalue represents the amount of variation corresponding to the *k*^*th*^ eigenfunction. The importance of the eigenvalues 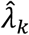 for 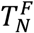 is seen in its convergence in distribution, *D*, under *H*_0_:

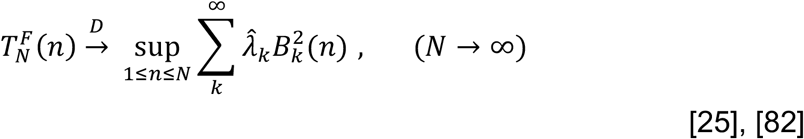

where *B*_*k*_(*n*) is a Brownian Bridge on the same values of *n* – a stochastic process *B*_*k*_(*n*) = *W*(*n*) − *nW*(*n*) such that *B*_*n*_(0) = *B*_*n*_(*N*) = 0 and *W*_n_(*n*) is a standard Brownian motion (main text, Figure 3b). In other words, the test statistic 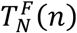 asymptotically converges to a weighted Brownian Bridge distribution under *H*_7_ [22], [47].

This convergence in distribution provides a possible method for significance testing using empirical critical values of the distribution itself [81]. To permit multiple comparison corrections, however, it was necessary to calculate exact significance values, so the distribution of 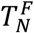 was approximated using Monte Carlo simulation.

During Monte Carlo simulation, for each *n*^∗^, the maximum value for the *m*^*th*^ Brownian Bridge, *B*_*m*_, was given by

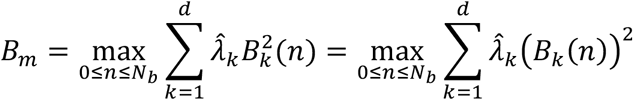

where *N*_*b*_ is the length of the interval being assessed at each stage of the recursive wild binary segmentation process. The *B*_*m*_model random fluctuations under the null hypothesis; 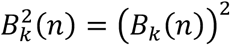 quantifies the variability contributed by each component; multiplying by 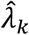 ensures more dominant modes of variability contribute more to the sum. Statistical testing of the significance of candidate change points *n*^∗^ was therefore conducted by comparing their associated test statistics to the maximum values of *M* = 10 000 generated using summed Brownian Bridges of length *N*_*b*_. The statistical significance of 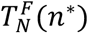 was then given by the proportion of the *M* randomly generated values larger than 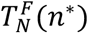.

(ii) Each comparison with Brownian Bridge distributions during wild binary segmentation amounts to a separate statistical test and increases the family-wise error rate (FWER). To correct for this, within-channel multiple comparison correction using the Bonferroni-Holm method [48] was conducted, using the generated significance values. To address the hierarchical nature of the binary segmentation process, in which the importance of change points at higher levels is predicated on the statistical significance of those which occur beforehand (see Figure 3) a step-down multiple comparison procedure was conducted within each channel at every binary search level. Each hierarchical level, *l*, of binary segmentation has a maximum of 2^*l*−1^ candidate change points; correction for a maximum of ∑_*l*_ 2^*l*−1^ tests at each level was therefore conducted, corresponding to the change points at the current and all preceding levels. If a test at any level failed to meet the significance threshold post-correction, all descendant tests were discarded. The Bonferroni-Holm method was applied at each level using the *statsmodels* Python package [48], [49]. An alpha value of 0.01 was used to determine the statistical significance of change points.

