## Supplementary Materials for "Characterizing developmental changes in infant habituation using functional change point detection"

#### Supplementary Materials 1: Additional FDA concepts and relevant detail

Within FDA, functions can form a mathematical space (a set of objects with underlying structure/rules); the most common is $L^2$-space which consists of square-integrable functions. Functions, $f(t)$, are square-integrable if the integral of its squared values across the entire functional domain, $\Omega$, is finite:

$$\int_T f^2(t)\, dt < \infty$$

*Equation S 1*

The variable name $t$ and functional domain name $T$ are chosen deliberately, as FDA is often used to model time-varying phenomena. $L^2$-space contains only functions with finite energy (i.e. not wildly erratic – see Figure S 1) and size (does not contain infinite values or measures– see Figure S 1). Size is measured by the $L^2$-norm, the square-root of the left-hand side of Equation S 1 [15, p. 38], [83, p. 88] $L^2$-space permits the definition of notions such as orthogonality, distance, and length [15, pp. 38–39].

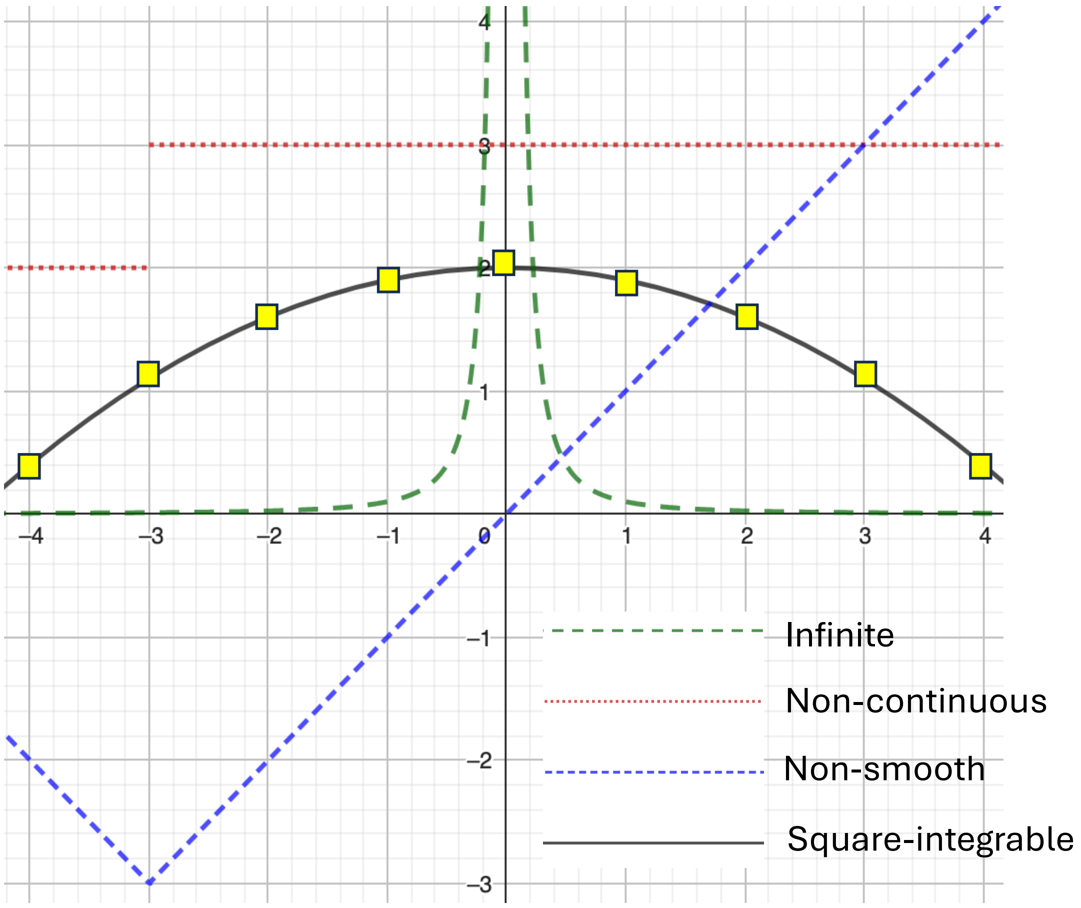


***Figure S 1: Discrete, functional, and square-integrable data.***

*The yellow boxes represent discrete sampling points of the underlying functional process given by the black curve. On the domain* $-4 \leq x \leq 4$ *the black curve is square-integrable, as the square of its integral is finite (the area under the curve can be accounted for). The other curves are not square integrable. The blue function is not smooth: at (−3, −3) there is a sharp turn which has no derivative. The red function is discontinuous at* $x=-3$*. The green function has no upper limit when* $x=0$*, meaning it does not have a finite integral.*

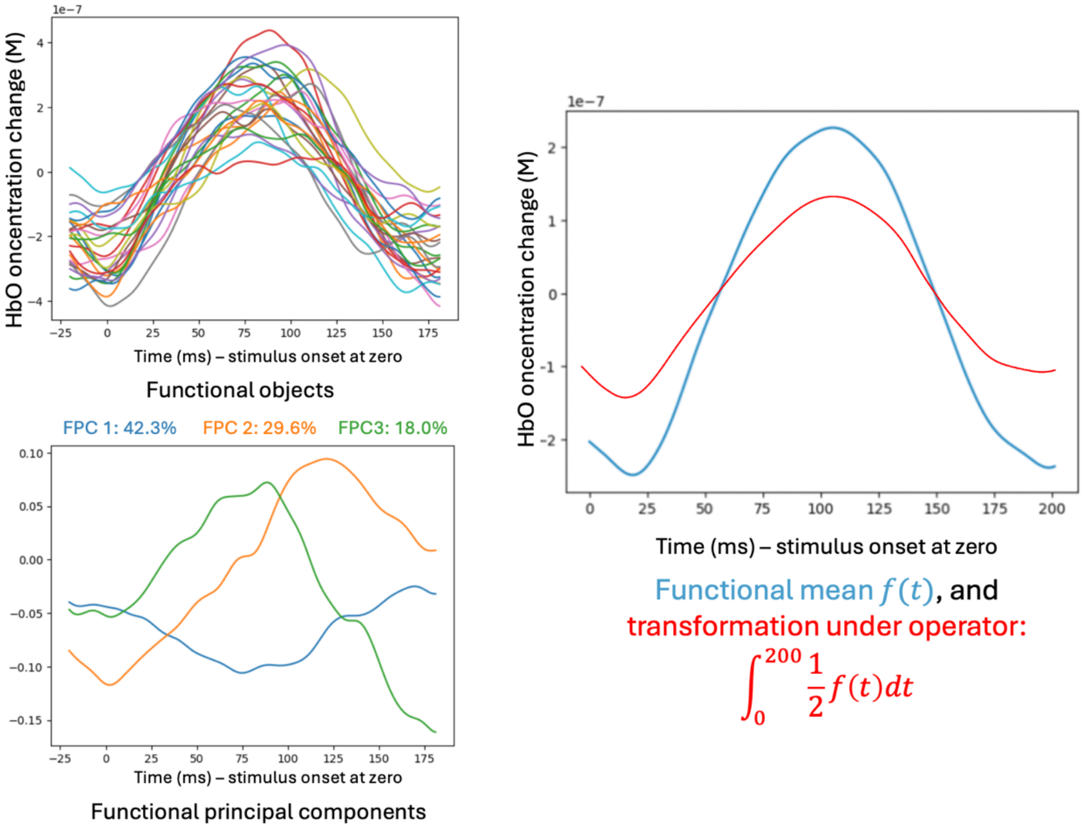


***Figure S 2: Examples of key concepts in FDA.***

*(a) functional observations, representing hemodynamic responses. (b) the functional mean of the observations in (a), plus a transformation under a given operator. (c) Functional principal components of the 25 functional observations.*

#### Supplementary Materials 2: FPCA & Drift

Mean functional principal components of individual functional observations in a single channel at 12mo. FPC2 shows the drift that was not originally removed using a high-pass filter.

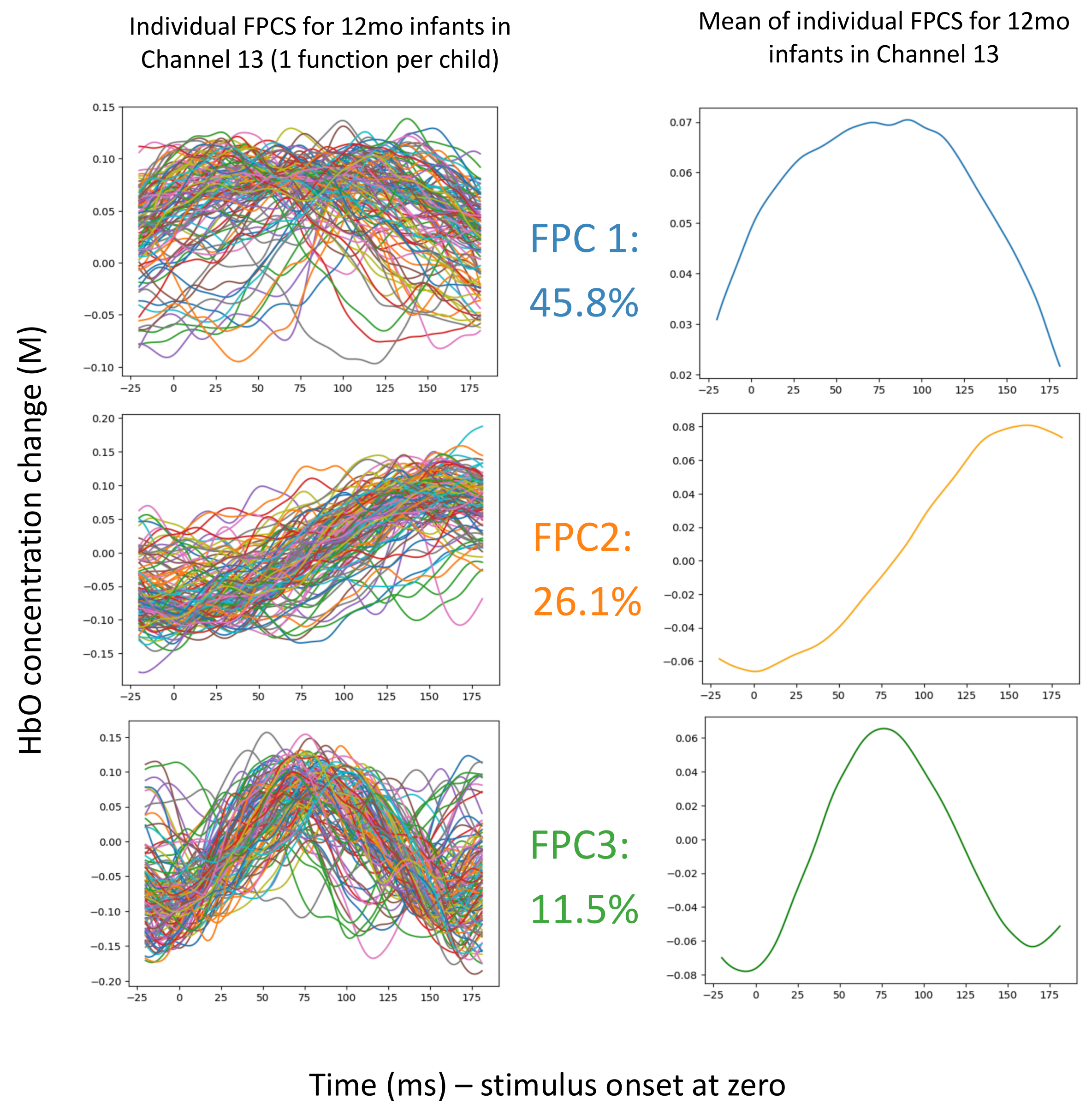


*Figure S 3: Functional data examples.*

#### Supplementary Materials 3: FPCA-based change point detection

The second FCPt detection method used employs dominant modes of data variation captured by FPCA. For full details, see Aue and colleagues [84], Berkes and colleagues [22] and Horváth and Kokoszka [44]. This method accurately detects change points when structural change align with dominant FPCs [25], [85]. Since exploratory analysis of HaND fNIRS data suggested the hemodynamic response corresponds one of the first 3 FPCs, depending on the individual (Supplementary Materials 2), this was a reasonable assumption. By projecting data onto a lower-dimensional basis, this method permits change point detection whilst decreasing data dimensionality but also risks information loss and subsequent false negatives during change point detection.

The eigenfunctions discussed prior to Equation 8 are those functions which remain unchanged under the estimated covariance operator aside from potential scaling (given by the eigenvalues). This is evident in the right-hand side of Equation 8 where the function $\hat{v}_k$ is multiplied by its corresponding eigenvalue $\hat{\lambda}_k$. The eigenfunctions form a fundamental set of functions (a 'basis') which can be used as building blocks to represent other functions. The sample covariance function (Equation 3) can therefore be expanded using its estimated eigenfunctions and eigenvalues:

$$\hat{c}(t,s) = \sum_k \hat{\lambda}_k \hat{v}_k(t)\hat{v}_k(s)$$

via the Karhunen-Loève (K-L) expansion [15, p. 11], [16, p. 152]. The K-L expansion for functional data is analogous to the multivariate representation of vectors using a linear combination of eigenvectors: an object is represented using a weighted sum of basis objects which act as building blocks – in this case the covariance function is represented using a combination of the eigenfunctions.

The error functions, $\epsilon_n(t)$, can also be represented using the estimated eigenfunctions:

$$\epsilon_n(t) = \sum_k \hat{\eta}_{n,k}\hat{v}_k(t)$$

*Equation 9*

where $\hat{\eta}_{n,k}$ is the score, or coefficient, corresponding to the contribution of the $k^{th}$ eigenfunction $\hat{v}_k$ to the representation of the $n^{th}$ error function $\epsilon_n$, used to model the $n^{th}$ hemodynamic response function $Y_n(t)$. Consequently, the eigenfunction expansion of the error term in Equation 9 can be used to rewrite Equation 1:

$$Y_n(t) = \mu(t) + \delta(t)\cdot \mathbf{1}_{\{n>n_c\}} + \sum_k \hat{\eta}_{n,k}\hat{v}_k(t)$$

*Equation 10*

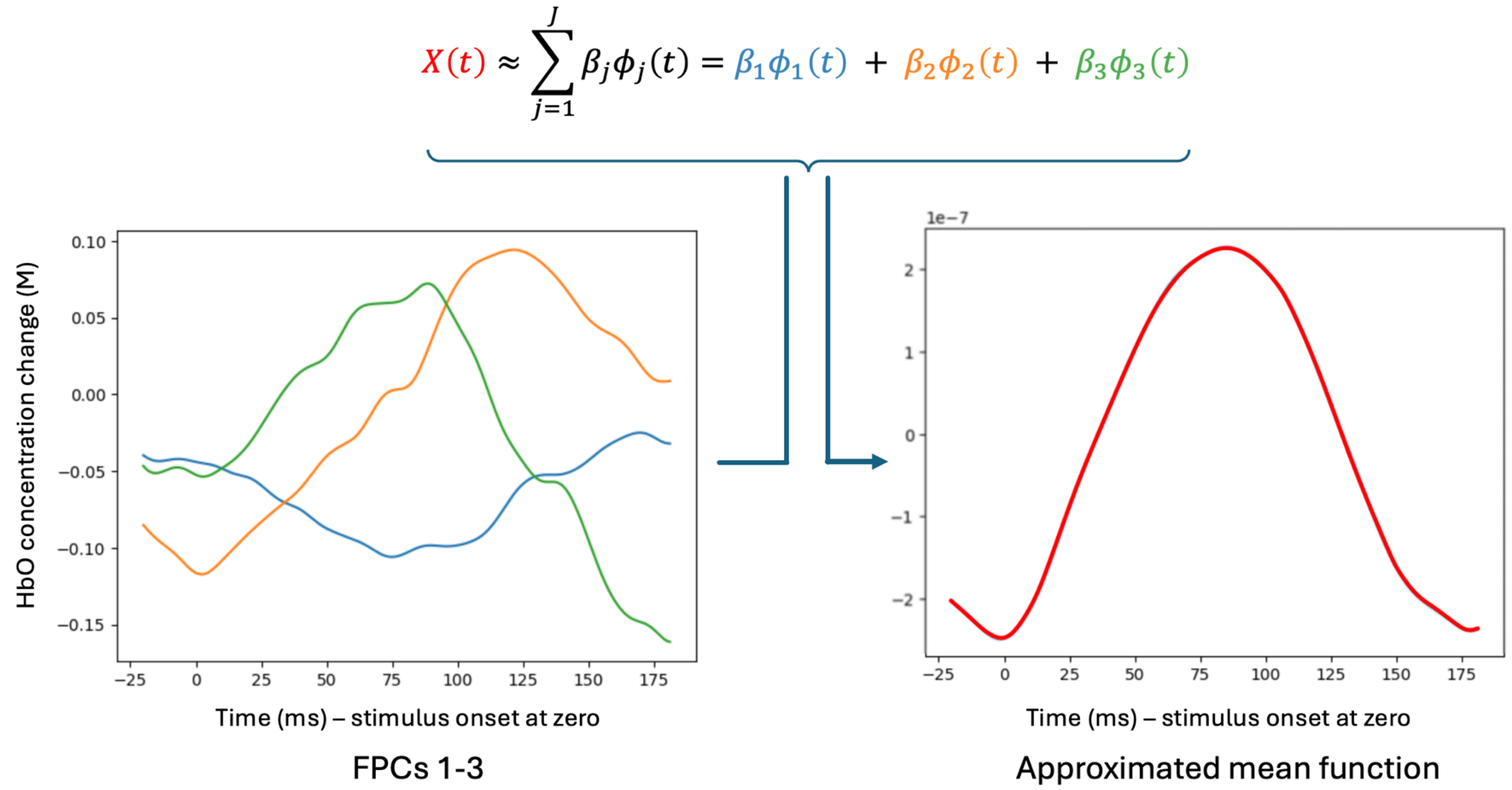


***Figure S 4: An example representation of the mean hemodynamic response function, using the Karhunen-Loève expansion truncated at J=3 principal components, demonstrating the way functional objects in L^2-space can be composed using other basis objects.***

FPCA reduces the dimensionality of this representation by retaining only the most dominant $d$ eigenfunctions. It is common to choose $d$ such that the first $d$ eigenfunctions account for at least 85% of the variance [15, p. 256], which is the approach taken, conducting the FPCA itself using the *scikit-fda* Python package [76]. This leads to the truncated K-L expansion for observed hemodynamic response function $y_i$:

$$Y_n^d(t) \approx \mu(t) + \delta(t) \cdot \mathbf{1}_{\{n>n_c\}} + \sum_{k=1}^{d} \hat{\eta}_{n,k}\hat{\upsilon}_k(t)$$

*Equation 11*

This expansion is exploited during the FCPt detection approach defined by Berkes and colleagues [22] and extended by Horváth and Kokoszka [44], [86]. Since each $Y_n(t)$ are functional and hence infinite-dimensional, it is helpful to work with the projection of $P_n(t)$ on to eigenfunctions:

$$\int P_n(t)\hat{\upsilon}_k \, dt = \int \left[\sum_{i=1}^{n} Y_i(t) - \frac{n}{N}\sum_{i=n+1}^{N} Y_i(t)\right]\hat{\upsilon}_k \, dt$$

*Equation 12*

because one can then choose to work with the leading eigenfunctions and eigenvalues which capture most of the data variance. Under $H_0$, Equation 10 reduces to

$$Y_n(t) = \mu(t) + \sum_{k} \hat{\eta}_{n,k}\hat{\upsilon}_k(t)$$

*Equation 13*

which means each of the scores, $\hat{\eta}_{n,k}$, can be found by projecting the mean-centered function $Y_n(t)$ on to each of the eigenfunctions:

$$\hat{\eta}_{n,k} = \int (Y_n(t) - \bar{Y}_N(t))\hat{\upsilon}_k(t) \, dt$$

*Equation 14*

[15, p. 41], [81, p. 84]

Centering the mean functions by replacing $Y_n(t)$ with $Y_n(t) - \bar{Y}_N(t)$ does not change Equation 5; consequently, Equation 12 can be rewritten as

$$\int P_n(t)\hat{\upsilon}_k \, dt = \int \left[\sum_{i=1}^{n}\left(Y_i(t) - \underline{Y}_N(t)\right) - \frac{n}{N}\sum_{i=n+1}^{N}\left(Y_n(t) - \underline{Y}_N(t)\right)\right]\hat{\upsilon}_k \, dt$$

which simplifies to

$$\int P_n(t)\hat{\upsilon}_k \, dt = \sum_{i=1}^{n} \hat{\eta}_{i,k} - \frac{n}{N}\sum_{i=1}^{N} \hat{\eta}_{i,k}$$

*Equation 15*

Equation 15 is another CUSUM-type statistic and shows that the scores for each fNIRS functional observation can be used, with respect to the eigenfunctions, to test for mean constancy. It is used in the following test statistic:

$$T_N^P(n) = \frac{1}{N}\sum_{k=1}^{d}\frac{1}{\hat{\lambda}_k}\left[\sum_{i=1}^{n}\hat{\eta}_{i,k} - \frac{n}{N}\sum_{i=1}^{N}\hat{\eta}_{i,k}\right]^2$$

*Equation 16*

Squaring the mean differences and normalising by $N$ have the same rationale as for $T_N^F(n)$. Normalising by the eigenvalues $\hat{\lambda}_k$ ensures those capturing large variation do not dominate the statistic; summing across scores corresponding to the $d$ leading eigenvalues captures most of the variability.

Similarly to $T_N^F$, under $H_0$ the test statistic $T_N^P(n)$ also converges to a sum of squared Brownian Bridges:

$$T_N^P(n) \overset{D}{\rightarrow} \sup_{1\le n\le N}\sum_{k=1}^{d} B_k^2(n), \quad (N \rightarrow \infty)$$

*Equation 17*

[44], [82, p. 2]

In contrast to the case for $T_N^F$, the sum is not infinite since the approach explicitly includes only the most dominant $d$ eigenfunctions, and the sum is not weighted as the test statistic already accounts for the differing variability of the eigenfunctions (see Equation 16). Under $H_A$, deviation caused by $\delta(t)$ in Equation 10 will lead to an unexpected increase in $T_N^P(n)$ near the true break point $n_c$. Candidate change points, $n^*$, were identified for each functional time series of fNIRS HaND paradigm trial data, as $n^* = \min\left(\arg\max_{0\le n\le N} T_N^P(n)\right)$. As with the FF method, Monte Carlo simulation was used to generate critical values to assess statistical significance of the change points.

Change point detection using the FPCA-based method was as described using the FF-based approach, using wild binary segmentation and distributions of test-statistics generated using Brownian Bridges. The key differences are (i) the test statistic used, which for FPCA-based detection was $T_N^P$, and the removal of the weights for the Brownian Bridges (Equation 17) since these are already accounted for in $T_N^P$ itself (Equation 16).

#### Supplementary Materials 4: Data retention

Sample size, data retention, reasons for exclusion or withdrawal, and descriptive statistics for remaining fNIRS data are detailed in the Figure below.

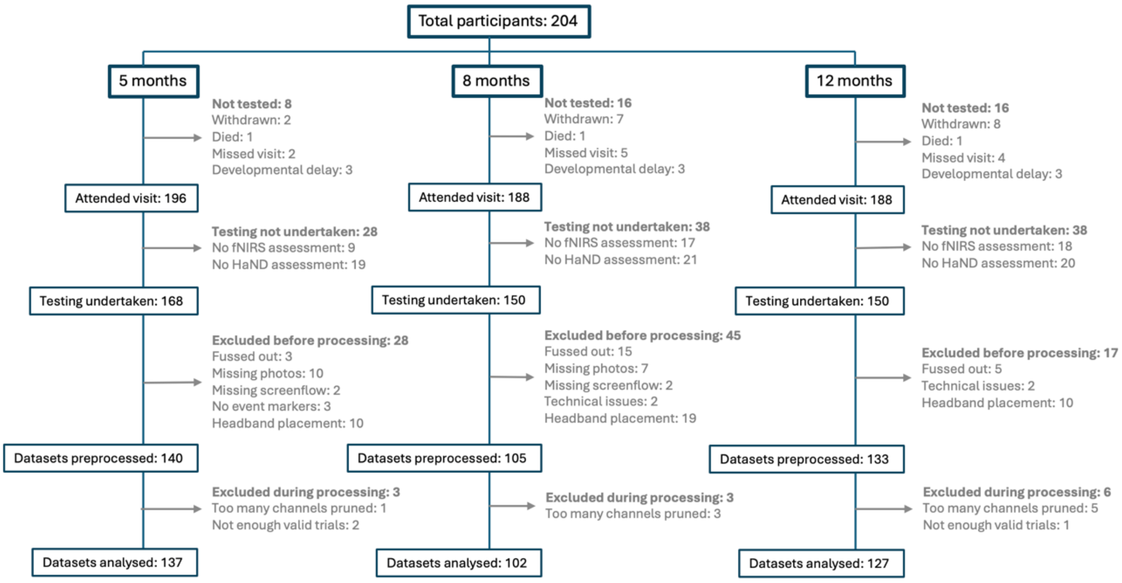


***Figure S5: Data retention for each of the 3 ages during HaND fNIRS assessment and processing.***

#### Supplementary Materials 5: Channel overlap with previous univariate analyses

The rationale for this is as follows: the detection of significant change in the form of FCPts necessarily requires a sufficiently consistent underlying data structure for a significant deviation from this structure to be detected; given the time-locked evoked hemodynamic response is likely to be one of few physiological components in the data which is consistent across trials, it should be the case that FCPt detection is sensitive to this structure.

##### Assessing channel overlap with previous analysis

For this specific dataset, previous work has been published which used $t$-test contrasts [8] and spatio-temporal clustering [12] to locate channels co-located with activation. FCPt detection is intended to complement analytical approaches to determine the location of evoked hemodynamic responses, as FDA methods already exist for one- and two-sample problems [87], [88]. Nevertheless, consistent structure across a set of functions is required for large test statistic values, which in this case would indicate a temporal structure time-locked to the HaND paradigm's stimulus presentation. Overlap with previously identified activation channels would indicate that FCPt detection captures physiologically meaningful structure.

Accordingly, agreement was assessed between channels with detected change points and those channels in which a hemodynamic response was identified across the first 5 trials, derived from previous work using the spatiotemporal clustering method Threshold-Free Cluster Enhancement (TFCE) [12], [89]. Using the survival function of the hypergeometric distribution, implemented with the Python class *scipy.stats.hypergeom* [50] the probability of observed overlap between the sets of FCPt channels generated using both the FF and FPCA-based FCPt detection method, and the set of channels with a hemodynamic response according to TFCE, was assessed. The channels identified for (i) HbO, (ii) HbR, and (iii) both chromophores were compared.

This analysis seeks to check inherent sensitivity to the data structure and detect meaningful overlap with prior work, so controlling the false discovery rate was prioritised rather than strict control of the FWER. Consequently, Benjamini-Hochberg's method for multiple comparison correction was conducted across 5-, 8- and 12mo data using the *statsmodels* Python package [49], [90].

**(a) Channel overlap between both functional change point detection methods and overall set of TFCE analysis channels.**

*Table S 1: Overlap between the two functional change point detection methods and cross-age ROI channels outlined using TFCE analysis.*

| | | | Fully functional ($T_N^F$) | | | FPCA ($T_N^P$) | | |
|---|---|---|---|---|---|---|---|---|
| Age | Chromophore | $K$ | $n$ | $k$ | $p$-value (adj.) | $n$ | $k$ | $p$-value (adj.) |
| 5mo | HbO | 10 | 15 | 6 | *0.307* | 1 | 0 | *1* |
| | HbR | 8 | 10 | 4 | **0.047*** | 7 | 4 | *0.112* |
| | Both | 6 | 6 | 4 | **0.004**** | 0 | 0 | *1* |
| 8mo | HbO | 10 | 10 | 4 | *0.316* | 5 | 4 | *0.057* |
| | HbR | 8 | 14 | 7 | **0.012*** | 4 | 2 | *0.344* |
| | Both | 6 | 6 | 4 | **0.004**** | 2 | 2 | *0.080* |
| 12mo | HbO | 10 | 12 | 7 | **0.031*** | 6 | 4 | *0.072* |
| | HbR | 8 | 11 | 5 | *0.052* | 2 | 1 | *0.421* |
| | Both | 6 | 6 | 4 | **0.004**** | 1 | 1 | *0.265* |

*Change points were identified using $\alpha$=0.01 after Benjamini-Hochberg correction. The fully functional method shows overlap for a larger number of chromophores/combinations. Key: $K$ = number of cross-age channels identified by TFCE; $n$ = number of channels identified by change point detection method; $k$ = number of channels which overlap between the two methods. Significance in overlap was determined using the survival function of the hypergeometric distribution and is rounded here to 3 decimal places ( $p$*<0.05; $p$**<0.01).*

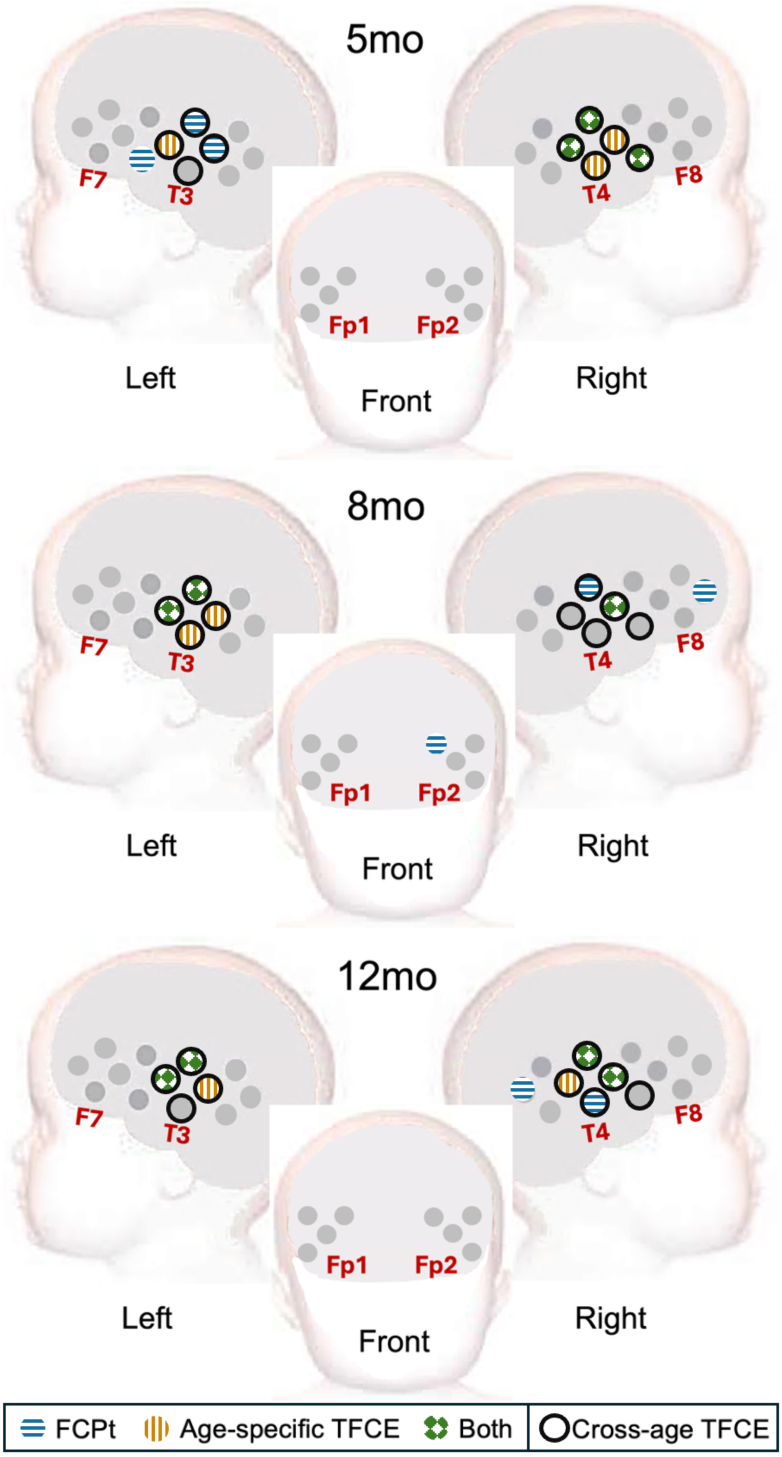


***Figure S6: Channel overlap between FF FCPt and TFCE methods***

*Blue, horizontal stripes are channels with identified FCPts. Orange, vertical lines represent age-specific channels with significant hemodynamic responses across all three ages (5-, 8- and 12mo) identified in the first 5 trials of the HaND paradigm, using the TFCE approach. Green, checkerboard represents channels which overlap between the two analyses. Black, dashed-circled channels are ROI channels for all 3 ages. Black, dashed-line square icons represent channels with identified FCPts in both chromophores for all 3 ages.*

1544

##### (b) Channel overlap between both functional change point detection methods and age-specific TFCE analysis channels.

*Table S 2: Overlap between the two functional change point detection methods and age-specific channels outlined using TFCE analysis.*

| | | | Overlap | | | | | |
|---|---|---|---|---|---|---|---|---|
| | | | Age specific TFCE channels | | | Overall TFCE channels | | |
| Age | Chromophore | $n$ | $K$ | $k$ | $p$-value (adj.) | $K$ | $k$ | $p$-value (adj.) |
| 5mo | HbO | 15 | 11 | 7 | 0.112 | 10 | 6 | 0.307 |
| | HbR | 10 | 8 | 4 | 0.154 | **8** | **4** | **0.047*** |
| | Both | 6 | 6 | 3 | 0.053 | **6** | **4** | **0.004**** |
| 8mo | HbO | 10 | 9 | 5 | 0.090 | 10 | 4 | 0.316 |
| | HbR | 14 | 6 | 5 | 0.078 | **8** | **7** | **0.012*** |
| | Both | 6 | **5** | **3** | **0.043*** | **6** | **4** | **0.004**** |
| 12mo | HbO | 12 | **10** | **7** | **0.030*** | **10** | **7** | **0.031*** |
| | HbR | 11 | 8 | 5 | 0.078 | 8 | 5 | 0.052 |
| | Both | 6 | **6** | **4** | **0.013*** | **6** | **4** | **0.004**** |

*Despite the lowest overall significance value exhibited by the FPCA-based FCPt detection method, for HbO data at 8mo, the FF method showed a pattern of greater overlap overall. with overlap for a larger number of age/chromophore combinations. In addition to the three statistically significant age/chromophore combinations for the FF FCPt detection method, an additional four tests produced marginal significance values, and the highest overall significance value for this method was 0.154. Key: $K$ = number of age-specific channels identified by TFCE; $n$ = number of channels identified by change point detection method; $k$ = number of channels which overlap between the two methods. Change points were identified using $\alpha = 0.01$ after Bonferroni-Holm correction. Significance in overlap was determined using the survival function of the hypergeometric distribution ( $p$*<0.05; $p$**<0.01).*

##### (c) Discussion of lack of overlap of the FPCA-based FCPt detection approach with channels identified by the prior TFCE analysis

The weaker overlap between FPCA-based FCPt channels and those identified by TFCE analysis when compared to the fully functional FCPt detection approach likely stems from identifying fewer channels. Raising the threshold to 0.05 as verification still yielded the same number of chromophore and channel combinations with significant overlaps, but a better overlap with channels identified for both chromophores than for the original $\alpha = 0.01$ change point threshold. These channels were similar spatially to those found using the higher 0.01 threshold for the FF method (see Supplementary Materials 7), confirming preference for the FF method. The comparative reduction in accuracy may be attributed to changes in the hemodynamic response occurring in sources of variation not captured by the orthogonal FPCs [25], [85].

The suitability of the number of FPCs used in the FPCA method was also evaluated, given that FPCA-based FCPt detection is known to be sensitive to noise

[91] and may require a greater number of FPCs in noisier datasets. When FPCs accounting for 90% and 95% of the data variation were used for verification, the resulting number of detected change points was prohibitively small for meaningful statistical comparison with TFCE channels. Using a similar number (four) of basis functions for the FF method as was typically utilized in the FPCA-based method yielded no significant overlap with the TFCE channels (Supplementary Materials 9). This suggests the FPCA-based method outperforms the FF method with fewer basis functions and that the underperformance of the FPCA method is due to an insufficiently detailed data representation. However, the lack of improvement when using greater numbers of FPCs during the FPCA-based approach precludes comparison of the methods when capturing the data structure sufficiently accurately and is therefore an inherent limitation of this particular FPCA-based method, and likely others.

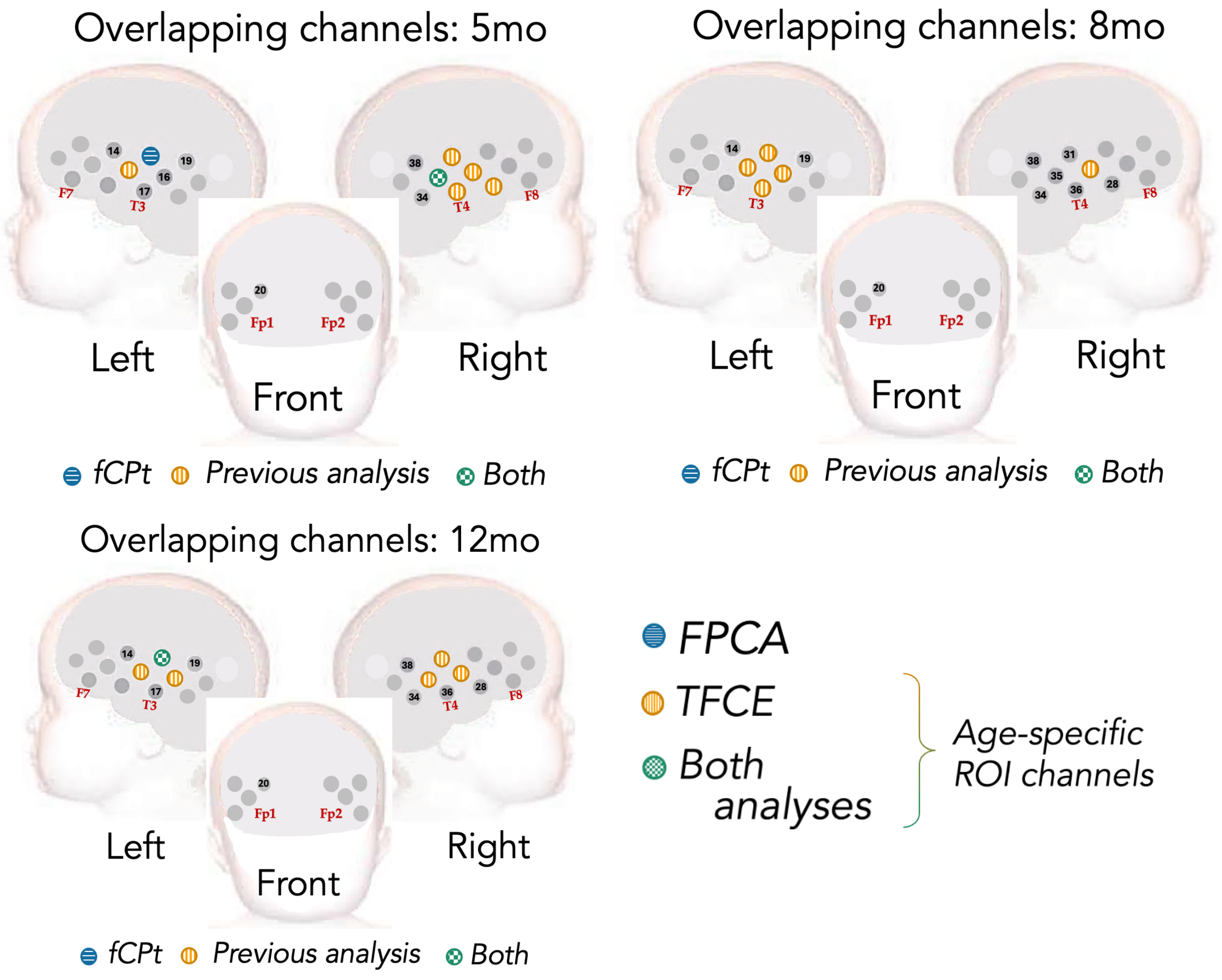


*Figure S7: FPCA Change point channels and overlap with age-specific channels*

#### Supplementary Materials 6: Full results from analysis of proportion of decreasing FCPts

**Table S3: Proportion of decreasing FCPts which occurred first, last, or anywhere during the channels' time functional series**

| | | Cross-age channels | | | | Age-specific channels | | | |
|---|---|---|---|---|---|---|---|---|---|
| Age | Change points | Total | Total | Total | Total | Total | Decreasing | Proportion (%) | $p$-value (adj.) |
| All | ALL | **43** | **28** | **28** | **28** | **28** | **37** | **86.0** | **<0.001*** ** |
| | First | **21** | **14** | **14** | **14** | **14** | **18** | **85.7** | **0.005** ** |
| | Last | **21** | **14** | **14** | **14** | **14** | **17** | **80.1** | **0.013*** |
| 5mo | ALL | *13* | *8* | *8* | *8* | *8* | *8* | *61.5* | *0.581* |
| | First | *6* | *4* | *4* | *4* | *4* | *4* | *66.7* | *0.688* |
| | Last | *6* | *4* | *4* | *4* | *4* | *2* | *33.3* | *0.688* |
| 8mo | ALL | **16** | **10** | **10** | **10** | **10** | **16** | **100** | **<0.001*** ** |
| | First | **8** | *5* | *5* | *5* | *5* | **8** | **100** | **0.013*** |
| | Last | **8** | *5* | *5* | *5* | *5* | **8** | **100** | **0.013*** |
| 12mo | ALL | **14** | **10** | **10** | **10** | **10** | **13** | **92.3** | **0.005** ** |
| | First | *7* | *5* | *5* | *5* | *5* | *6* | *85.7* | *0.167* |
| | Last | **7** | *5* | *5* | *5* | *5* | **7** | **100** | **0.023*** |

*Significance reported after Bonferroni-Holm correction (p*<0.05; p**<0.01, p***<0.001). For the age-specific channels, all detected 8- and 12mo FCPts were decreasing; for the overall channels, only one FCPt did not decrease within the two ages: one of the first FCPts at 12mo Proportions for the 5mo data were more varied, with no FCPt category containing a significant number of decreasing changes either in age-specific or cross- age ROI channels.*

#### Reporting for age-specific channels

At 5mo, no statistically significant differences were observed in the proportion of decreasing change points across all categories of change points ($p$-values > 0.05). At 8mo all detected change points were decreasing there were significant proportions of decreasing change points when considering all detected change points ($p$-value $= 0.006$ for both ages). No statistical significance was observed for the sets of first or last change points in each channel, for either 8- or 12mo ($p$-values > 0.05). Despite this, the relevant significance values were all 0.083, and examination of the proportion of decreasing FCPts shows that in all four cases, all five of five possible identified FCPts were decreasing. This suggests that small sample size is likely driving these $p$-values which fail to meet the significance threshold, rather than the proportion itself.

#### Supplementary Materials 7: Full results for analysis of age-effects on FCPts

##### (a) Full weighted ordinal regression results

**Table S4: Weighted ordinal logistic regression results for FCPts**.

| | Cross-age | | | | Age-specific | | | |
|---|---|---|---|---|---|---|---|---|
| Change points | $\beta$ | S.E. | $z$ | $p$ (adj.) | $\beta$ | S.E. | $z$ | $p$ (adj.) |
| Decreasing | **-0.247** | **0.114** | **-2.17** | **0.030*** | **-0.272** | **0.117** | **-2.33** | **0.020*** |
| First And Decreasing | *-0.216* | *0.165* | *-1.30* | *0.192* | *-0.216* | *0.165* | *-1.30* | *0.192* |
| Final And decreasing | **-0.497** | **0.218** | **-2.28** | **0.022*** | **-0.495** | **0.218** | **-2.27** | **0.023*** |
| Earliest Decreasing | *-0.195* | *0.145* | *-1.34* | *0.180* | *-0.195* | *0.145* | *-1.34* | *0.180* |
| Latest Decreasing | **-0.359** | **0.175** | **-2.06** | ***0.040**** | **-0.424** | **0.176** | **-2.41** | ***0.016**** |

*Detected across both chromophores, after Bonferroni-Holm correction. Significant effects for age were found in the change points which decreased, particularly those occurring later in the series of 25 trials. Significance: p*<0.05; p**<0.01, p***<0.001. Age-specific: channels in which a significant hemodynamic response had been identified using conventional analyses; overall: cross-age set of channels in which a significant hemodynamic response had been identified using conventional analyses.*

##### (b) Reporting of age-specific results

Similar results to those for cross-age channels were found when analyzing the age-specific set of channels with hemodynamic responses for both chromophores. Weighted ordinal logistic regression results for age-related differences in the timing of Final change points was identical (see above Table) – as were the post-test findings (see Table below). Both the effect size and statistical significance of age on Decreasing ($\beta = -0.272,\ SE = 0.117, z = -2.33,\ p = 0.020$) and Latest Decreasing ($\beta = -0.424,\ SE = 0.176, z = -2.41,\ p = 0.016$) change points decreased slightly, whereas those for the Final And Decreasing FCPts decreased slightly ($\beta = -0.495,\ SE = 0.218, z = -2.27,\ p = 0.023$). Wald tests (see Table below) confirmed the significance of these effects; pairwise differences were again driven by later occurrences at 5mo than the two later Ages for Final And Decreasing (5- & 8mo: $M = -4.54, p = 0.006$; 5- & 12mo: $M = -4.81, p < 0.001$) and Latest Decreasing (5- & 8mo: $M = -3.08, p = 0.049$; 5- & 12mo: $M =$

$-3.34, p = 0.036$). Interestingly, the earliest occurring FCPts were relatively consistent across age groups, with no other significant results found. No significant effects for the Change Point Count Per Channel covariate were identified for any of the FCPt types.

In all three cases, Tukey's HSD results for age-specific channels showed that differences between age groups were driven by later change point occurrences at 5mo than the two later time points. Decreasing change points overall occurred a mean of 3.58 trials later than at 8mo ($p = 0.026$) and 3.66 trials later than at 12mo ($p = 0.029$). More specific analysis of late-occurring, decreasing change points showed that change points which were both Final And Decreasing had mean trial occurrences 4.54 later at 5mo when compared to 8mo (p=0.006) and 4.81 trials later when compared to 12mo ($p = 0.005$). Similarly, the Latest Decreasing change points occurred an average of 2.88 trials later at 5mo than at 8mo ($p = 0.046$) and 3.14 trials later than at 12mo ($p = 0.033$). No significant pairwise effects were found for Decreasing change points. No significant effects of Change Point Count Per Channel were found for any of the change point types.

##### (c) Post-hoc testing for age-specific channels

**Table S5: Post-hoc testing of change point categories for which significant channel effects were identified using weighted ordinal logistic regression**.

| | | Wald test | | 5 & 8 pairwise | | 5 & 12 pairwise | | 8 & 12 pairwise | |
|---|---|---|---|---|---|---|---|---|---|
| Channels | Change points | $\chi^2(1)$ | $p$ | $M$ | $p$ | $M$ | $p$ | $M$ | $p$ |
| Cross-age | Decreasing | **4.70** | **0.030*** | *-3.15* | *0.062* | *-3.24* | *0.066* | *-0.087* | *0.997* |
| | Final And Decreasing | **5.21** | **0.024*** | **-4.54** | **0.006**** | **-4.81** | **0.005**** | *-0.268* | *0.957* |
| | Latest Decreasing | **4.23** | **0.040*** | **-3.08** | **0.049*** | **-3.34** | **0.036*** | *-0.268* | *0.957* |
| Age-specific | Decreasing | **6.00** | **0.014*** | **-3.58** | **0.026*** | **-3.66** | **0.029*** | *-0.087* | *0.997* |
| | Final And Decreasing | **5.15** | **0.023*** | **-4.54** | **0.006**** | **-4.81** | **0.005**** | *-0.268* | *0.957* |
| | Latest Decreasing | **5.80** | **0.016*** | **-2.88** | **0.046*** | **-3.14** | **0.033*** | *-0.268* | *0.957* |

Wald testing confirmed the significance of Age effects in all cases. Significant pairwise differences were found between 5- and both 8- and 12mo data for all categories tested, aside from the decreasing FCPts identified in the cross-age ROI channels, suggesting the habituation process continues for longer at younger ages. Significance: p*<0.05; p**<0.01, p***<0.001. Age-specific: channels in which a significant hemodynamic response had been identified using conventional analyses; overall: cross-age set of channels in which a significant hemodynamic response had been identified using conventional analyses.

**(d) Figures analogous to those in the main manuscript, which examine the longitudinal differences between the other four types of FCPts, using cross-age channels.**

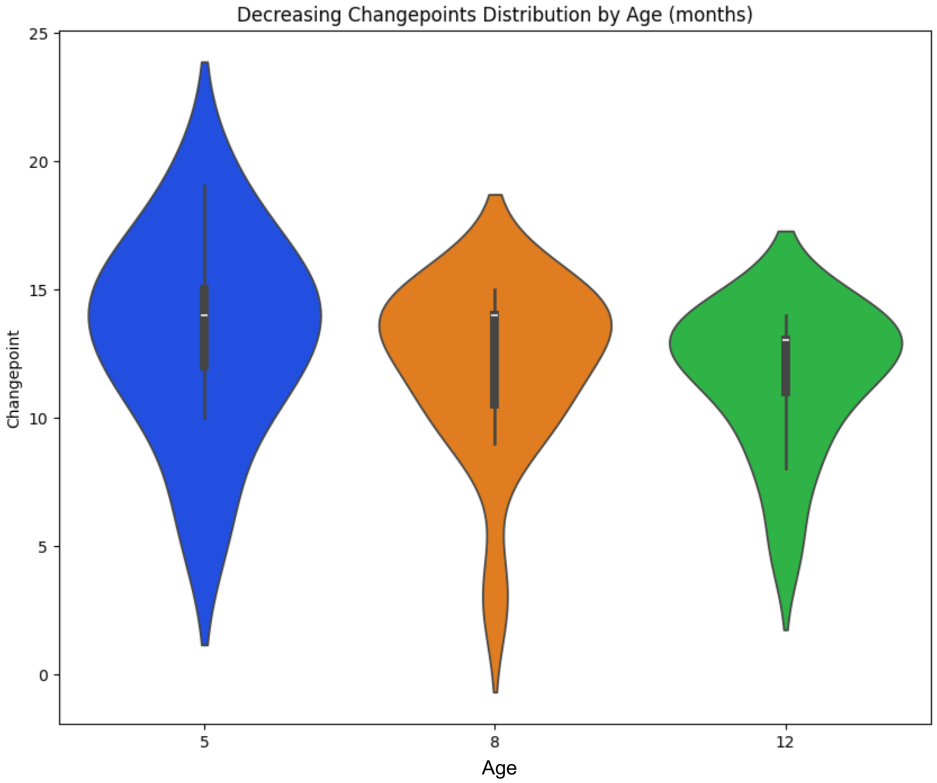


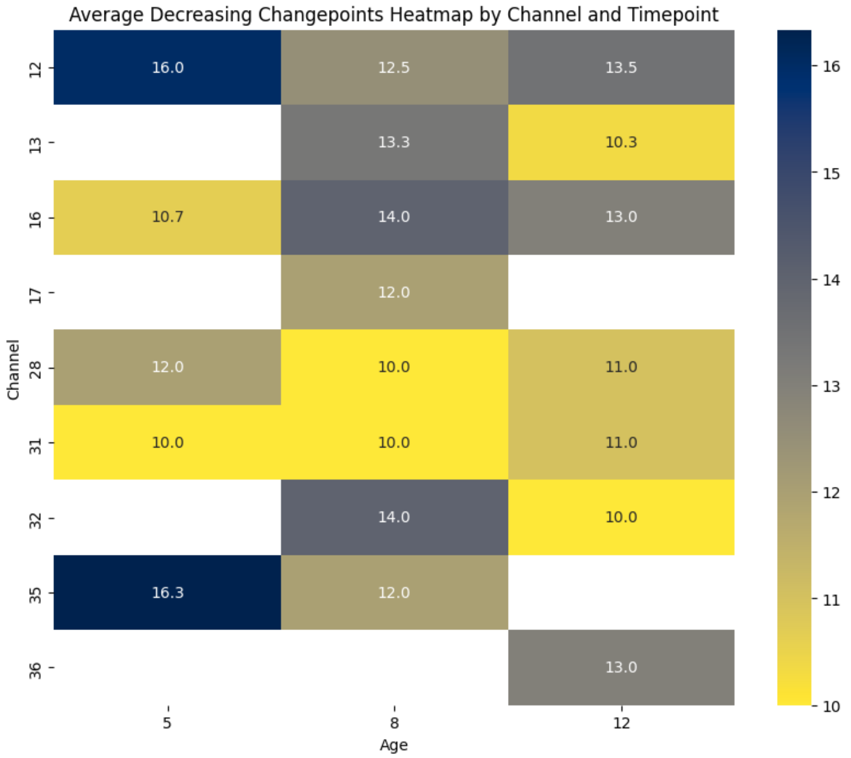

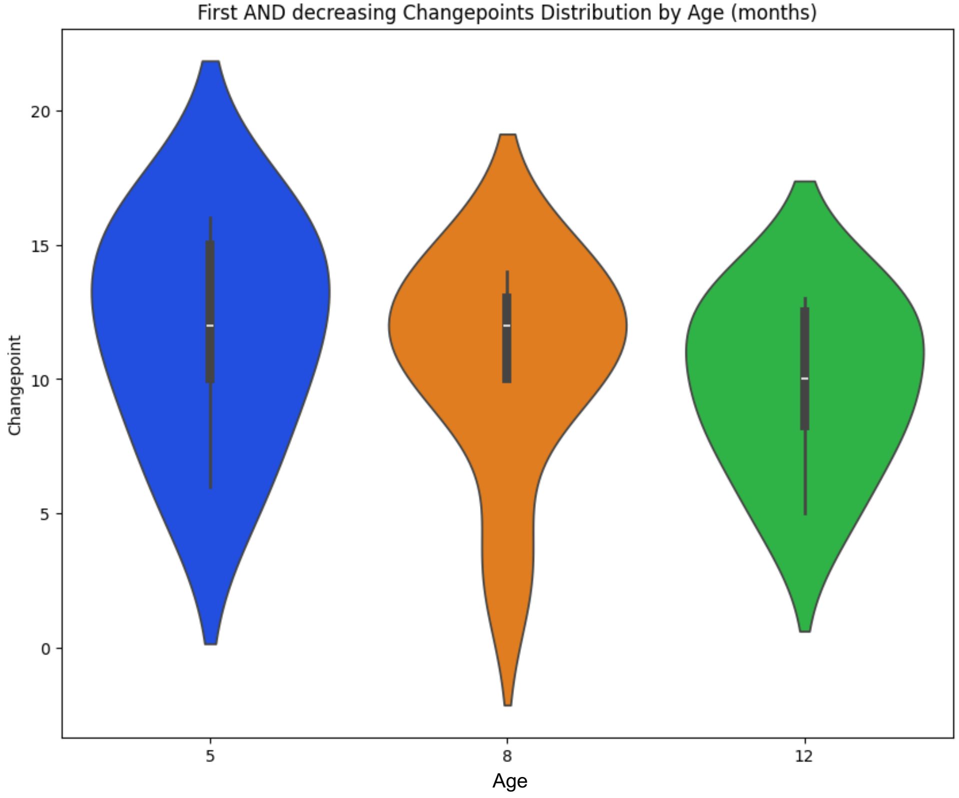
First AND decreasing Changepoints Distribution by Age (months)
Changepoint
Age
5
8
12
0
5
10
15
20

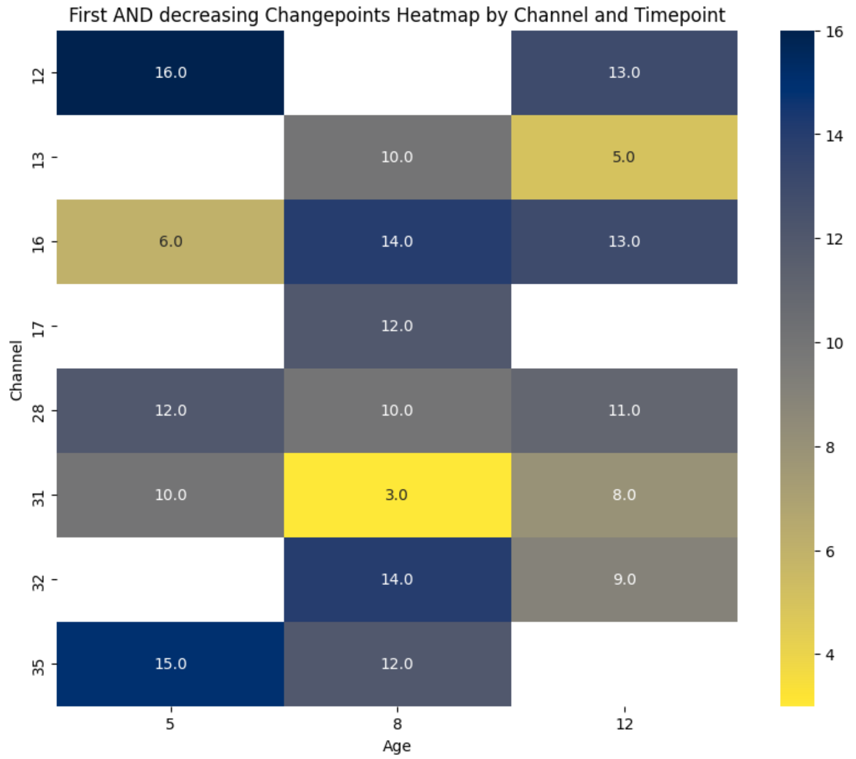
First AND decreasing Changepoints Heatmap by Channel and Timepoint
Channel
Age
12
13
16
17
28
31
32
35
5
8
12
16.0
13.0
10.0
5.0
6.0
14.0
13.0
12.0
12.0
10.0
11.0
10.0
3.0
8.0
14.0
9.0
15.0
12.0
4
6
8
10
12
14
16

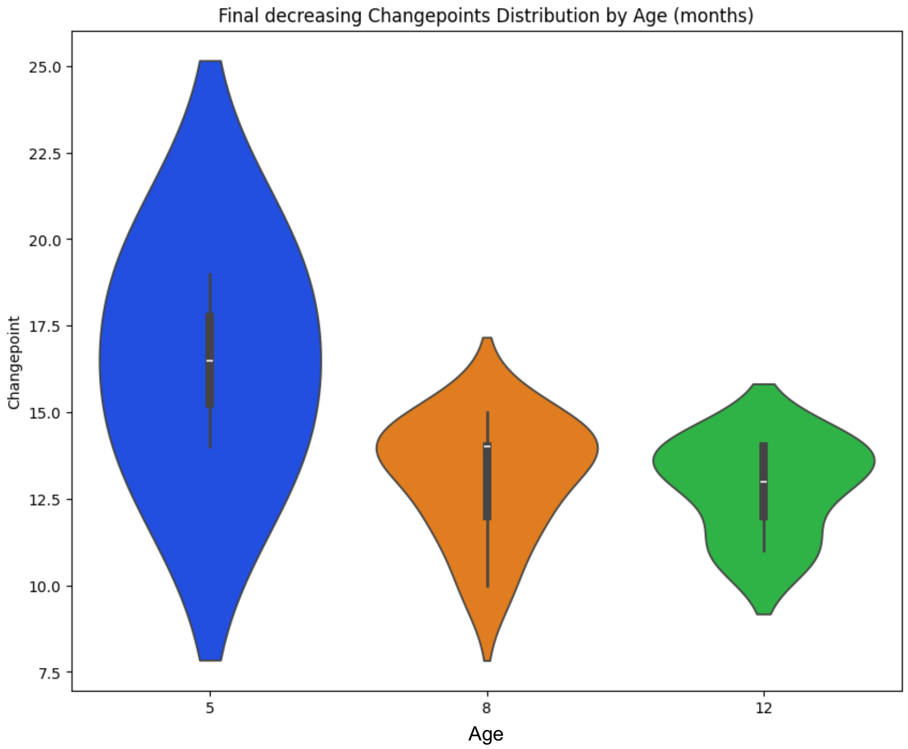
Final decreasing Changepoints Distribution by Age (months)
Changepoint
25.0
22.5
20.0
17.5
15.0
12.5
10.0
7.5
5
8
12
Age

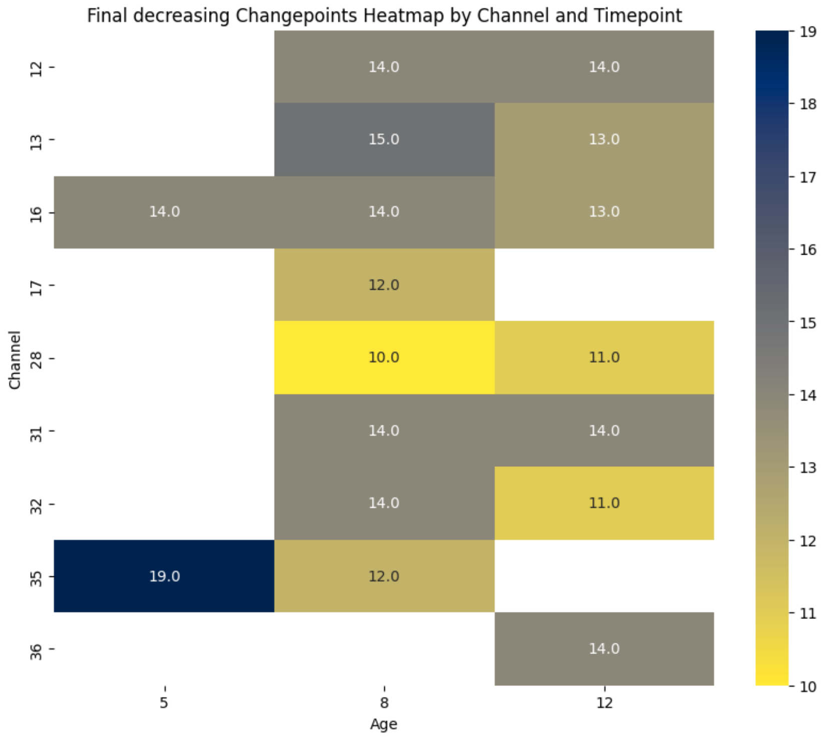
Final decreasing Changepoints Heatmap by Channel and Timepoint
Channel
12
13
16
17
28
31
32
35
36
14.0
14.0
15.0
13.0
14.0
14.0
13.0
12.0
10.0
11.0
14.0
14.0
14.0
11.0
19.0
12.0
14.0
5
8
12
Age
19
18
17
16
15
14
13
12
11
10

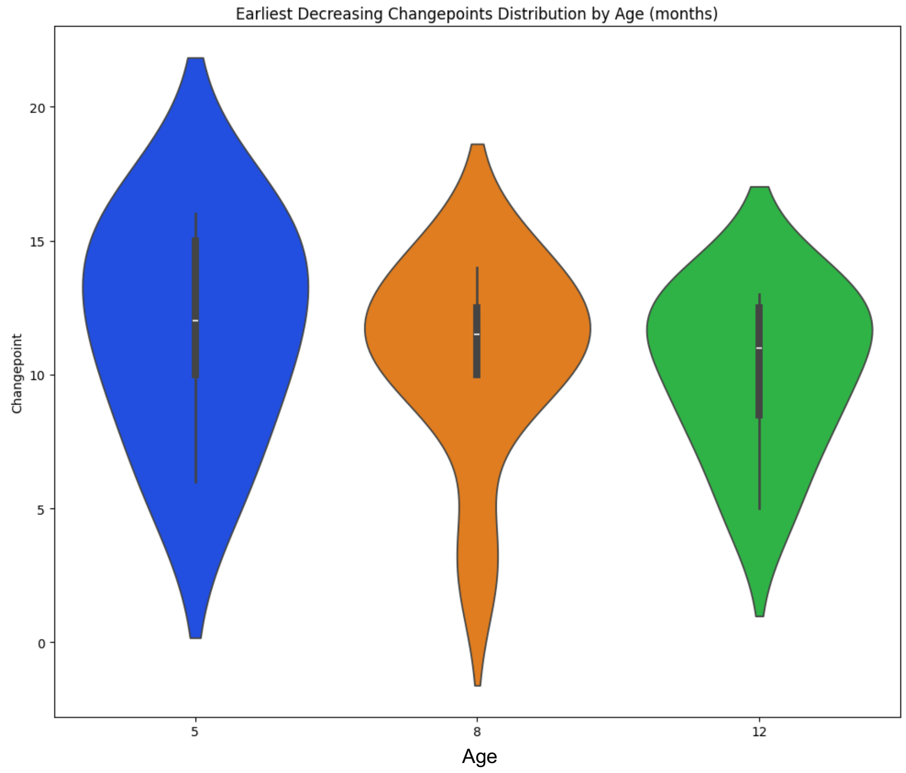
Earliest Decreasing Changepoints Distribution by Age (months)
Changepoint
0
5
10
15
20
5
8
12
Age

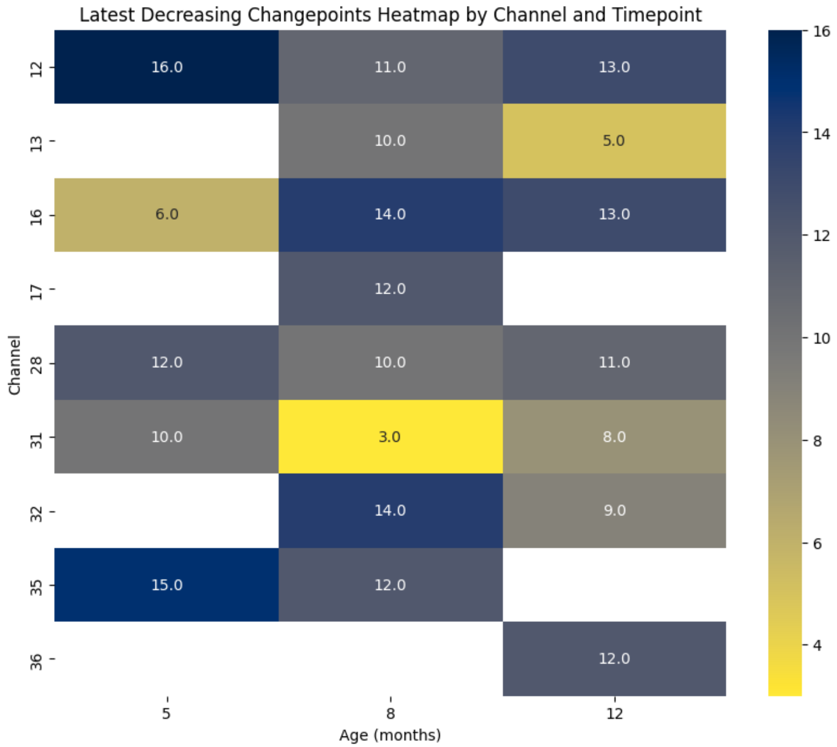
Latest Decreasing Changepoints Heatmap by Channel and Timepoint
Channel
12
13
16
17
28
31
32
35
36
16.0
11.0
13.0
10.0
5.0
6.0
14.0
13.0
12.0
12.0
10.0
11.0
10.0
3.0
8.0
14.0
9.0
15.0
12.0
12.0
5
8
12
Age (months)
16
14
12
10
8
6
4

### Supplementary Materials 8: FCPts and the mean surface between them

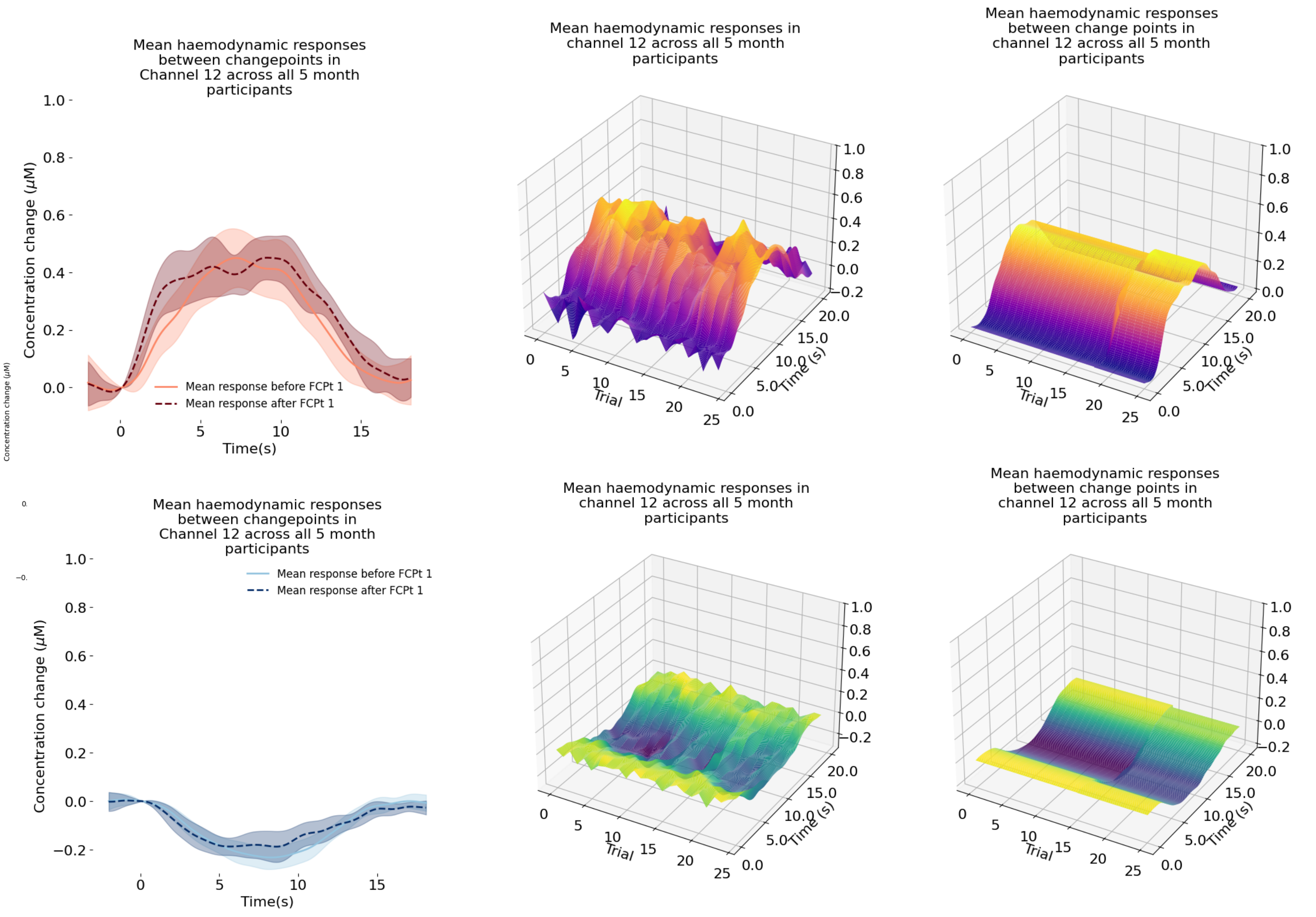

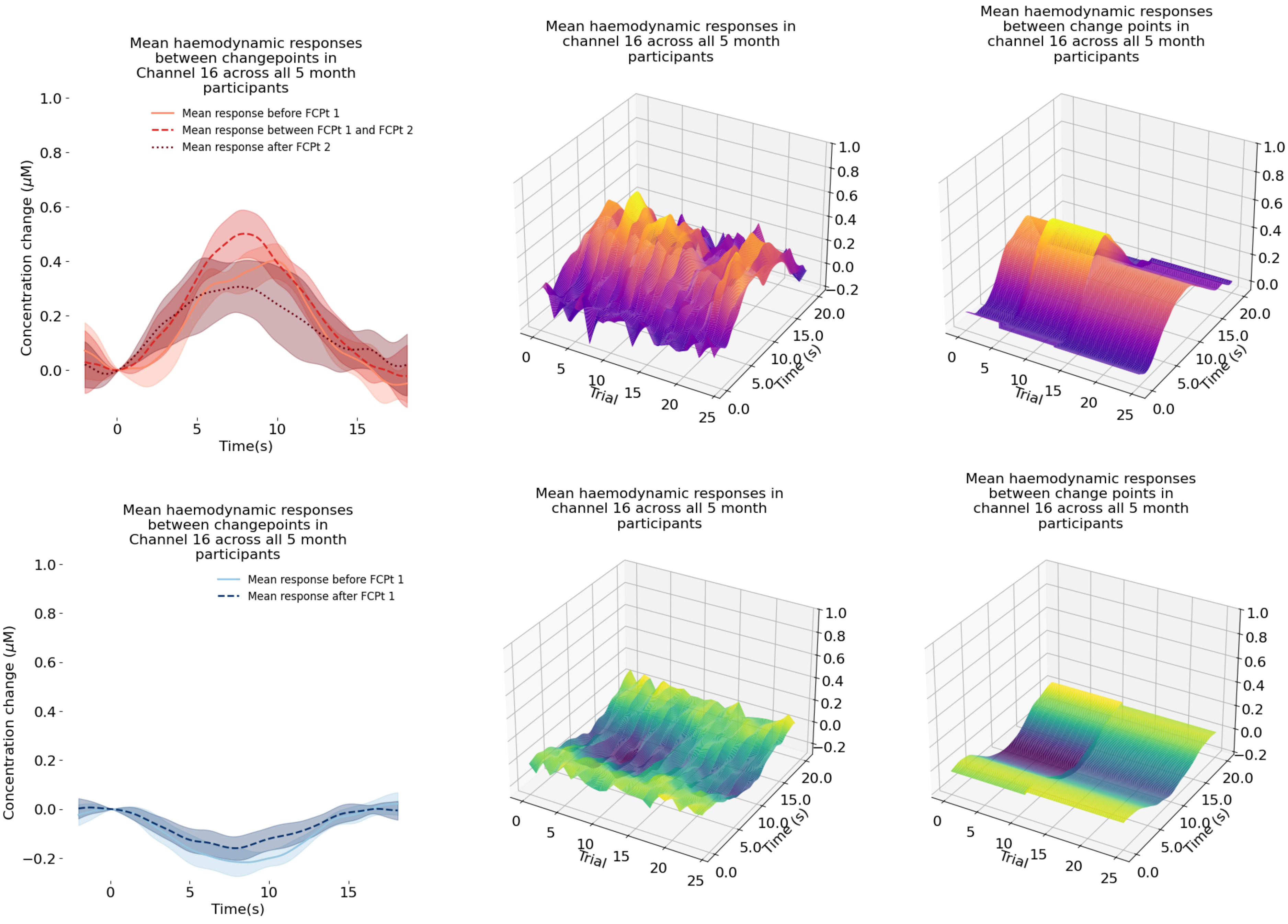
Mean haemodynamic responses between changepoints in Channel 16 across all 5 month participants
Mean response before FCPt 1
Mean response between FCPt 1 and FCPt 2
Mean response after FCPt 2
Concentration change (μM)
Time(s)
Mean haemodynamic responses in channel 16 across all 5 month participants
Trial
Time (s)
Mean haemodynamic responses between change points in channel 16 across all 5 month participants
Mean haemodynamic responses between changepoints in Channel 16 across all 5 month participants
Mean response before FCPt 1
Mean response after FCPt 1
Concentration change (μM)
Time(s)
Mean haemodynamic responses in channel 16 across all 5 month participants
Trial
Time (s)
Mean haemodynamic responses between change points in channel 16 across all 5 month participants

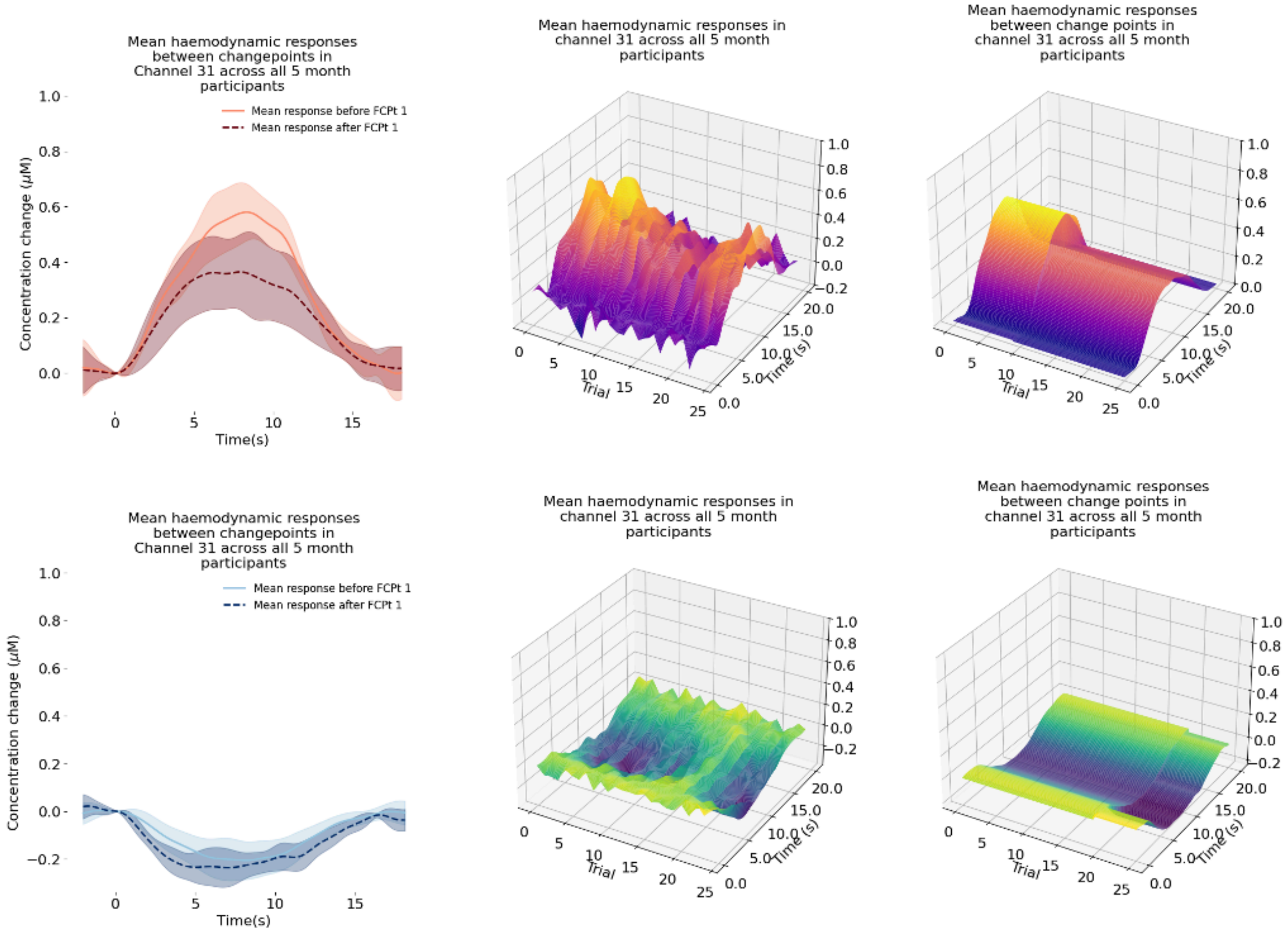
Mean haemodynamic responses between changepoints in Channel 31 across all 5 month participants
Mean response before FCPt 1
Mean response after FCPt 1
Concentration change (μM)
Time(s)
Mean haemodynamic responses in channel 31 across all 5 month participants
Trial
Time (s)
Mean haemodynamic responses between change points in channel 31 across all 5 month participants
Trial
Time (s)
Mean haemodynamic responses between changepoints in Channel 31 across all 5 month participants
Mean response before FCPt 1
Mean response after FCPt 1
Concentration change (μM)
Time(s)
Mean haemodynamic responses in channel 31 across all 5 month participants
Trial
Time (s)
Mean haemodynamic responses between change points in channel 31 across all 5 month participants
Trial
Time (s)

1702

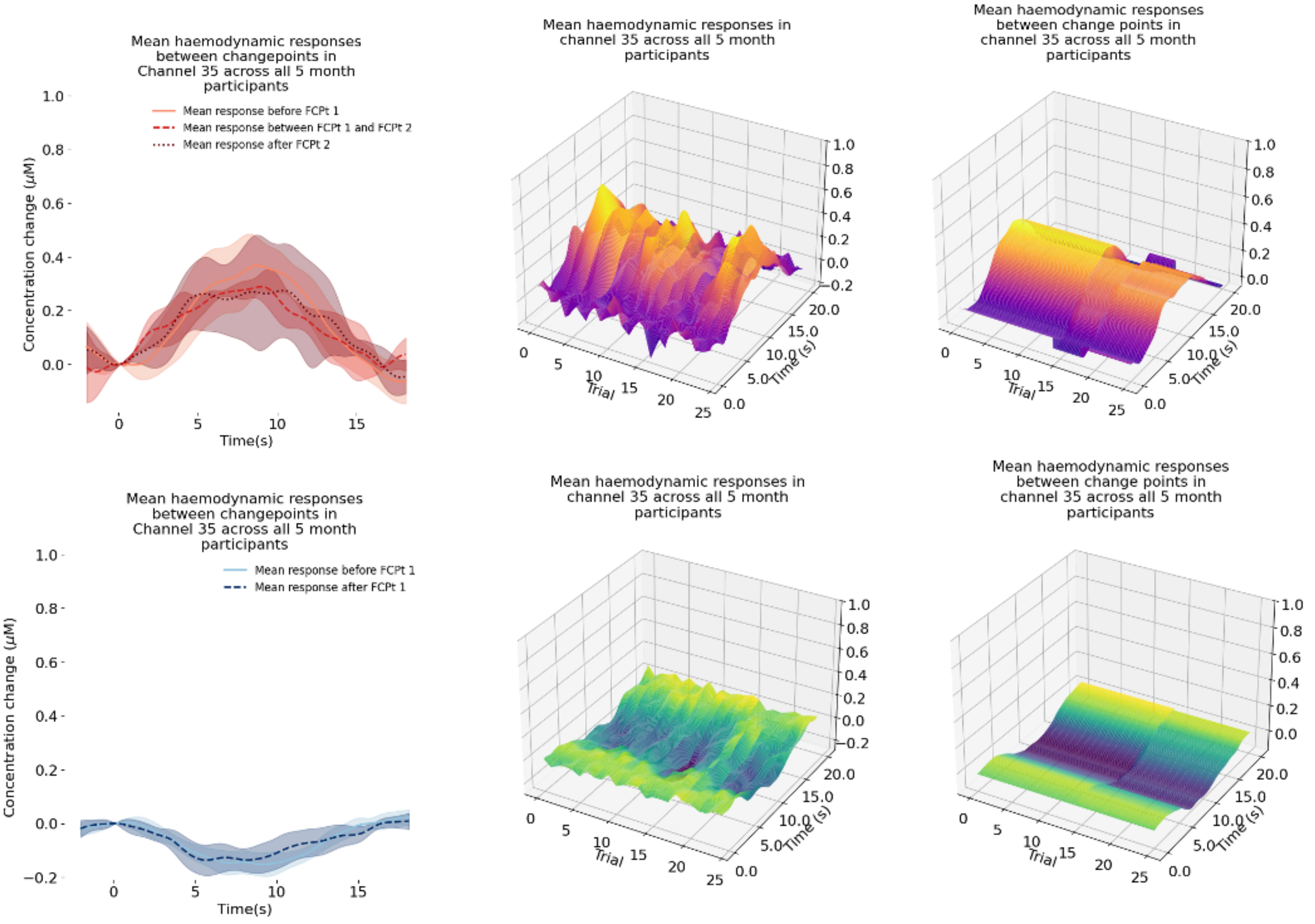

Mean haemodynamic responses between changepoints in Channel 35 across all 5 month participants
Mean response before FCPt 1
Mean response between FCPt 1 and FCPt 2
Mean response after FCPt 2
Concentration change (μM)
Time(s)
Mean haemodynamic responses in channel 35 across all 5 month participants
Trial
Time (s)
Mean haemodynamic responses between change points in channel 35 across all 5 month participants
Trial
Time (s)
Mean haemodynamic responses between changepoints in Channel 35 across all 5 month participants
Mean response before FCPt 1
Mean response after FCPt 1
Concentration change (μM)
Time(s)
Mean haemodynamic responses in channel 35 across all 5 month participants
Trial
Time (s)
Mean haemodynamic responses between change points in channel 35 across all 5 month participants
Trial
Time (s)

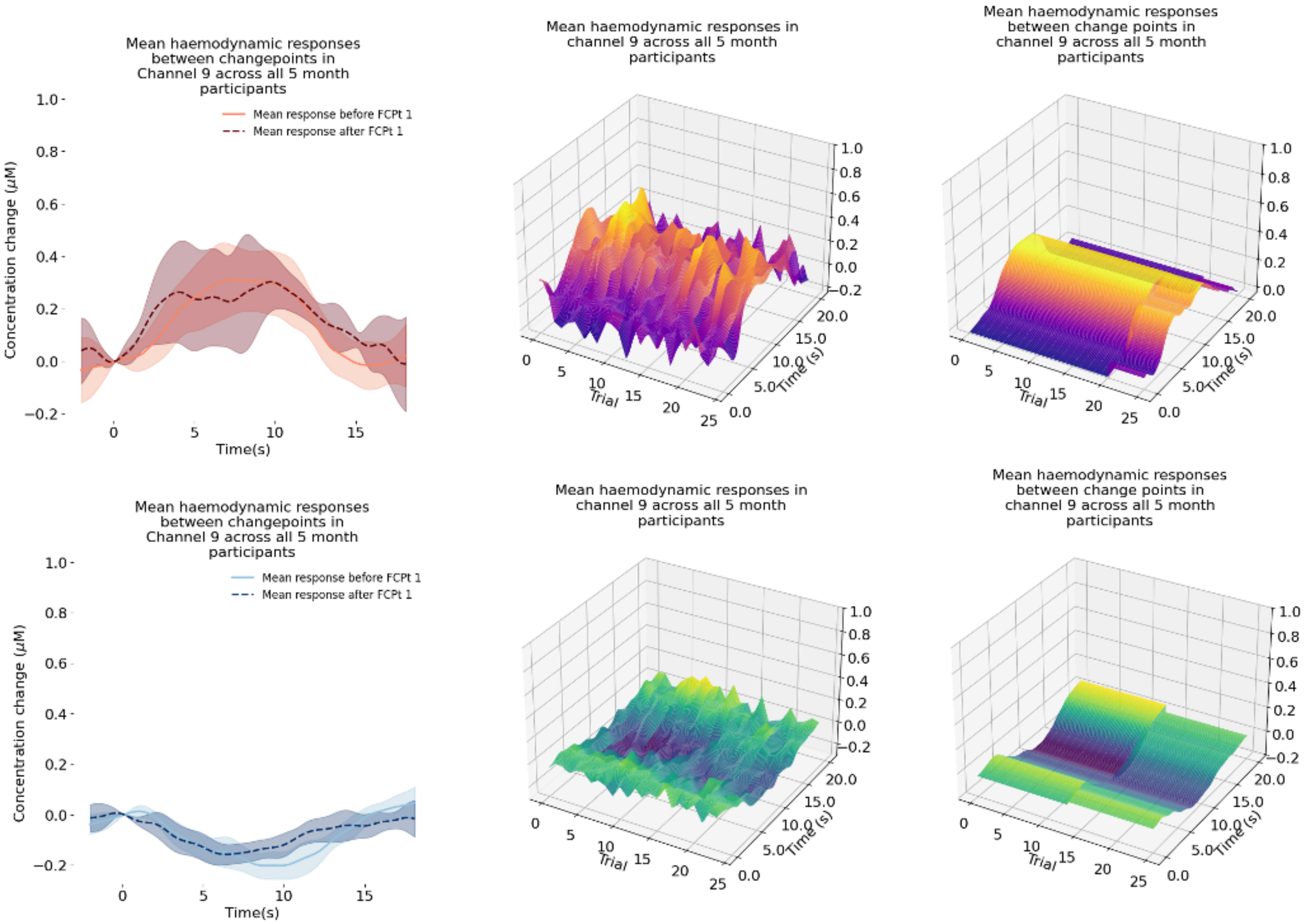
Mean haemodynamic responses between changepoints in Channel 9 across all 5 month participants
Mean response before FCPt 1
Mean response after FCPt 1
Concentration change (μM)
Time(s)
Mean haemodynamic responses in channel 9 across all 5 month participants
Trial
Time (s)
Mean haemodynamic responses between change points in channel 9 across all 5 month participants
Mean haemodynamic responses between changepoints in Channel 9 across all 5 month participants
Mean response before FCPt 1
Mean response after FCPt 1
Concentration change (μM)
Time(s)
Mean haemodynamic responses in channel 9 across all 5 month participants
Mean haemodynamic responses between change points in channel 9 across all 5 month participants

1703

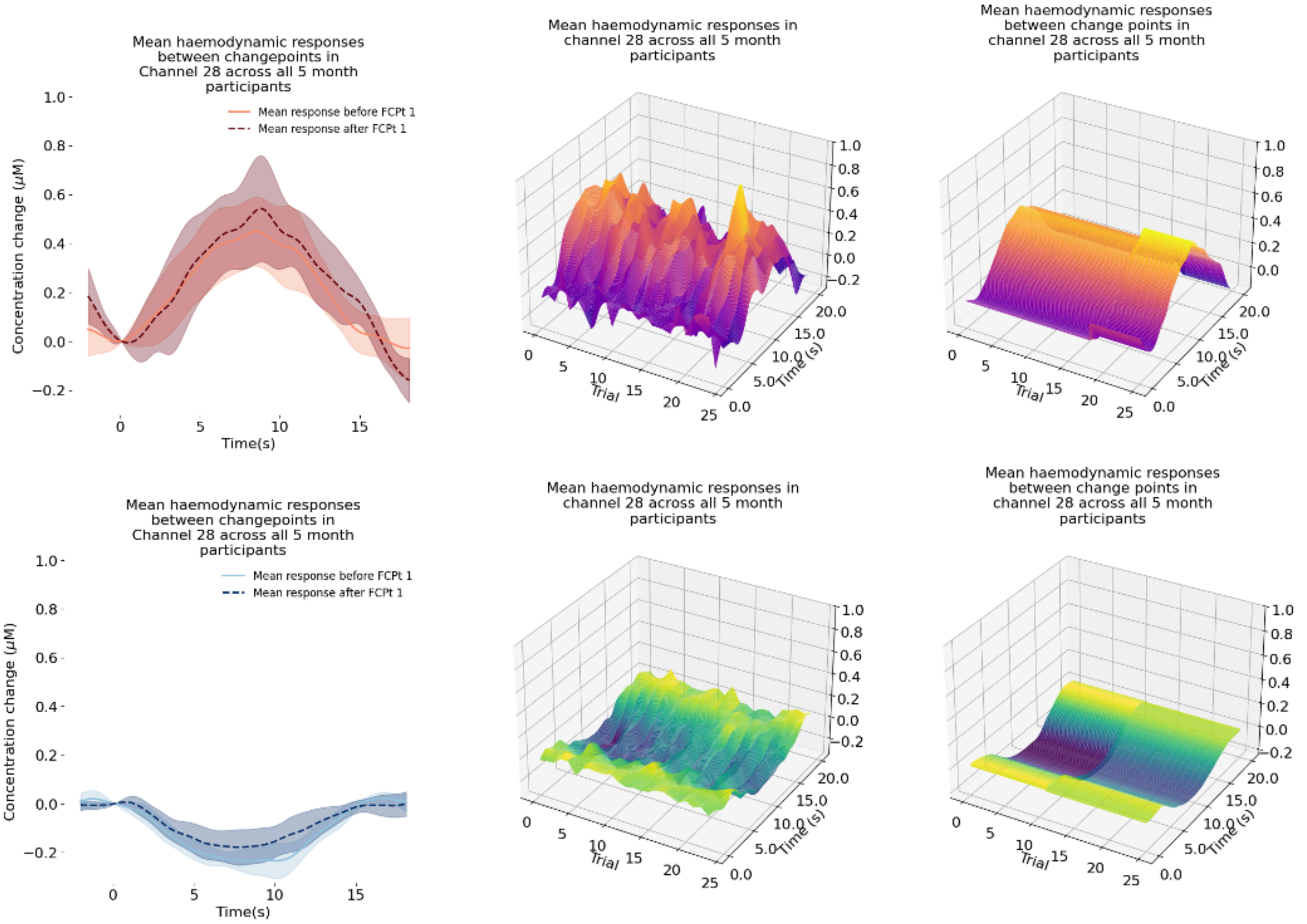
Mean haemodynamic responses between changepoints in Channel 28 across all 5 month participants
Mean response before FCPt 1
Mean response after FCPt 1
Concentration change (μM)
Time(s)
Mean haemodynamic responses in channel 28 across all 5 month participants
Trial
Time (s)
Mean haemodynamic responses between change points in channel 28 across all 5 month participants
Trial
Time (s)
Mean haemodynamic responses between changepoints in Channel 28 across all 5 month participants
Mean response before FCPt 1
Mean response after FCPt 1
Concentration change (μM)
Time(s)
Mean haemodynamic responses in channel 28 across all 5 month participants
Trial
Time (s)
Mean haemodynamic responses between change points in channel 28 across all 5 month participants
Trial
Time (s)

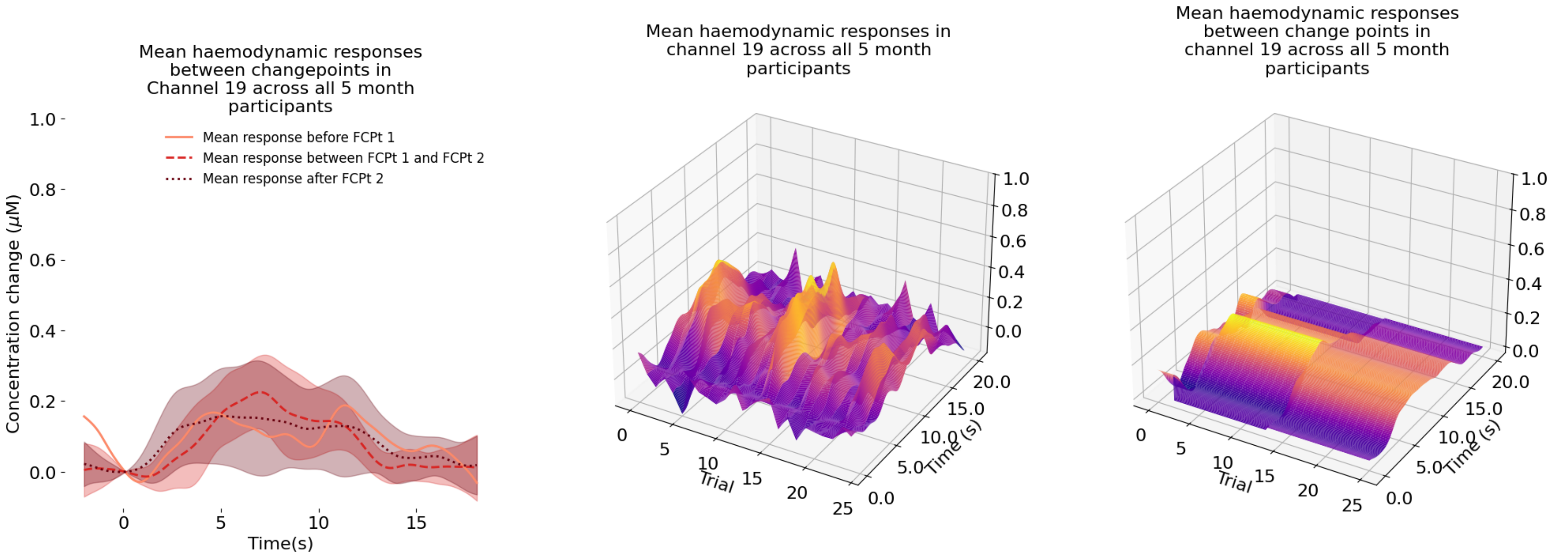
Mean haemodynamic responses between changepoints in Channel 19 across all 5 month participants
Mean response before FCPt 1
Mean response between FCPt 1 and FCPt 2
Mean response after FCPt 2
Concentration change (μM)
Time(s)
Mean haemodynamic responses in channel 19 across all 5 month participants
Trial
Time (s)
Mean haemodynamic responses between change points in channel 19 across all 5 month participants
Trial
Time (s)

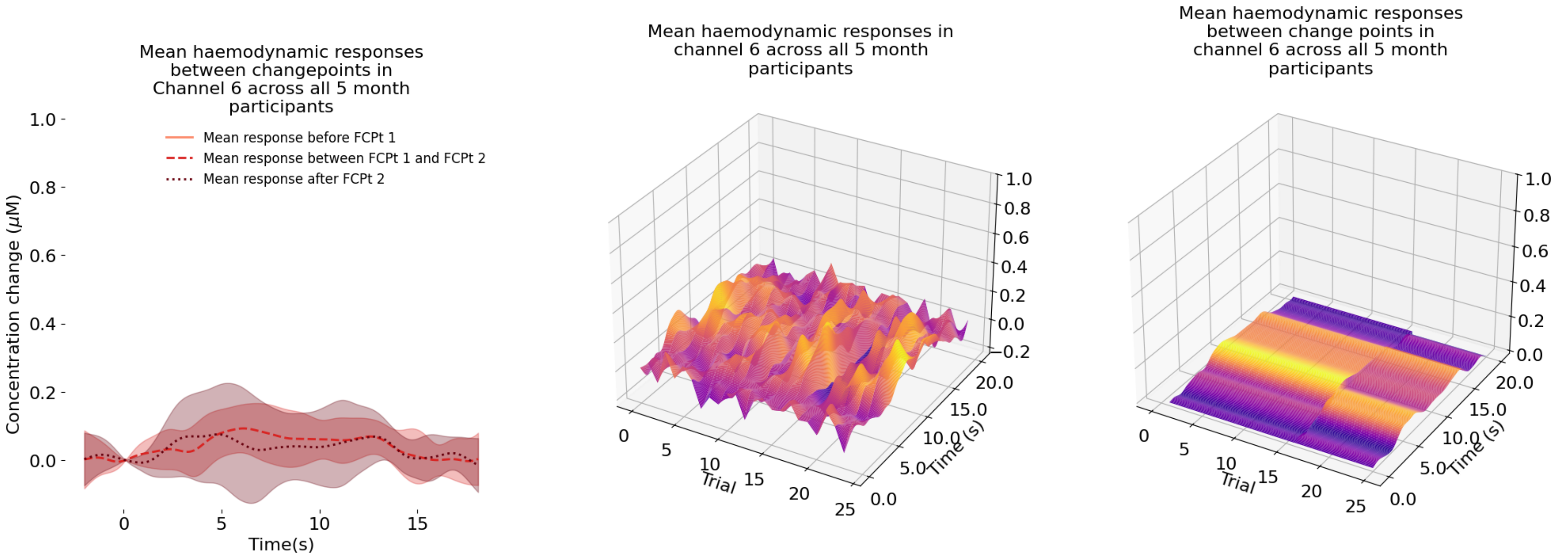
Mean haemodynamic responses between changepoints in Channel 6 across all 5 month participants
Mean response before FCPt 1
Mean response between FCPt 1 and FCPt 2
Mean response after FCPt 2
Concentration change (μM)
Time(s)
Mean haemodynamic responses in channel 6 across all 5 month participants
Trial
Time (s)
Mean haemodynamic responses between change points in channel 6 across all 5 month participants
Trial
Time (s)

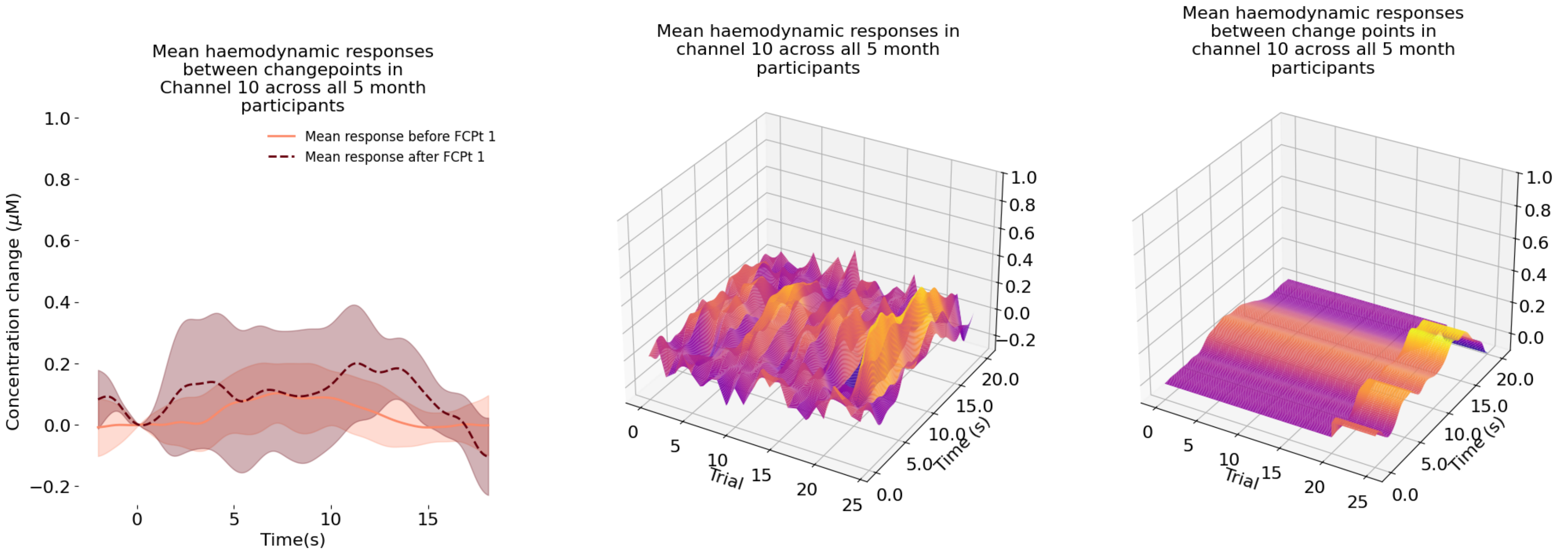
Mean haemodynamic responses between changepoints in Channel 10 across all 5 month participants
Mean response before FCPt 1
Mean response after FCPt 1
Concentration change (μM)
Time(s)
Mean haemodynamic responses in channel 10 across all 5 month participants
Trial
Time (s)
Mean haemodynamic responses between change points in channel 10 across all 5 month participants
Trial
Time (s)

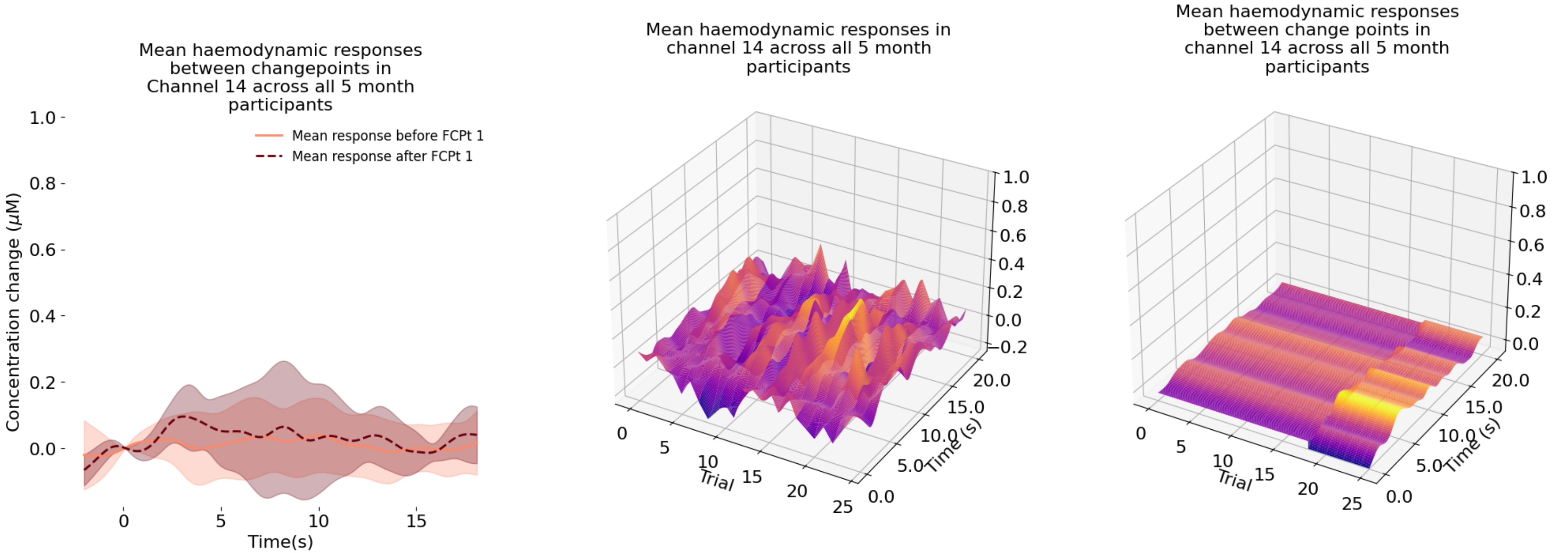
Mean haemodynamic responses between changepoints in Channel 14 across all 5 month participants
Mean response before FCPt 1
Mean response after FCPt 1
Concentration change (μM)
Time(s)
Mean haemodynamic responses in channel 14 across all 5 month participants
Trial
Time (s)
Mean haemodynamic responses between change points in channel 14 across all 5 month participants
Trial
Time (s)

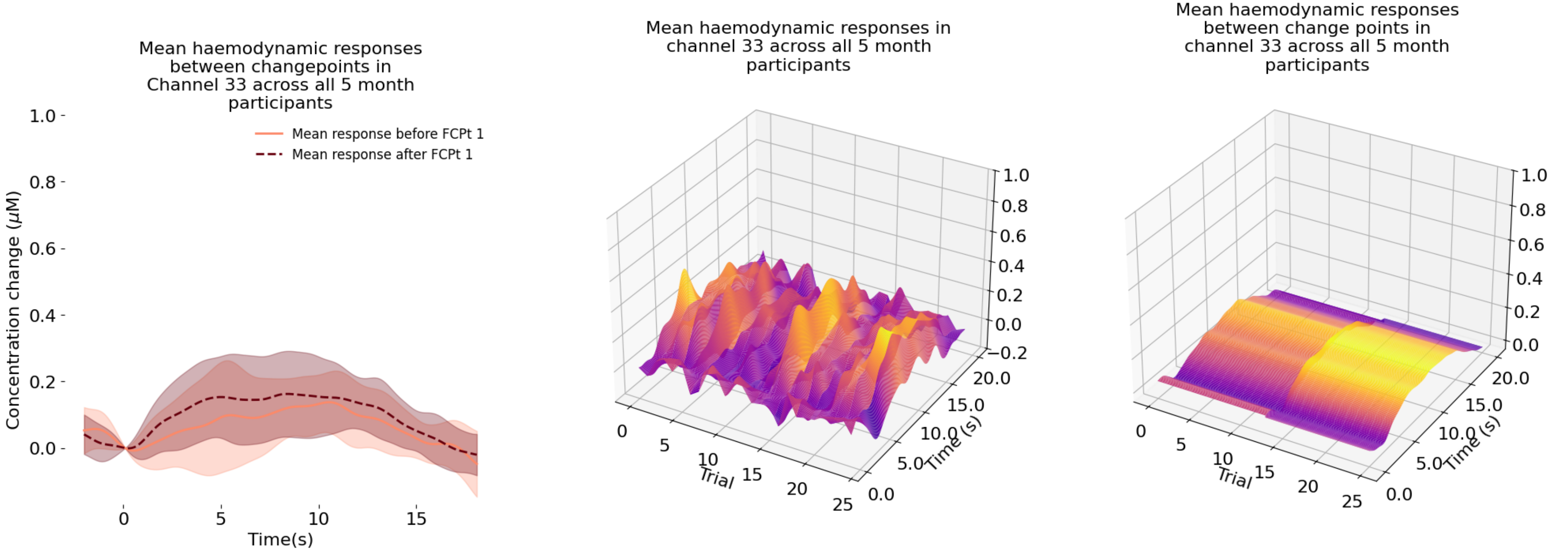
Mean haemodynamic responses between changepoints in Channel 33 across all 5 month participants
Mean response before FCPt 1
Mean response after FCPt 1
Concentration change (μM)
Time(s)
Mean haemodynamic responses in channel 33 across all 5 month participants
Trial
Time (s)
Mean haemodynamic responses between change points in channel 33 across all 5 month participants
Trial
Time (s)

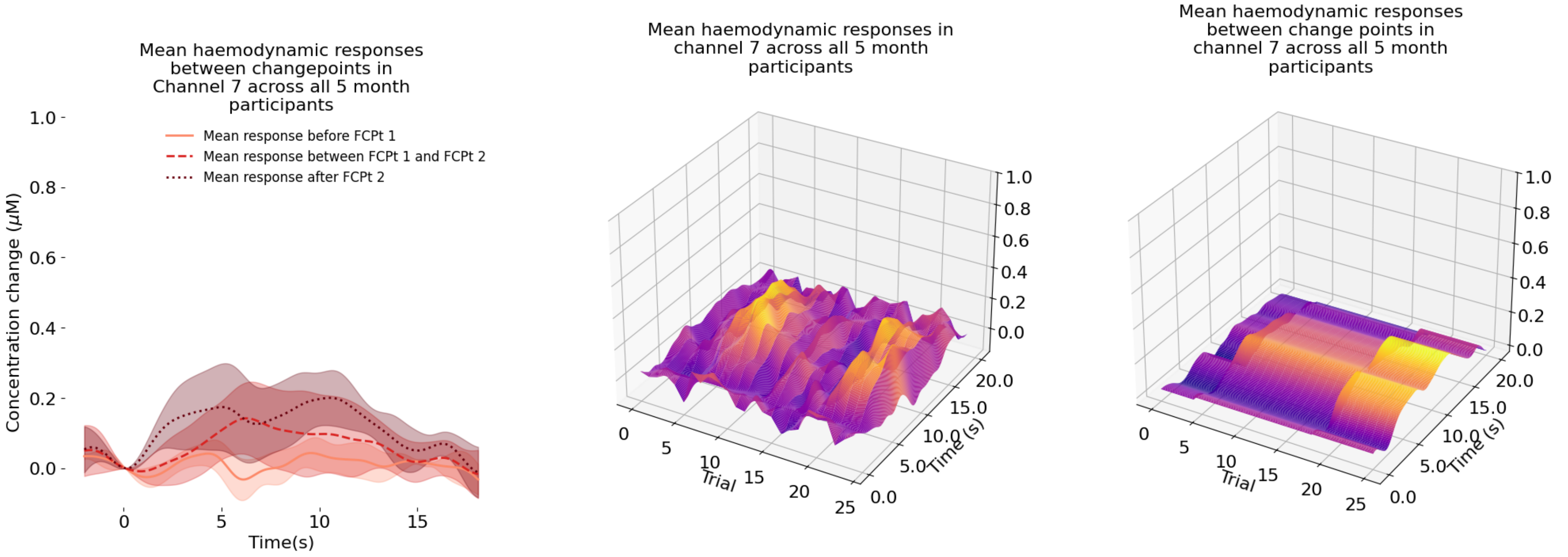
Mean haemodynamic responses between changepoints in Channel 7 across all 5 month participants
Mean response before FCPt 1
Mean response between FCPt 1 and FCPt 2
Mean response after FCPt 2
Concentration change (μM)
Time(s)
Mean haemodynamic responses in channel 7 across all 5 month participants
Trial
Time (s)
Mean haemodynamic responses between change points in channel 7 across all 5 month participants
Trial
Time (s)

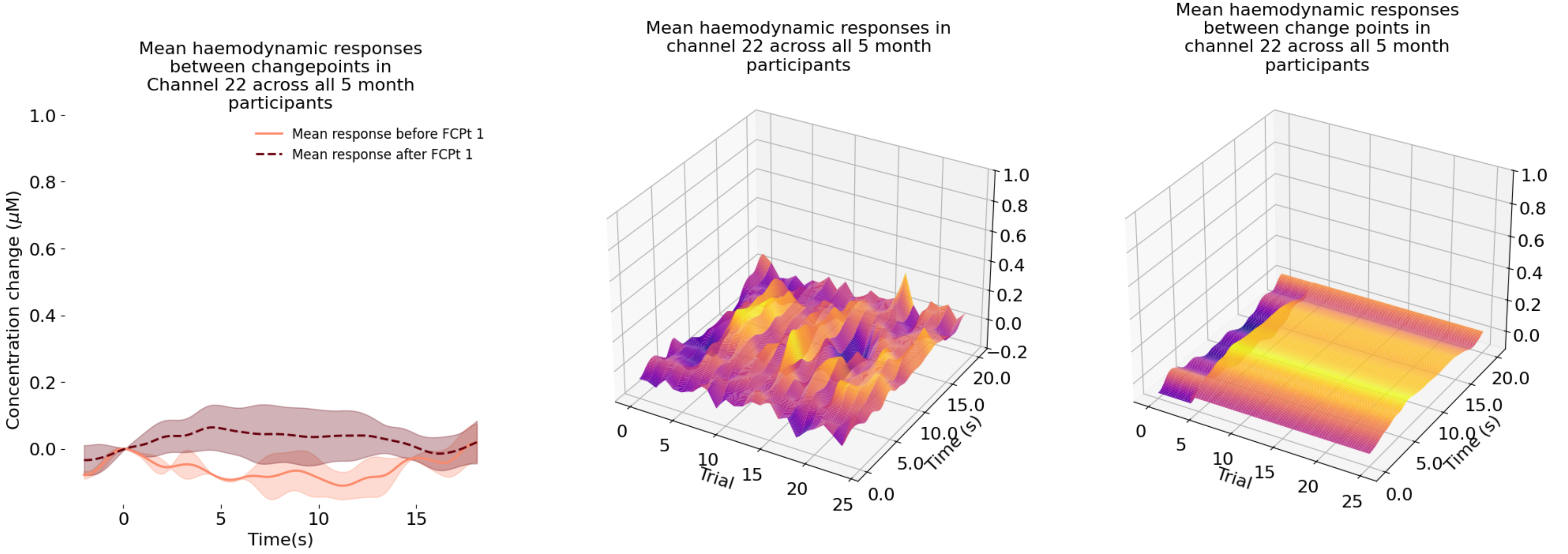
Mean haemodynamic responses between changepoints in Channel 22 across all 5 month participants
Mean response before FCPt 1
Mean response after FCPt 1
Concentration change (μM)
Time(s)
Mean haemodynamic responses in channel 22 across all 5 month participants
Trial
Time (s)
Mean haemodynamic responses between change points in channel 22 across all 5 month participants
Trial
Time (s)

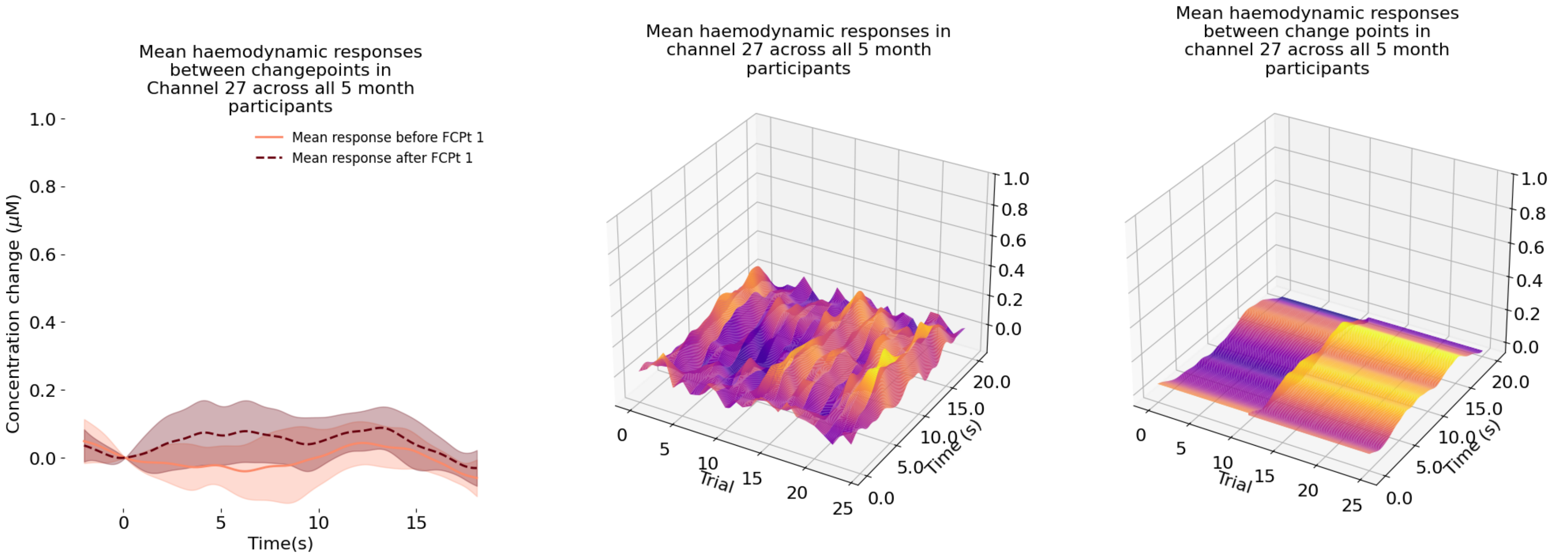
Mean haemodynamic responses between changepoints in Channel 27 across all 5 month participants
Mean response before FCPt 1
Mean response after FCPt 1
Concentration change (μM)
Time(s)
Mean haemodynamic responses in channel 27 across all 5 month participants
Trial
Time (s)
Mean haemodynamic responses between change points in channel 27 across all 5 month participants
Trial
Time (s)

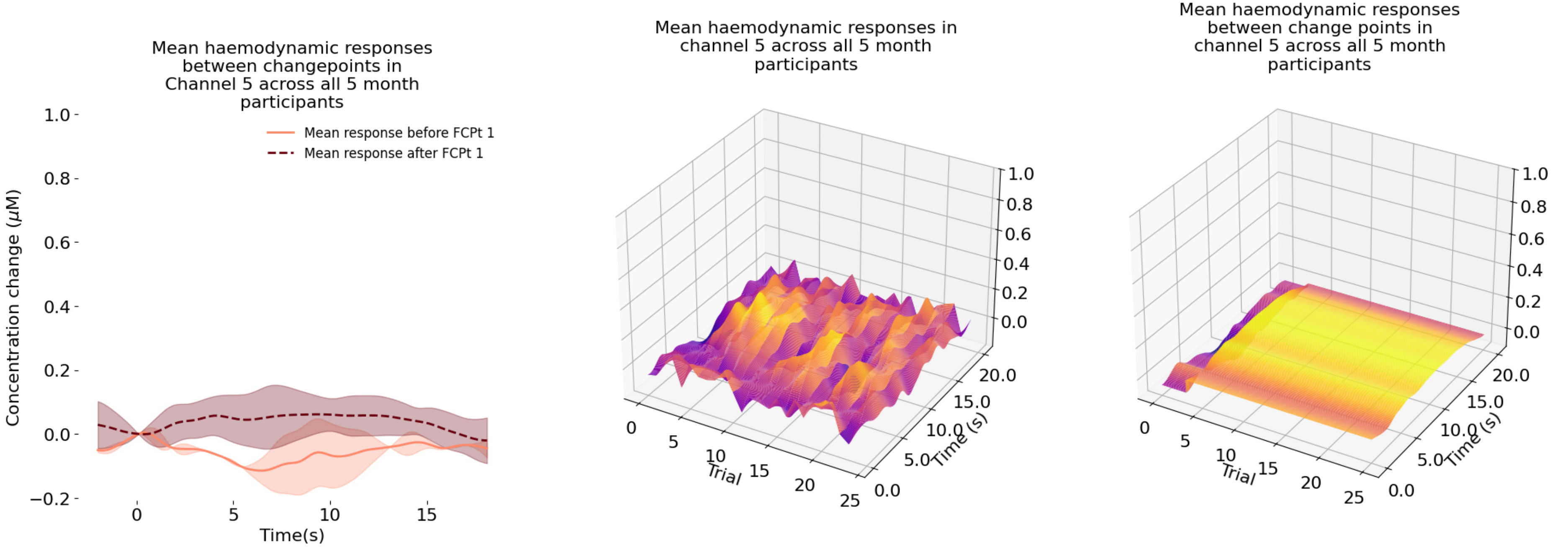
Mean haemodynamic responses between changepoints in Channel 5 across all 5 month participants
Mean response before FCPt 1
Mean response after FCPt 1
Concentration change (μM)
Time(s)
Mean haemodynamic responses in channel 5 across all 5 month participants
Trial
Time (s)
Mean haemodynamic responses between change points in channel 5 across all 5 month participants
Trial
Time (s)

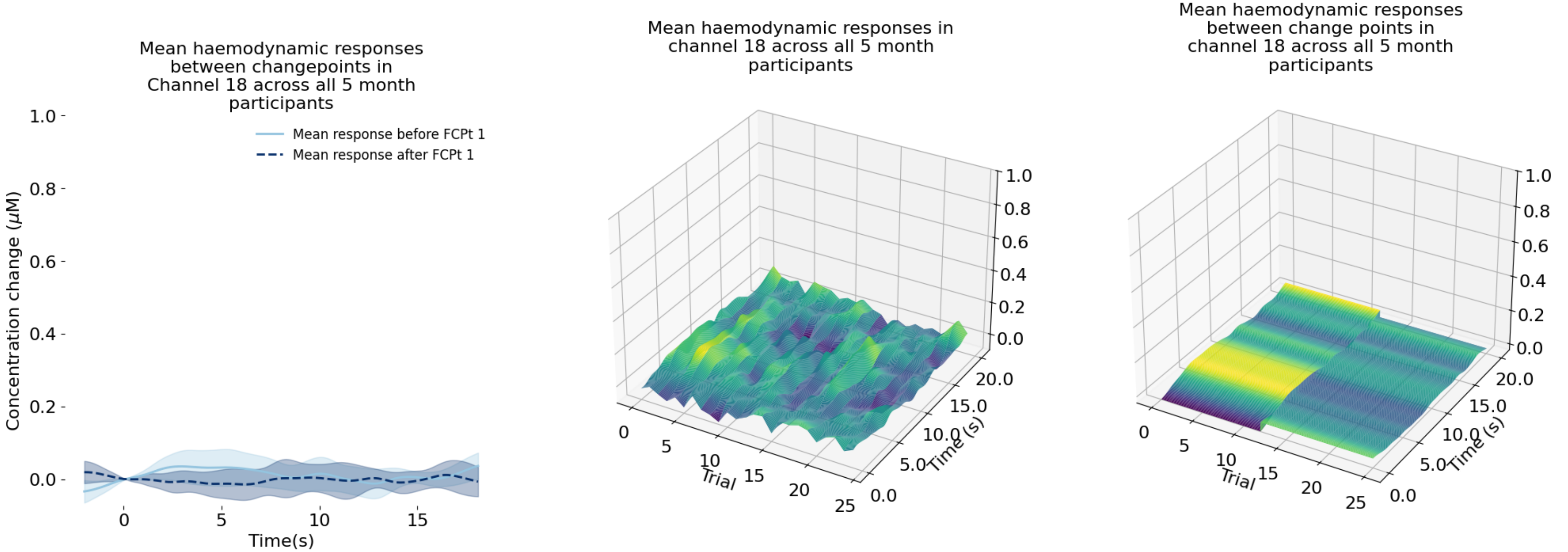
Mean haemodynamic responses between changepoints in Channel 18 across all 5 month participants
Mean response before FCPt 1
Mean response after FCPt 1
Concentration change (μM)
Time(s)
Mean haemodynamic responses in channel 18 across all 5 month participants
Trial
Time (s)
Mean haemodynamic responses between change points in channel 18 across all 5 month participants
Trial
Time (s)

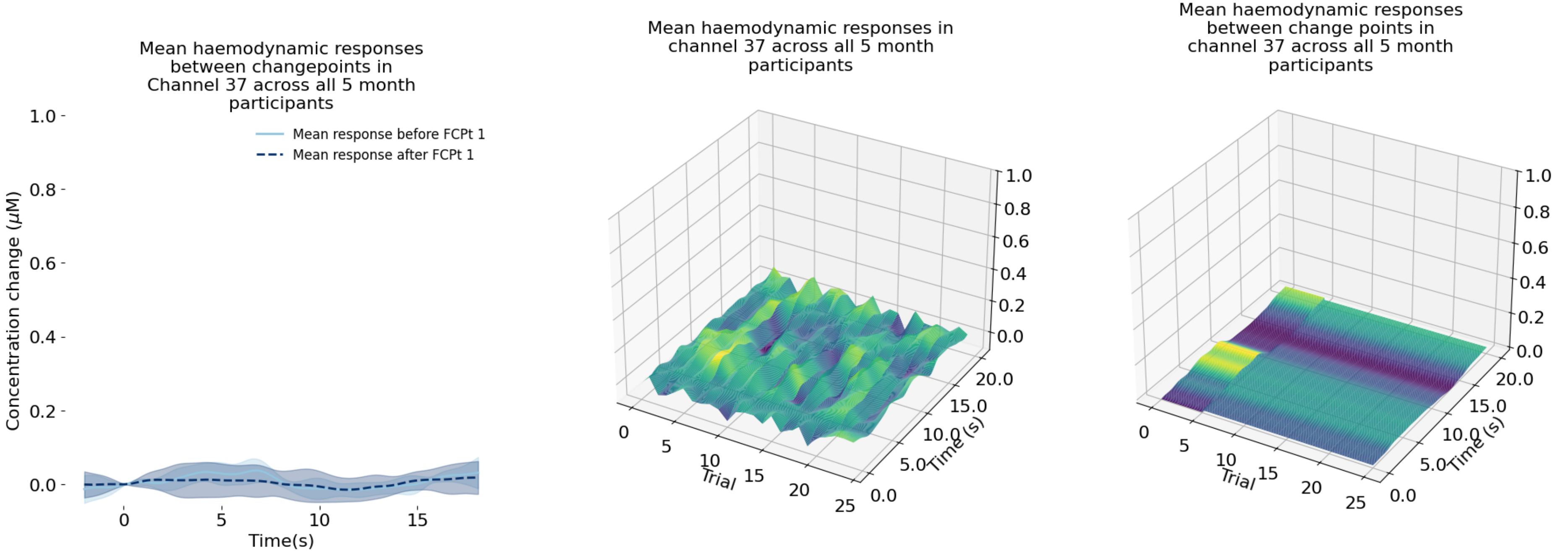
Mean haemodynamic responses between changepoints in Channel 37 across all 5 month participants
Mean response before FCPt 1
Mean response after FCPt 1
Concentration change ($\mu$M)
Time(s)
Mean haemodynamic responses in channel 37 across all 5 month participants
Trial
Time (s)
Mean haemodynamic responses between change points in channel 37 across all 5 month participants
Trial
Time (s)

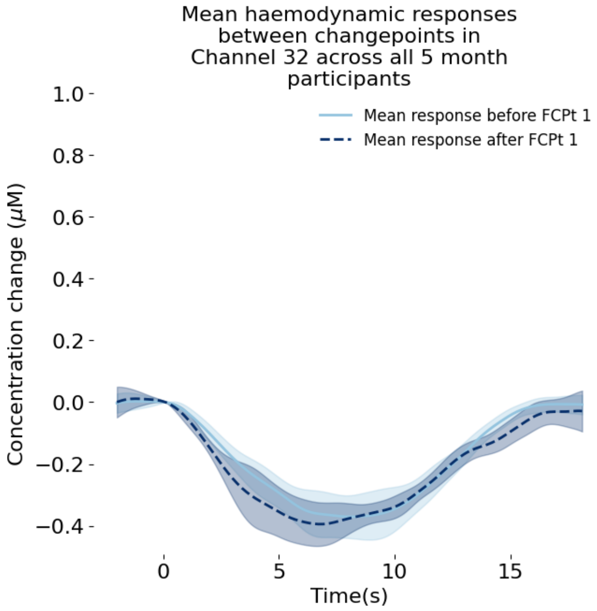
Mean haemodynamic responses between changepoints in Channel 32 across all 5 month participants
Mean response before FCPt 1
Mean response after FCPt 1
Concentration change (μM)
1.0
0.8
0.6
0.4
0.2
0.0
−0.2
−0.4
0
5
10
15
Time(s)

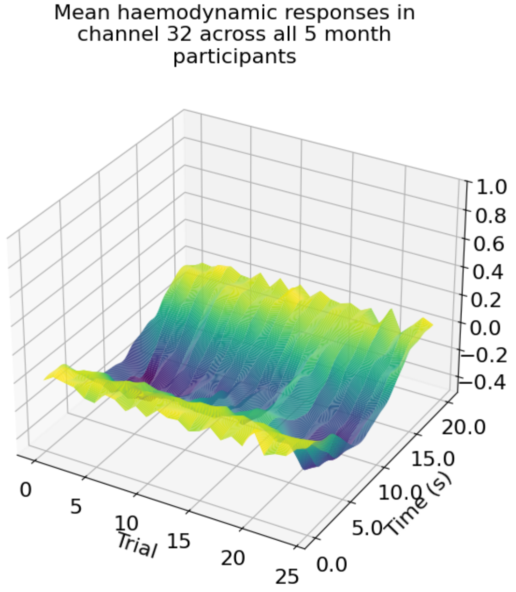
Mean haemodynamic responses in channel 32 across all 5 month participants
1.0
0.8
0.6
0.4
0.2
0.0
−0.2
−0.4
0
5
10
15
20
25
Trial
0.0
5.0
10.0
15.0
20.0
Time (s)

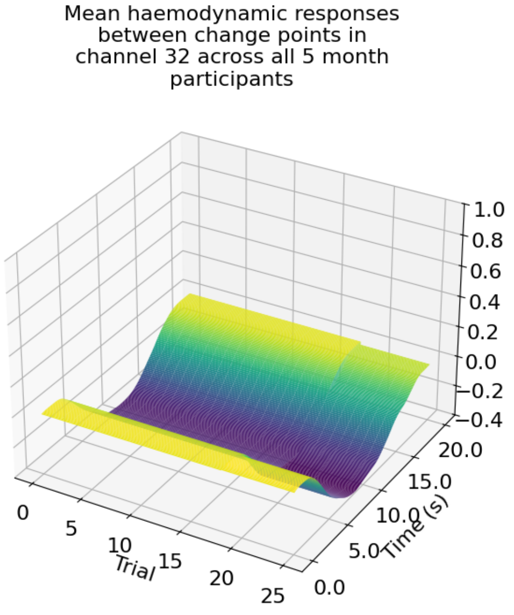
Mean haemodynamic responses between change points in channel 32 across all 5 month participants
1.0
0.8
0.6
0.4
0.2
0.0
−0.2
−0.4
0
5
10
15
20
25
Trial
0.0
5.0
10.0
15.0
20.0
Time (s)

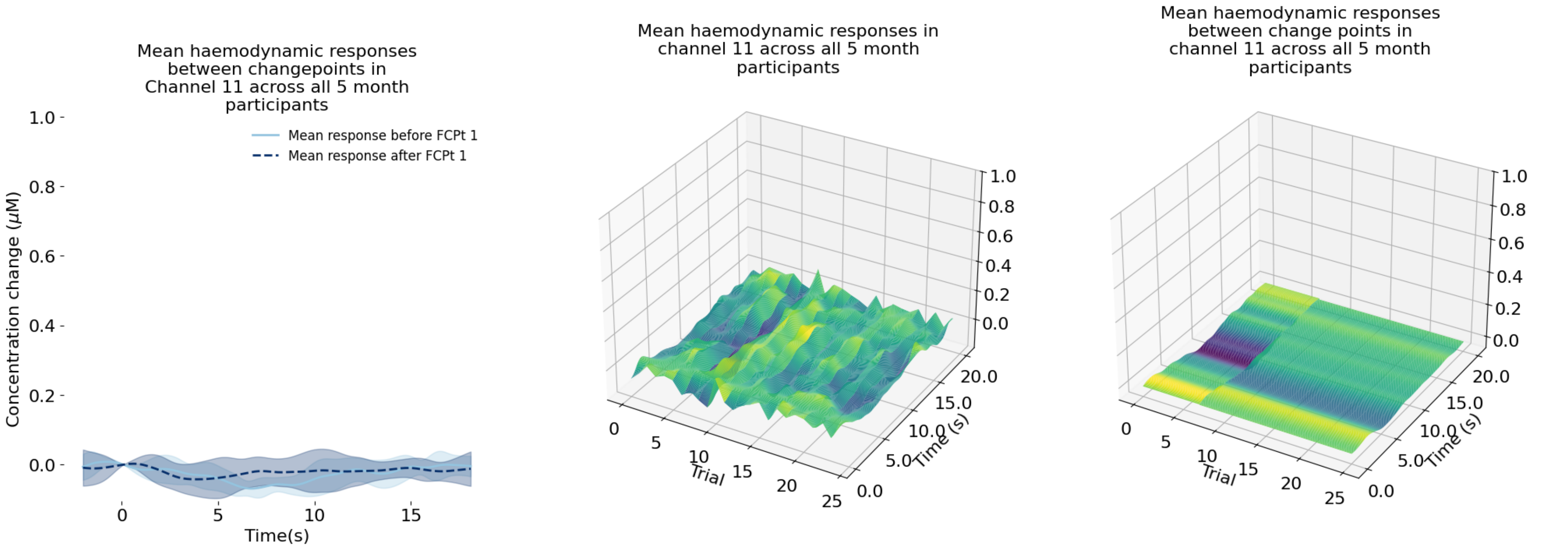
Mean haemodynamic responses between changepoints in Channel 11 across all 5 month participants
Mean response before FCPt 1
Mean response after FCPt 1
Concentration change (μM)
Time(s)
Mean haemodynamic responses in channel 11 across all 5 month participants
Trial
Time (s)
Mean haemodynamic responses between change points in channel 11 across all 5 month participants
Trial
Time (s)

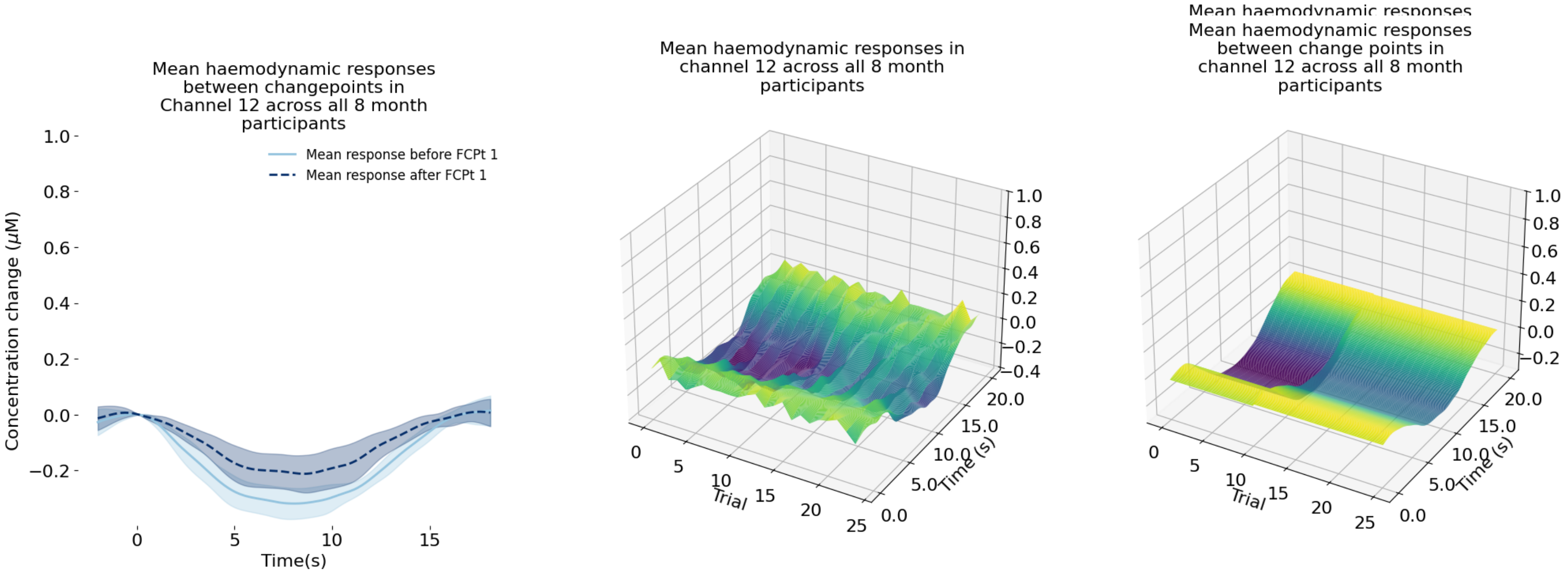
Mean haemodynamic responses between changepoints in Channel 12 across all 8 month participants
Mean response before FCPt 1
Mean response after FCPt 1
Concentration change (μM)
Time(s)
Mean haemodynamic responses in channel 12 across all 8 month participants
Trial
Time (s)
Mean haemodynamic responses between change points in channel 12 across all 8 month participants
Trial
Time (s)

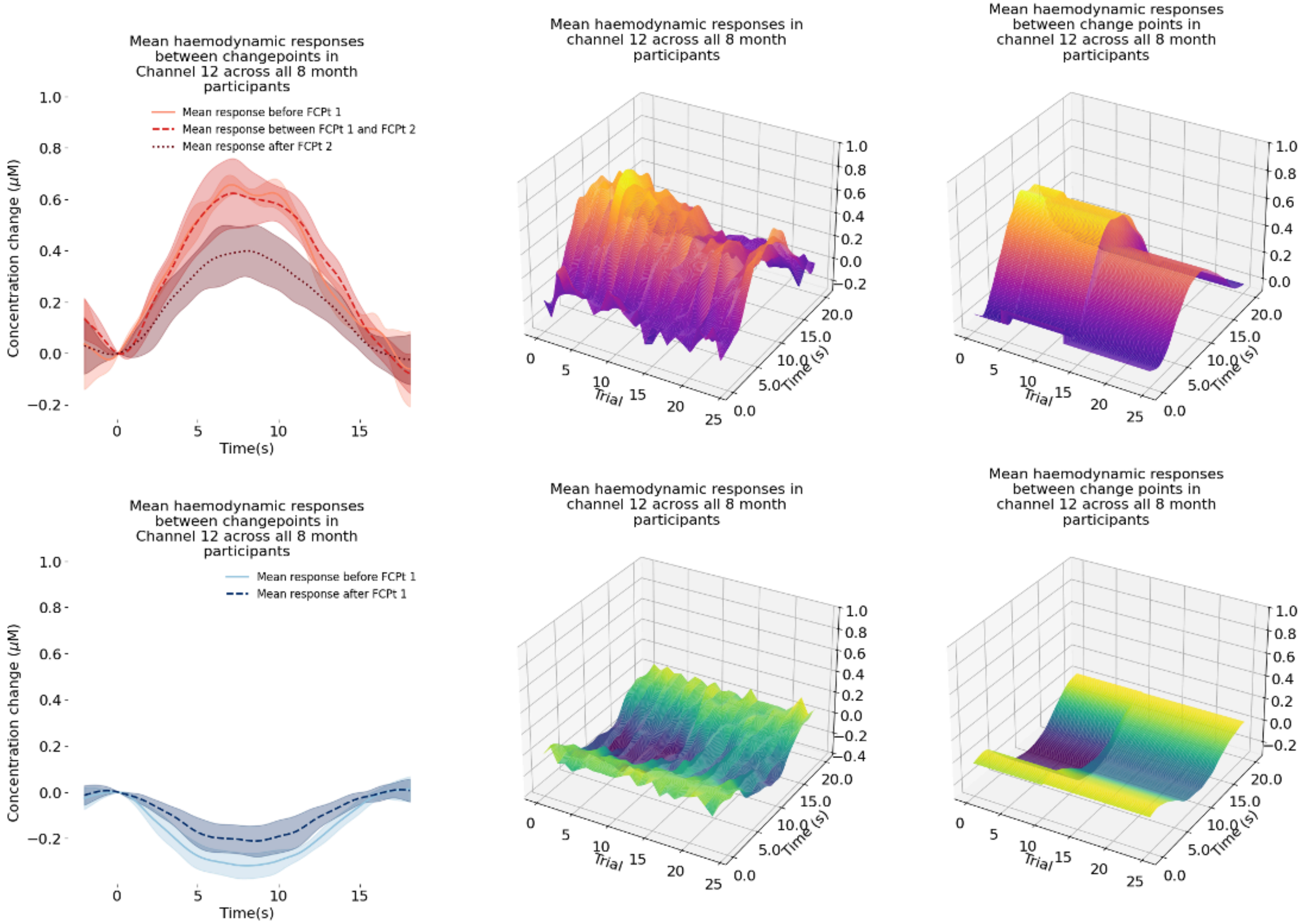
Mean haemodynamic responses between changepoints in Channel 12 across all 8 month participants
Mean response before FCPt 1
Mean response between FCPt 1 and FCPt 2
Mean response after FCPt 2
Concentration change (μM)
Time(s)
Mean haemodynamic responses in channel 12 across all 8 month participants
Trial
Time (s)
Mean haemodynamic responses between change points in channel 12 across all 8 month participants
Trial
Time (s)
Mean haemodynamic responses between changepoints in Channel 12 across all 8 month participants
Mean response before FCPt 1
Mean response after FCPt 1
Concentration change (μM)
Time(s)
Mean haemodynamic responses in channel 12 across all 8 month participants
Trial
Time (s)
Mean haemodynamic responses between change points in channel 12 across all 8 month participants
Trial
Time (s)

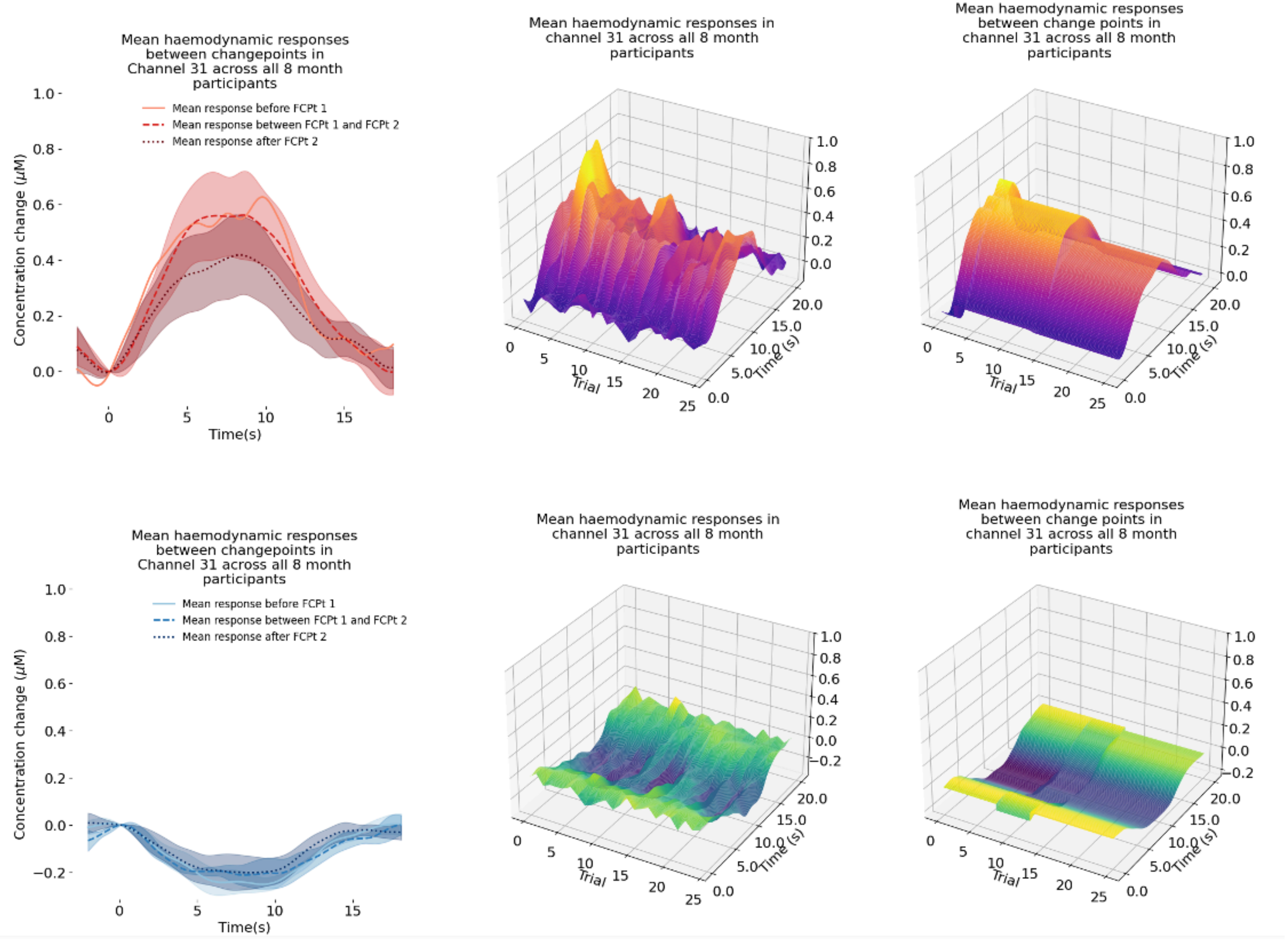
Mean haemodynamic responses between changepoints in Channel 31 across all 8 month participants
Mean response before FCPt 1
Mean response between FCPt 1 and FCPt 2
Mean response after FCPt 2
Concentration change (μM)
Time(s)
Mean haemodynamic responses in channel 31 across all 8 month participants
Trial
Time (s)
Mean haemodynamic responses between change points in channel 31 across all 8 month participants
Trial
Time (s)
Mean haemodynamic responses between changepoints in Channel 31 across all 8 month participants
Mean response before FCPt 1
Mean response between FCPt 1 and FCPt 2
Mean response after FCPt 2
Concentration change (μM)
Time(s)
Mean haemodynamic responses in channel 31 across all 8 month participants
Trial
Time (s)
Mean haemodynamic responses between change points in channel 31 across all 8 month participants
Trial
Time (s)

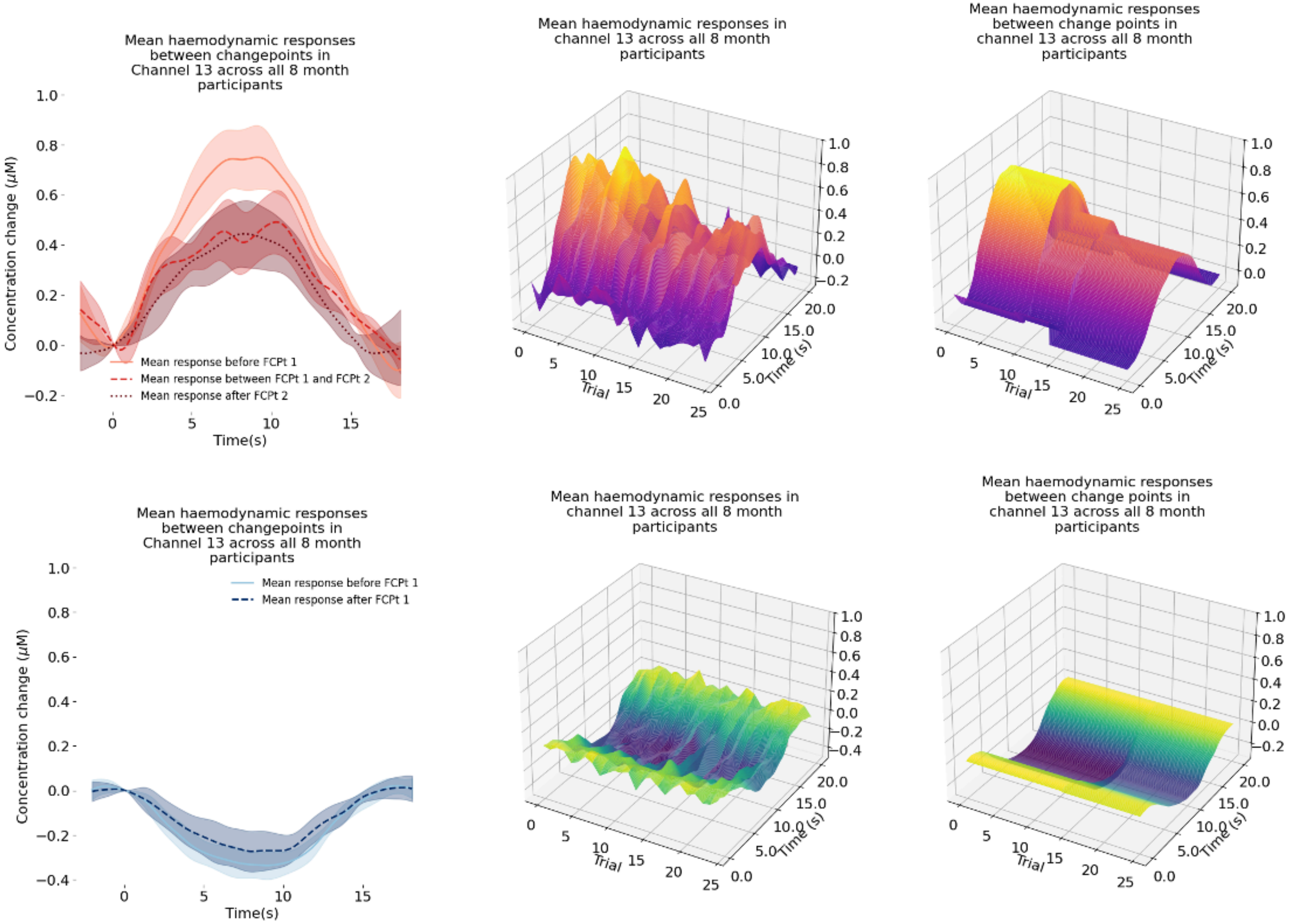
Mean haemodynamic responses between changepoints in Channel 13 across all 8 month participants
Concentration change (μM)
Time(s)
Mean response before FCPt 1
Mean response between FCPt 1 and FCPt 2
Mean response after FCPt 2
Mean haemodynamic responses in channel 13 across all 8 month participants
Trial
Time (s)
Mean haemodynamic responses between change points in channel 13 across all 8 month participants
Trial
Time (s)
Mean haemodynamic responses between changepoints in Channel 13 across all 8 month participants
Concentration change (μM)
Time(s)
Mean response before FCPt 1
Mean response after FCPt 1
Mean haemodynamic responses in channel 13 across all 8 month participants
Trial
Time (s)
Mean haemodynamic responses between change points in channel 13 across all 8 month participants
Trial
Time (s)

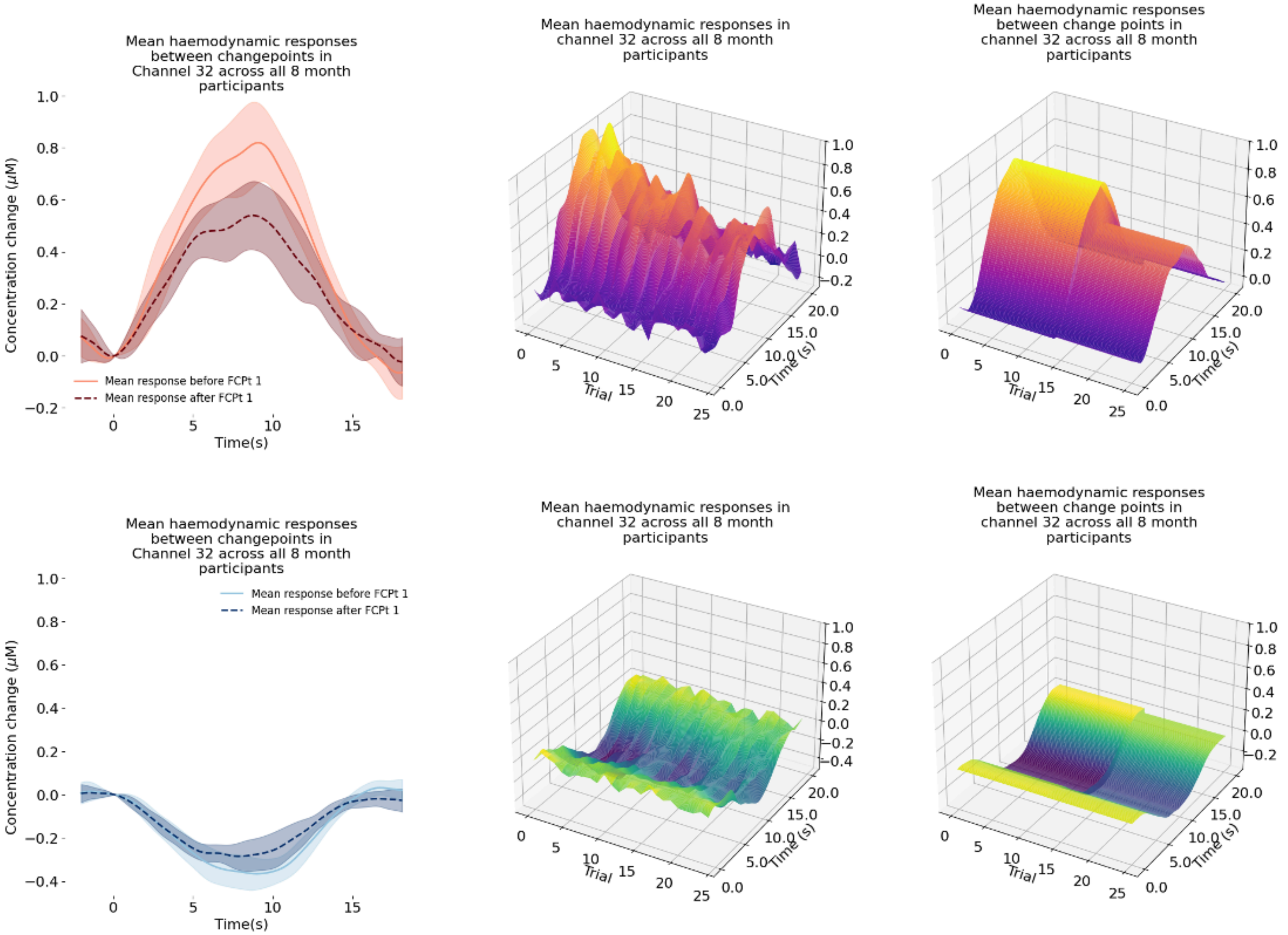
Mean haemodynamic responses between changepoints in Channel 32 across all 8 month participants
Concentration change (μM)
Time(s)
Mean response before FCPt 1
Mean response after FCPt 1
Mean haemodynamic responses in channel 32 across all 8 month participants
Trial
Time (s)
Mean haemodynamic responses between change points in channel 32 across all 8 month participants
Trial
Time (s)
Mean haemodynamic responses between changepoints in Channel 32 across all 8 month participants
Concentration change (μM)
Time(s)
Mean response before FCPt 1
Mean response after FCPt 1
Mean haemodynamic responses in channel 32 across all 8 month participants
Trial
Time (s)
Mean haemodynamic responses between change points in channel 32 across all 8 month participants
Trial
Time (s)

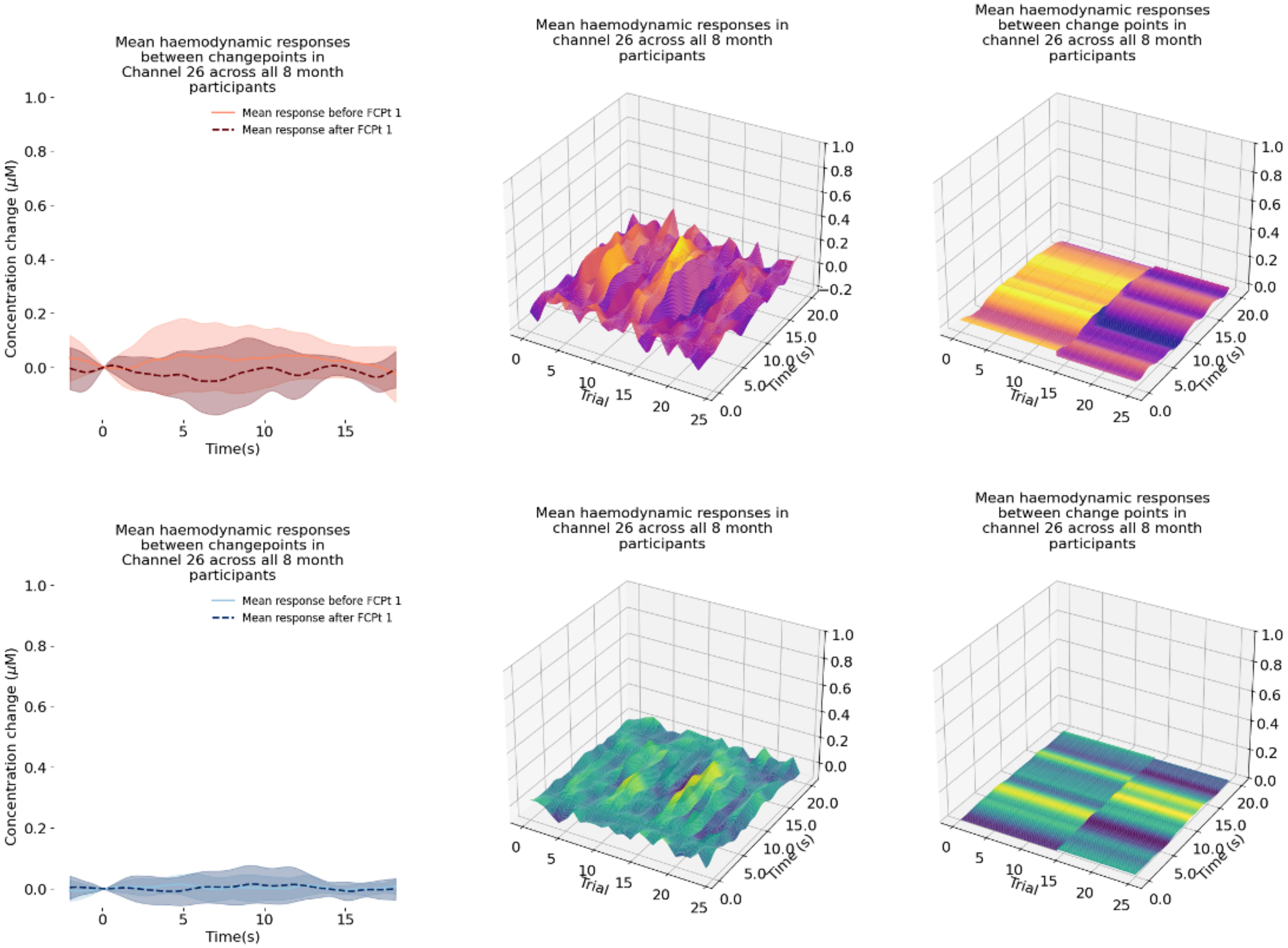
Mean haemodynamic responses between changepoints in Channel 26 across all 8 month participants
Mean response before FCPt 1
Mean response after FCPt 1
Concentration change (μM)
Time(s)
Mean haemodynamic responses in channel 26 across all 8 month participants
Mean haemodynamic responses between change points in channel 26 across all 8 month participants
Trial
Time (s)
Mean haemodynamic responses between changepoints in Channel 26 across all 8 month participants
Mean response before FCPt 1
Mean response after FCPt 1
Concentration change (μM)
Time(s)
Mean haemodynamic responses in channel 26 across all 8 month participants
Mean haemodynamic responses between change points in channel 26 across all 8 month participants
Trial
Time (s)

1847

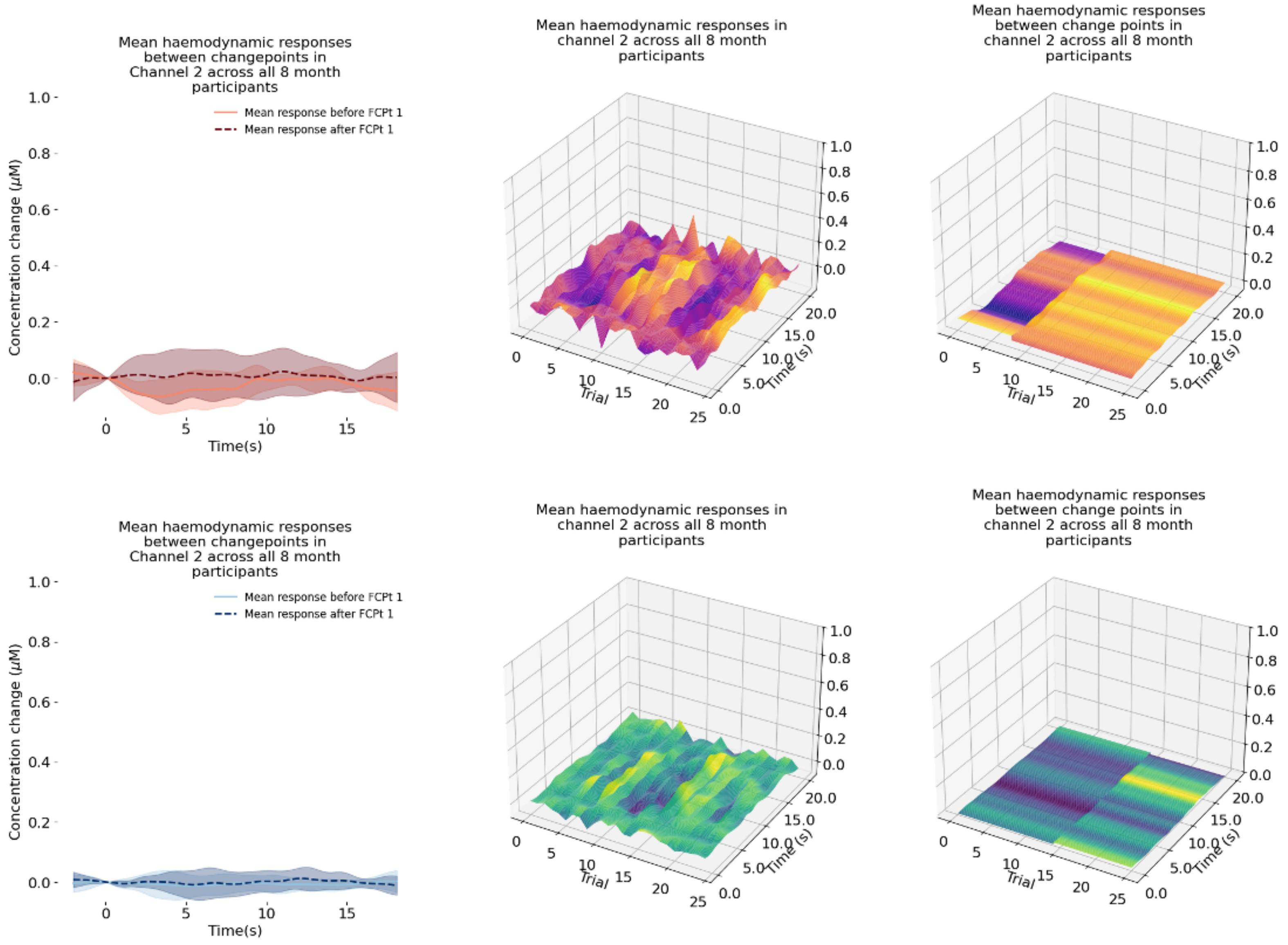
Mean haemodynamic responses between changepoints in Channel 2 across all 8 month participants
Mean response before FCPt 1
Mean response after FCPt 1
Concentration change (μM)
Time(s)
Mean haemodynamic responses in channel 2 across all 8 month participants
Trial
Time (s)
Mean haemodynamic responses between change points in channel 2 across all 8 month participants
Trial
Time (s)
Mean haemodynamic responses between changepoints in Channel 2 across all 8 month participants
Mean response before FCPt 1
Mean response after FCPt 1
Concentration change (μM)
Time(s)
Mean haemodynamic responses in channel 2 across all 8 month participants
Trial
Time (s)
Mean haemodynamic responses between change points in channel 2 across all 8 month participants
Trial
Time (s)

1848

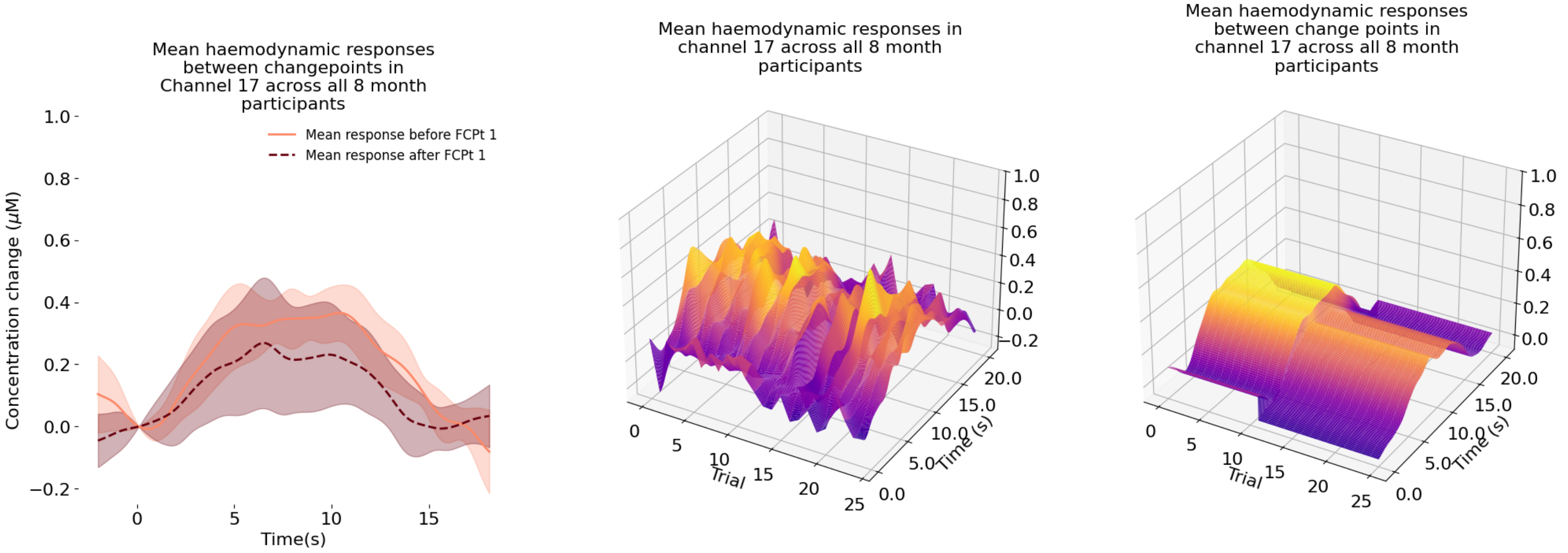
Mean haemodynamic responses between changepoints in Channel 17 across all 8 month participants
Mean response before FCPt 1
Mean response after FCPt 1
Concentration change (μM)
Time(s)
Mean haemodynamic responses in channel 17 across all 8 month participants
Trial
Time (s)
Mean haemodynamic responses between change points in channel 17 across all 8 month participants
Trial
Time (s)

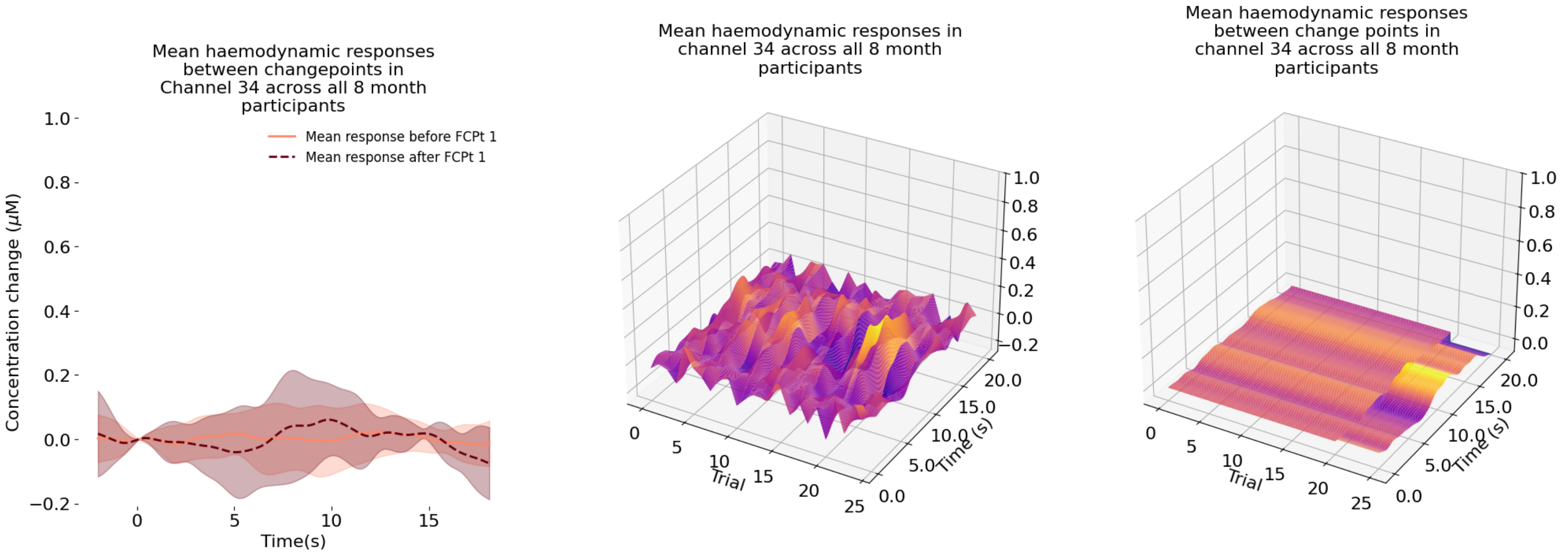
Mean haemodynamic responses between changepoints in Channel 34 across all 8 month participants
Mean response before FCPt 1
Mean response after FCPt 1
Concentration change (μM)
Time(s)
Mean haemodynamic responses in channel 34 across all 8 month participants
Trial
Time (s)
Mean haemodynamic responses between change points in channel 34 across all 8 month participants
Trial
Time (s)

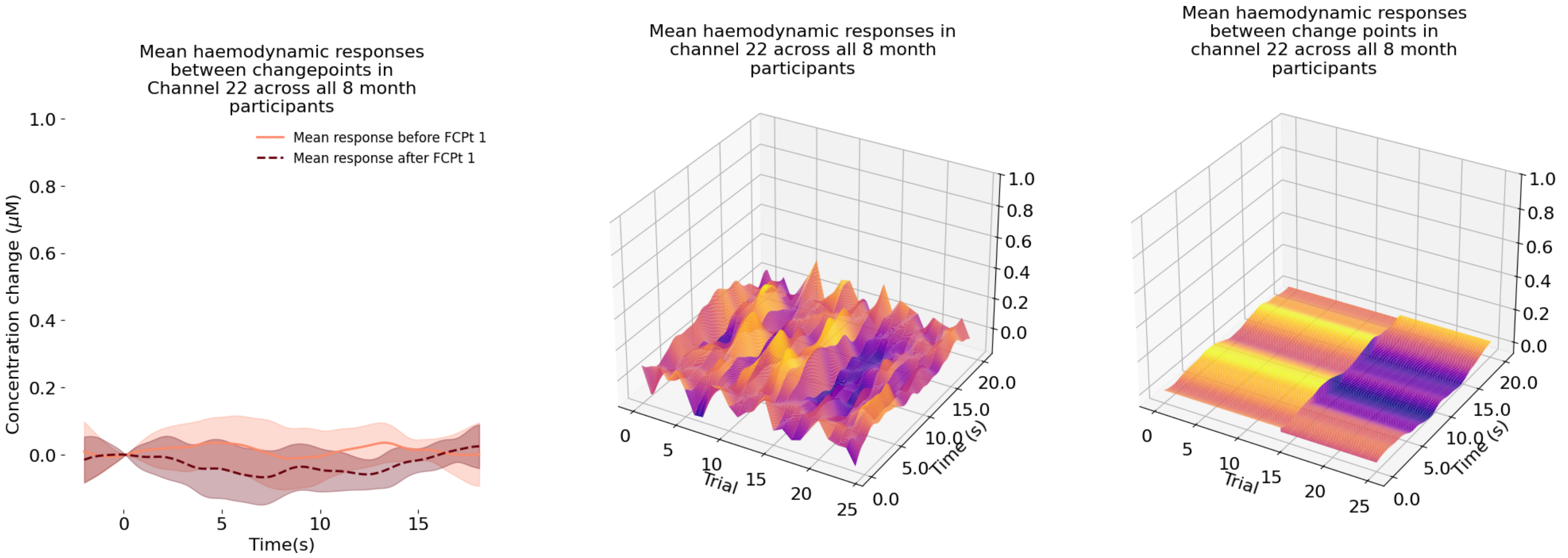
Mean haemodynamic responses between changepoints in Channel 22 across all 8 month participants
Mean response before FCPt 1
Mean response after FCPt 1
Concentration change (μM)
Time(s)
Mean haemodynamic responses in channel 22 across all 8 month participants
Trial
Time (s)
Mean haemodynamic responses between change points in channel 22 across all 8 month participants
Trial
Time (s)

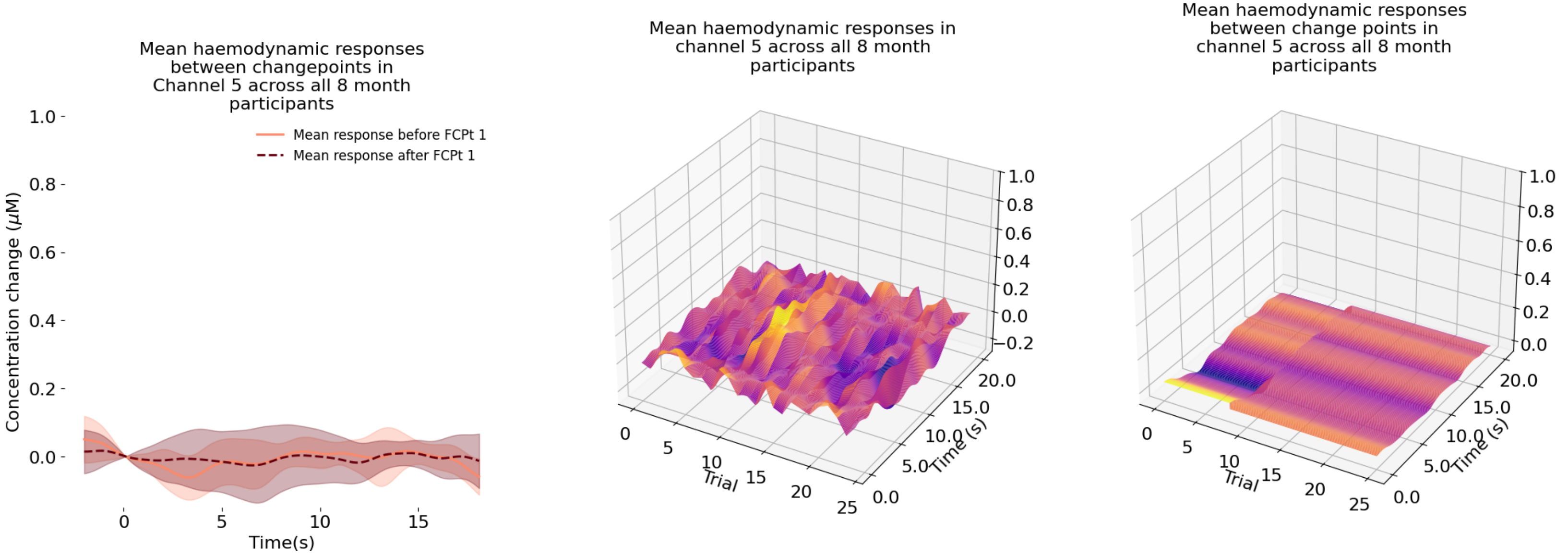
Mean haemodynamic responses between changepoints in Channel 5 across all 8 month participants
Mean response before FCPt 1
Mean response after FCPt 1
Concentration change (μM)
Time(s)
Mean haemodynamic responses in channel 5 across all 8 month participants
Trial
Time (s)
Mean haemodynamic responses between change points in channel 5 across all 8 month participants
Trial
Time (s)

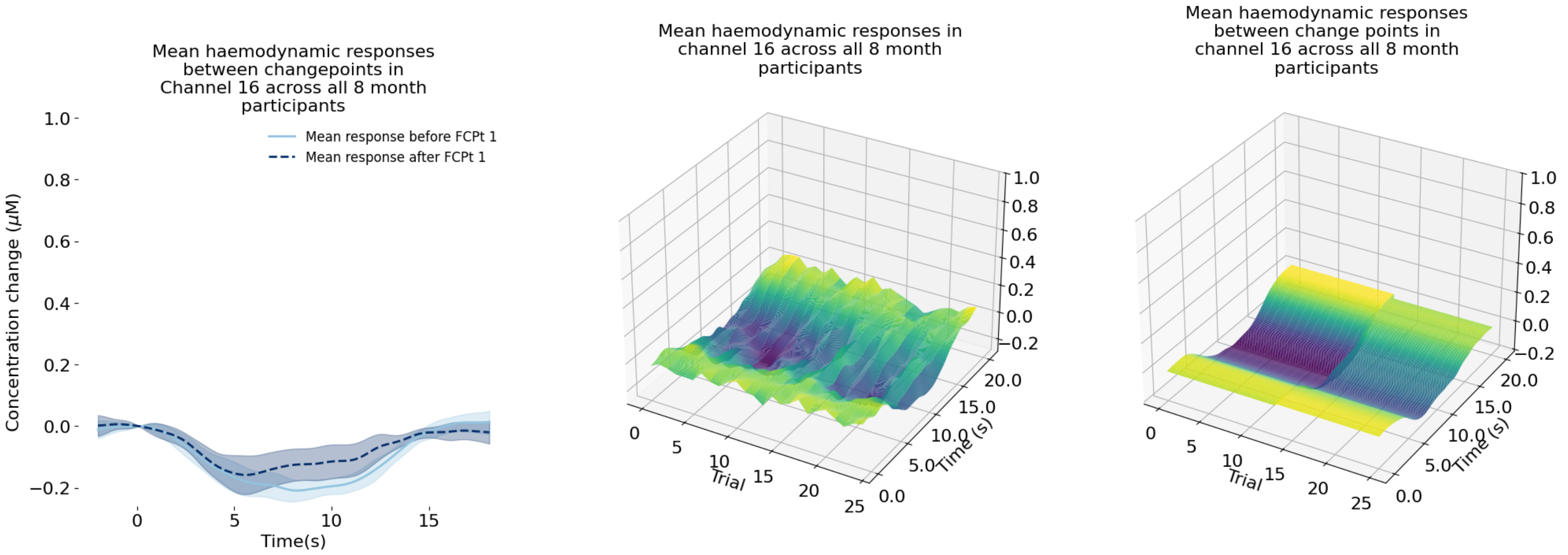
Mean haemodynamic responses between changepoints in Channel 16 across all 8 month participants
Mean response before FCPt 1
Mean response after FCPt 1
Concentration change (μM)
Time(s)
Mean haemodynamic responses in channel 16 across all 8 month participants
Trial
Time (s)
Mean haemodynamic responses between change points in channel 16 across all 8 month participants
Trial
Time (s)

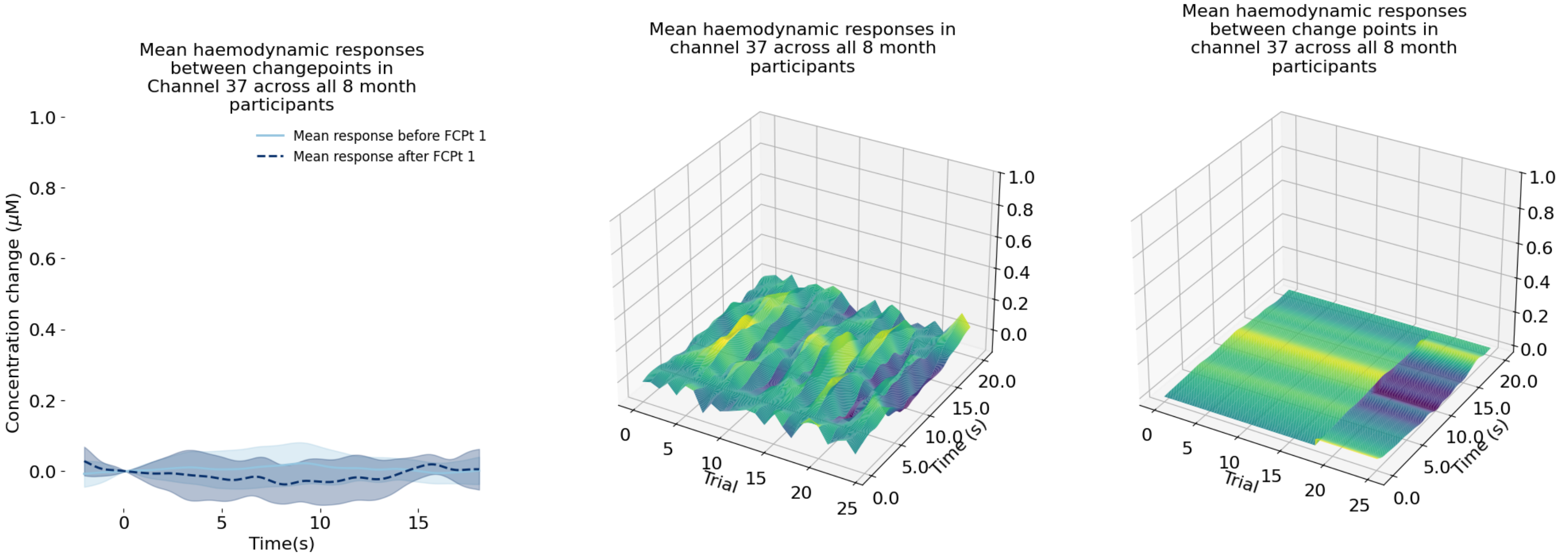
Mean haemodynamic responses between changepoints in Channel 37 across all 8 month participants
Mean response before FCPt 1
Mean response after FCPt 1
Concentration change (μM)
Time(s)
Mean haemodynamic responses in channel 37 across all 8 month participants
Trial
Time (s)
Mean haemodynamic responses between change points in channel 37 across all 8 month participants
Trial
Time (s)

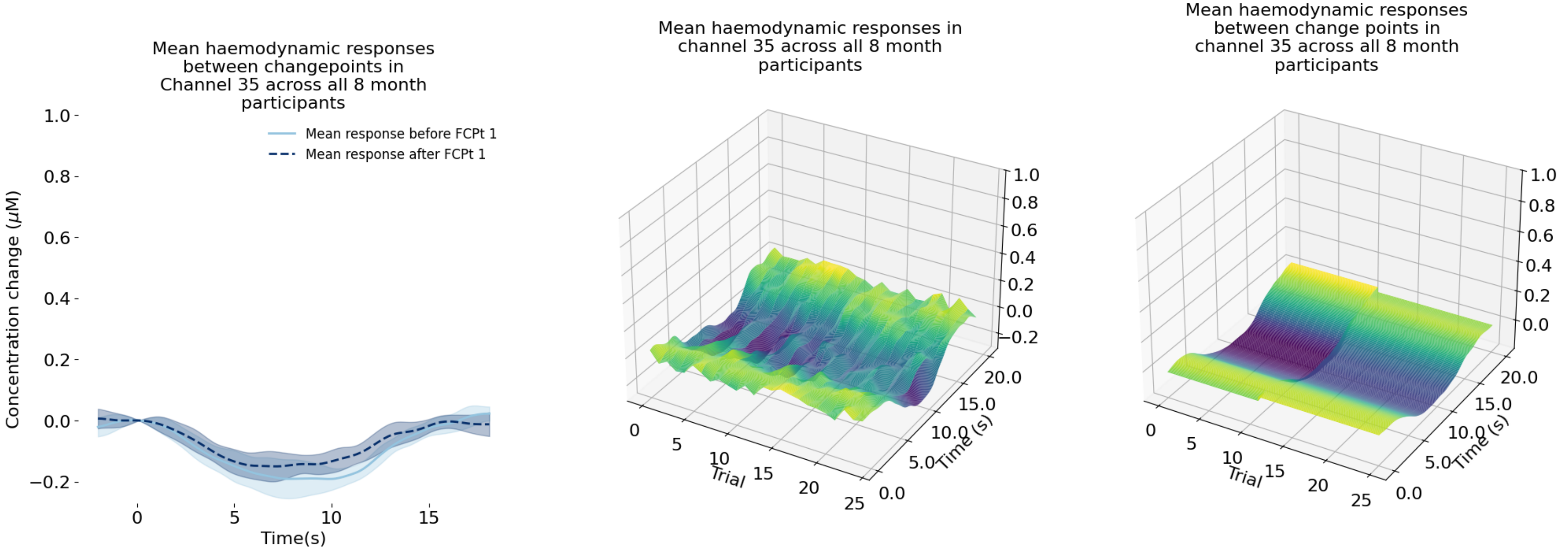
Mean haemodynamic responses between changepoints in Channel 35 across all 8 month participants
Mean response before FCPt 1
Mean response after FCPt 1
Concentration change (μM)
Time(s)
Mean haemodynamic responses in channel 35 across all 8 month participants
Trial
Time (s)
Mean haemodynamic responses between change points in channel 35 across all 8 month participants
Trial
Time (s)

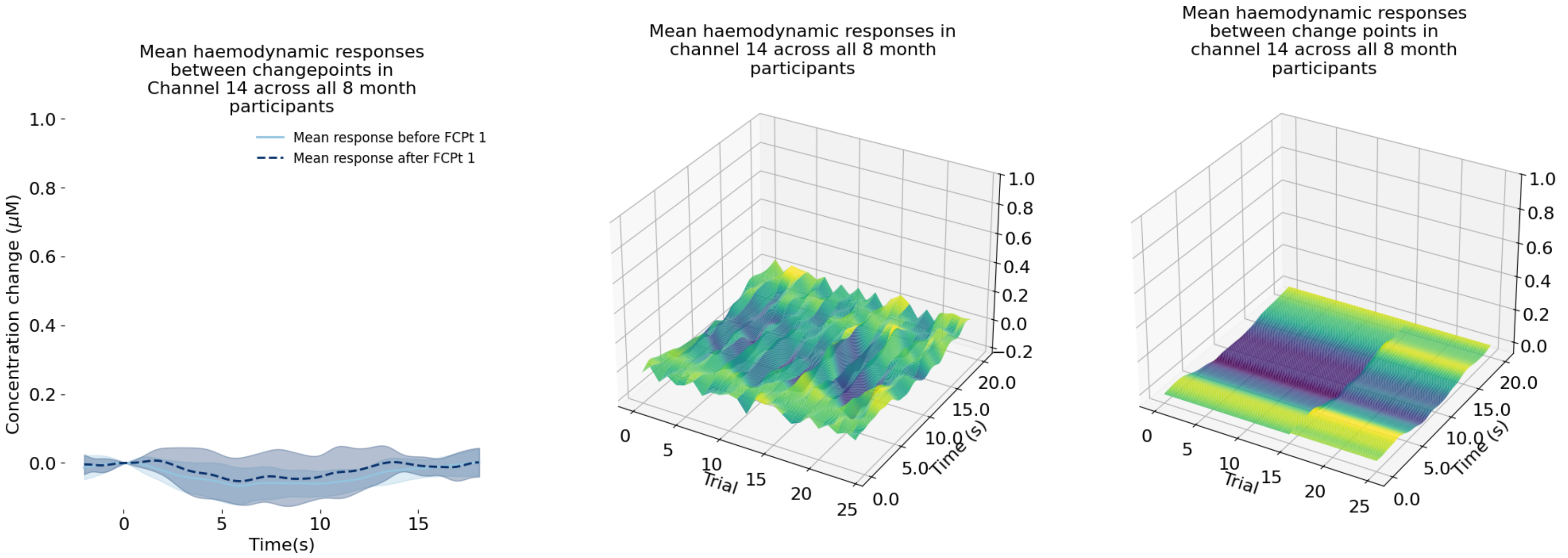
Mean haemodynamic responses between changepoints in Channel 14 across all 8 month participants
Mean response before FCPt 1
Mean response after FCPt 1
Concentration change (μM)
Time(s)
Mean haemodynamic responses in channel 14 across all 8 month participants
Trial
Time (s)
Mean haemodynamic responses between change points in channel 14 across all 8 month participants
Trial
Time (s)

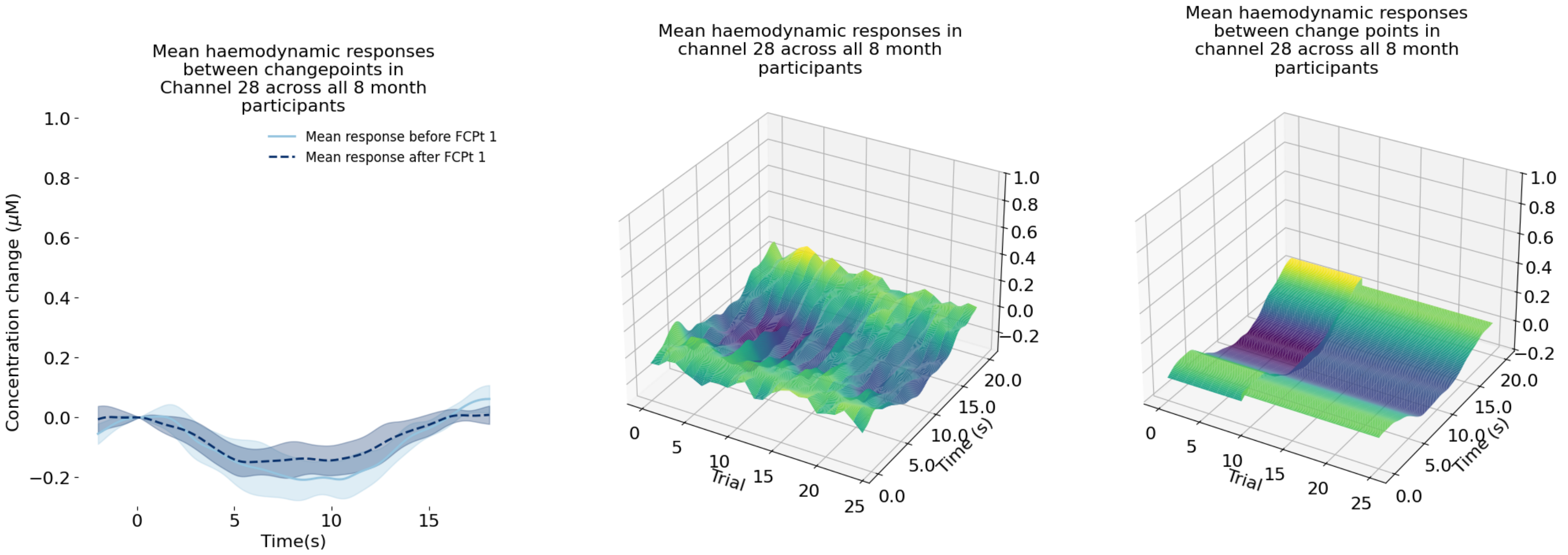
Mean haemodynamic responses between changepoints in Channel 28 across all 8 month participants
Mean response before FCPt 1
Mean response after FCPt 1
Concentration change (μM)
Time(s)
Mean haemodynamic responses in channel 28 across all 8 month participants
Trial
Time (s)
Mean haemodynamic responses between change points in channel 28 across all 8 month participants
Trial
Time (s)

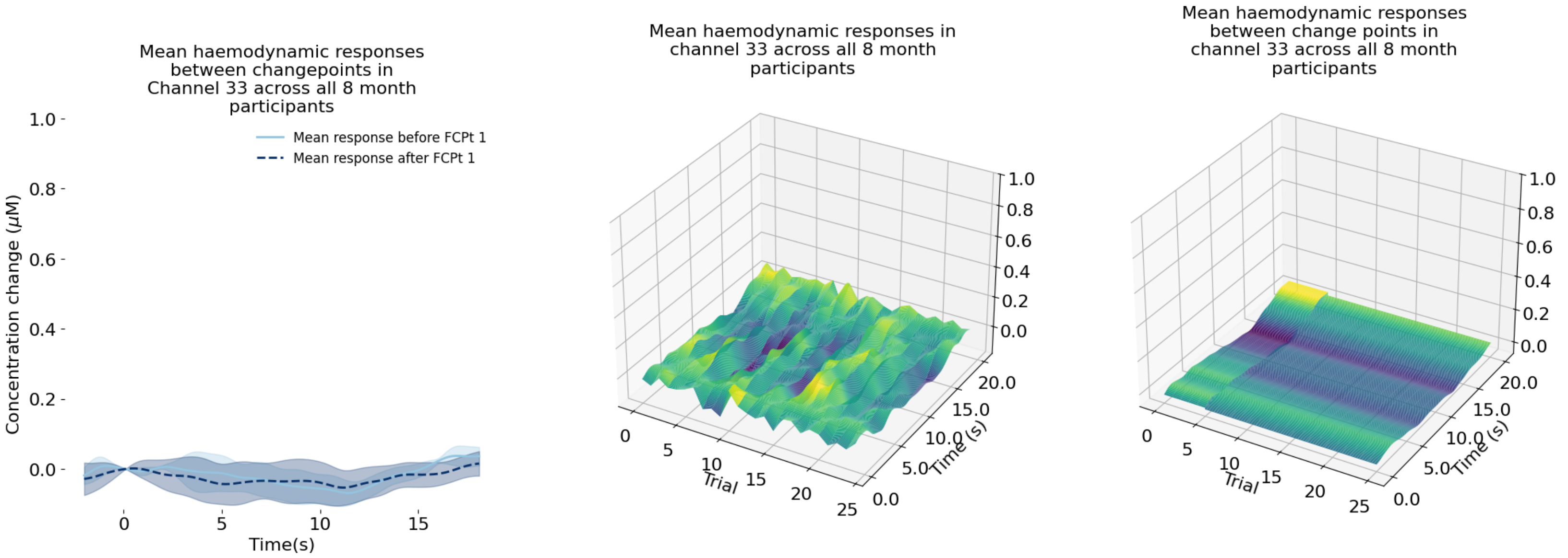
Mean haemodynamic responses between changepoints in Channel 33 across all 8 month participants
Mean response before FCPt 1
Mean response after FCPt 1
Concentration change (μM)
Time(s)
Mean haemodynamic responses in channel 33 across all 8 month participants
Trial
Time (s)
Mean haemodynamic responses between change points in channel 33 across all 8 month participants
Trial
Time (s)

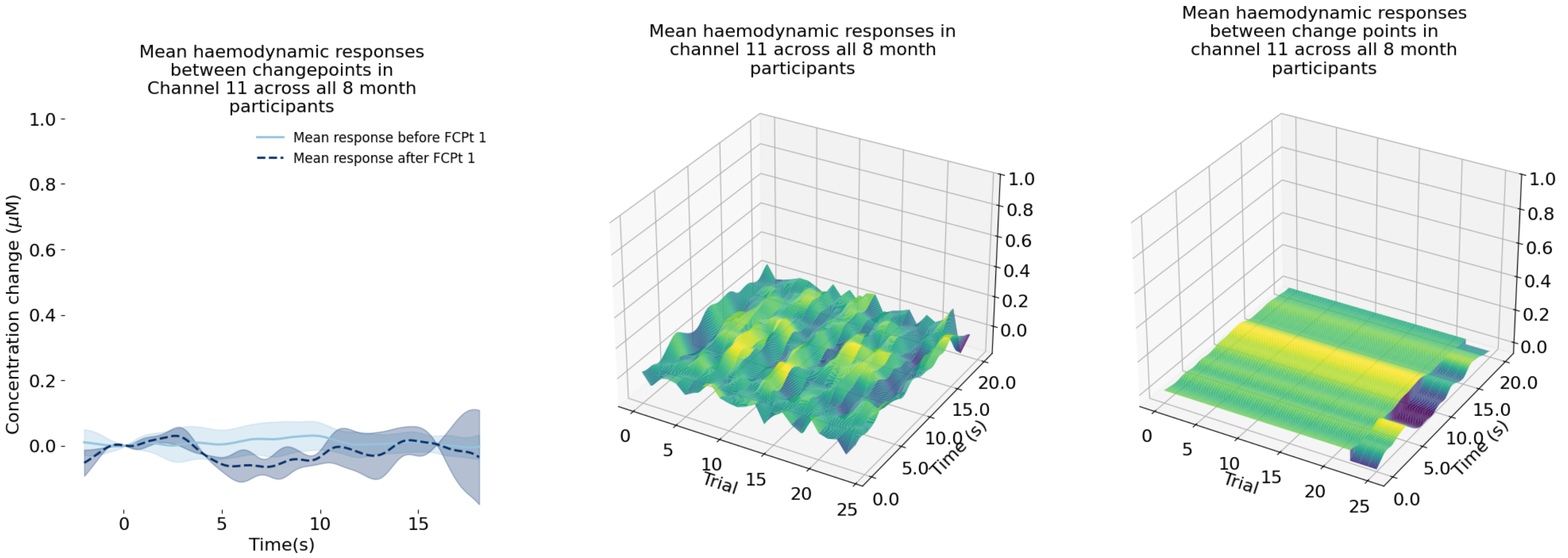
Mean haemodynamic responses between changepoints in Channel 11 across all 8 month participants
Mean response before FCPt 1
Mean response after FCPt 1
Concentration change (μM)
Time(s)
Mean haemodynamic responses in channel 11 across all 8 month participants
Trial
Time (s)
Mean haemodynamic responses between change points in channel 11 across all 8 month participants
Trial
Time (s)

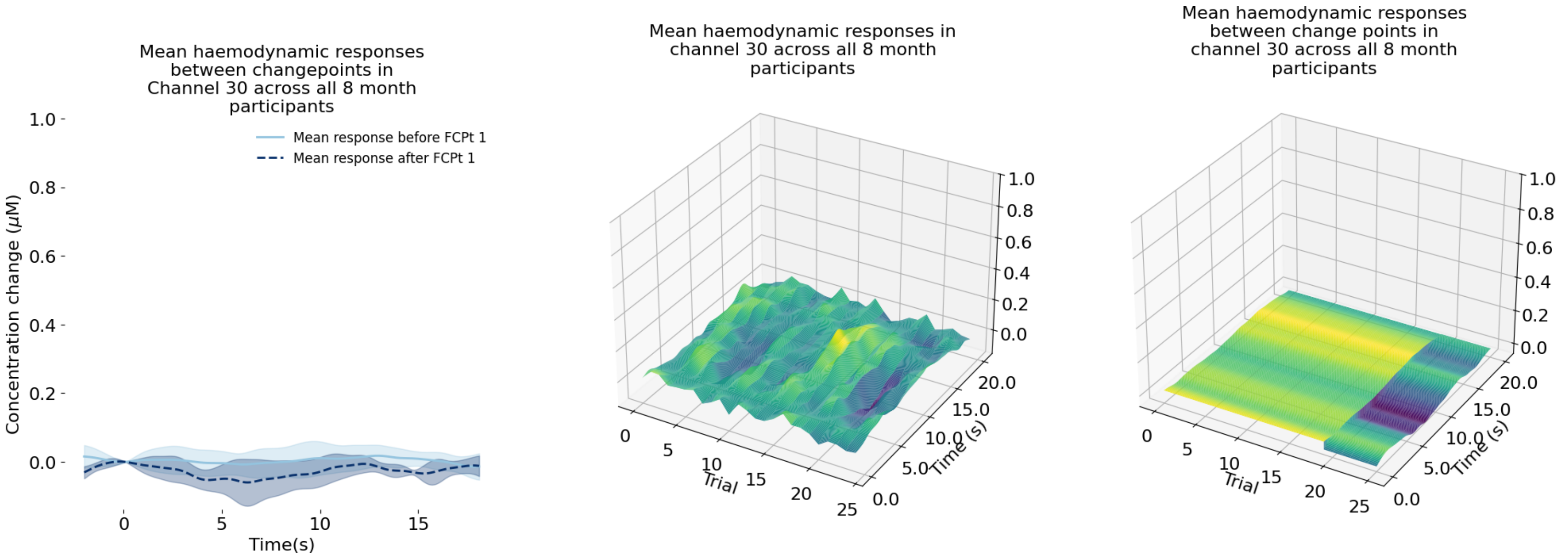
Mean haemodynamic responses between changepoints in Channel 30 across all 8 month participants
Mean response before FCPt 1
Mean response after FCPt 1
Concentration change (μM)
Time(s)
Mean haemodynamic responses in channel 30 across all 8 month participants
Trial
Time (s)
Mean haemodynamic responses between change points in channel 30 across all 8 month participants
Trial
Time (s)

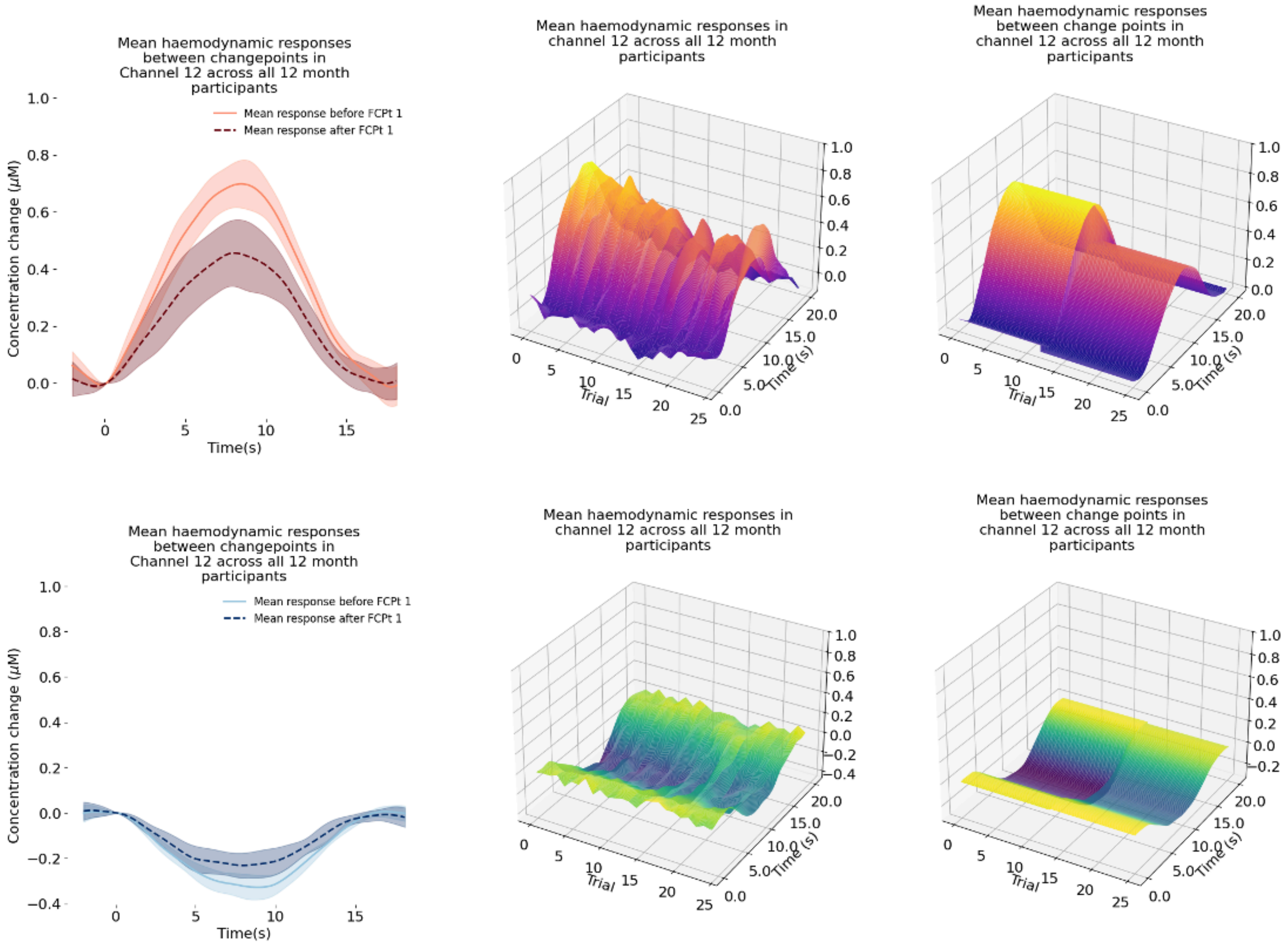

Mean haemodynamic responses between changepoints in Channel 12 across all 12 month participants
Mean response before FCPt 1
Mean response after FCPt 1
Concentration change (μM)
Time(s)
Mean haemodynamic responses in channel 12 across all 12 month participants
Trial
Time (s)
Mean haemodynamic responses between change points in channel 12 across all 12 month participants
Trial
Time (s)
Mean haemodynamic responses between changepoints in Channel 12 across all 12 month participants
Mean response before FCPt 1
Mean response after FCPt 1
Concentration change (μM)
Time(s)
Mean haemodynamic responses in channel 12 across all 12 month participants
Trial
Time (s)
Mean haemodynamic responses between change points in channel 12 across all 12 month participants
Trial
Time (s)

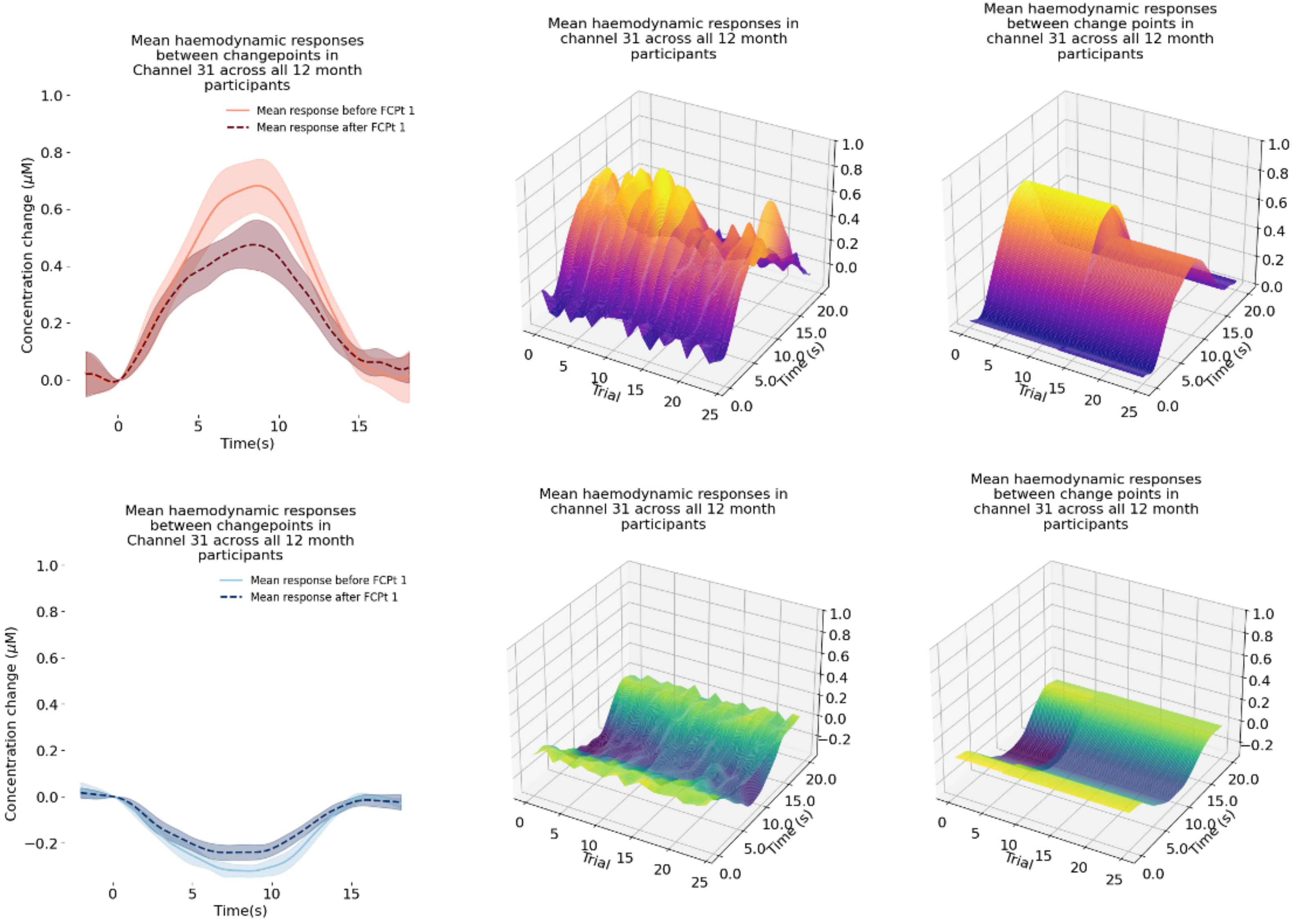
Mean haemodynamic responses between changepoints in Channel 31 across all 12 month participants
Mean response before FCPt 1
Mean response after FCPt 1
Concentration change (μM)
Time(s)
Mean haemodynamic responses in channel 31 across all 12 month participants
Trial
Time (s)
Mean haemodynamic responses between change points in channel 31 across all 12 month participants
Trial
Time (s)
Mean haemodynamic responses between changepoints in Channel 31 across all 12 month participants
Mean response before FCPt 1
Mean response after FCPt 1
Concentration change (μM)
Time(s)
Mean haemodynamic responses in channel 31 across all 12 month participants
Trial
Time (s)
Mean haemodynamic responses between change points in channel 31 across all 12 month participants
Trial
Time (s)

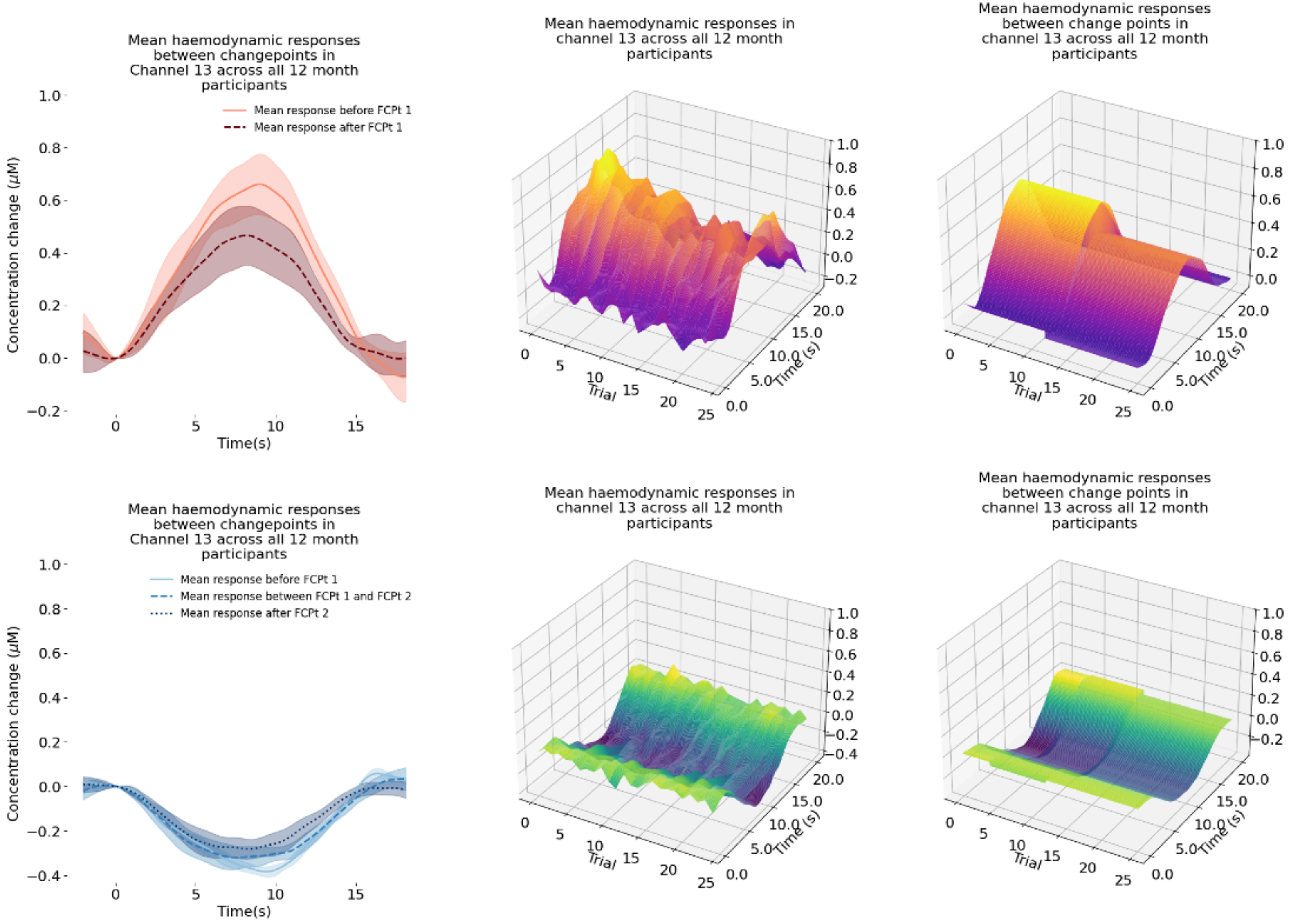
Mean haemodynamic responses between changepoints in Channel 13 across all 12 month participants
Mean response before FCPt 1
Mean response after FCPt 1
Concentration change (μM)
Time(s)
Mean haemodynamic responses in channel 13 across all 12 month participants
Trial
Time (s)
Mean haemodynamic responses between change points in channel 13 across all 12 month participants
Trial
Time (s)
Mean haemodynamic responses between changepoints in Channel 13 across all 12 month participants
Mean response before FCPt 1
Mean response between FCPt 1 and FCPt 2
Mean response after FCPt 2
Concentration change (μM)
Time(s)
Mean haemodynamic responses in channel 13 across all 12 month participants
Trial
Time (s)
Mean haemodynamic responses between change points in channel 13 across all 12 month participants
Trial
Time (s)

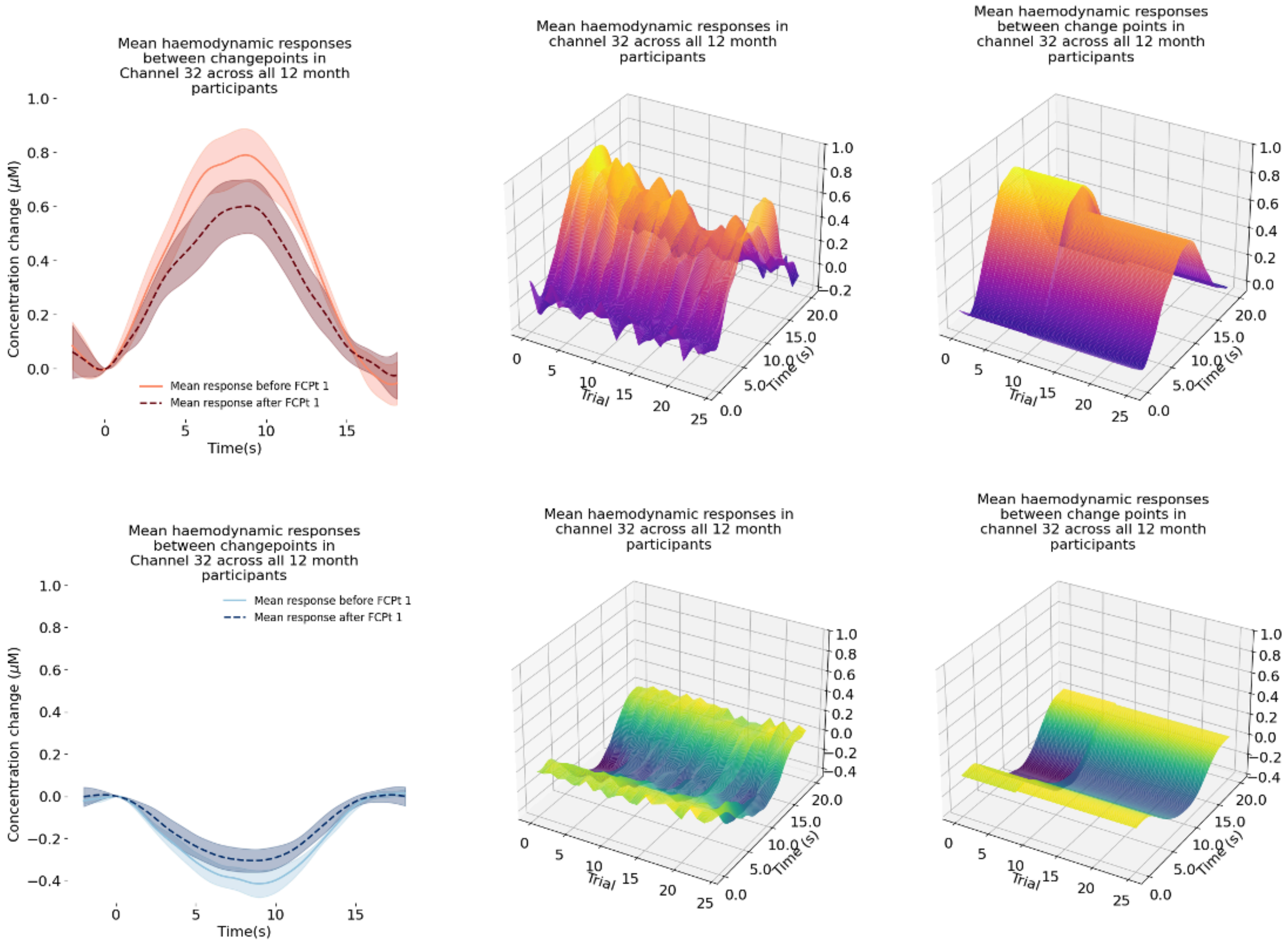
Mean haemodynamic responses between changepoints in Channel 32 across all 12 month participants
Concentration change (μM)
Time(s)
Mean response before FCPt 1
Mean response after FCPt 1
Mean haemodynamic responses in channel 32 across all 12 month participants
Trial
Time (s)
Mean haemodynamic responses between change points in channel 32 across all 12 month participants
Trial
Time (s)
Mean haemodynamic responses between changepoints in Channel 32 across all 12 month participants
Concentration change (μM)
Time(s)
Mean response before FCPt 1
Mean response after FCPt 1
Mean haemodynamic responses in channel 32 across all 12 month participants
Trial
Time (s)
Mean haemodynamic responses between change points in channel 32 across all 12 month participants
Trial
Time (s)

1996

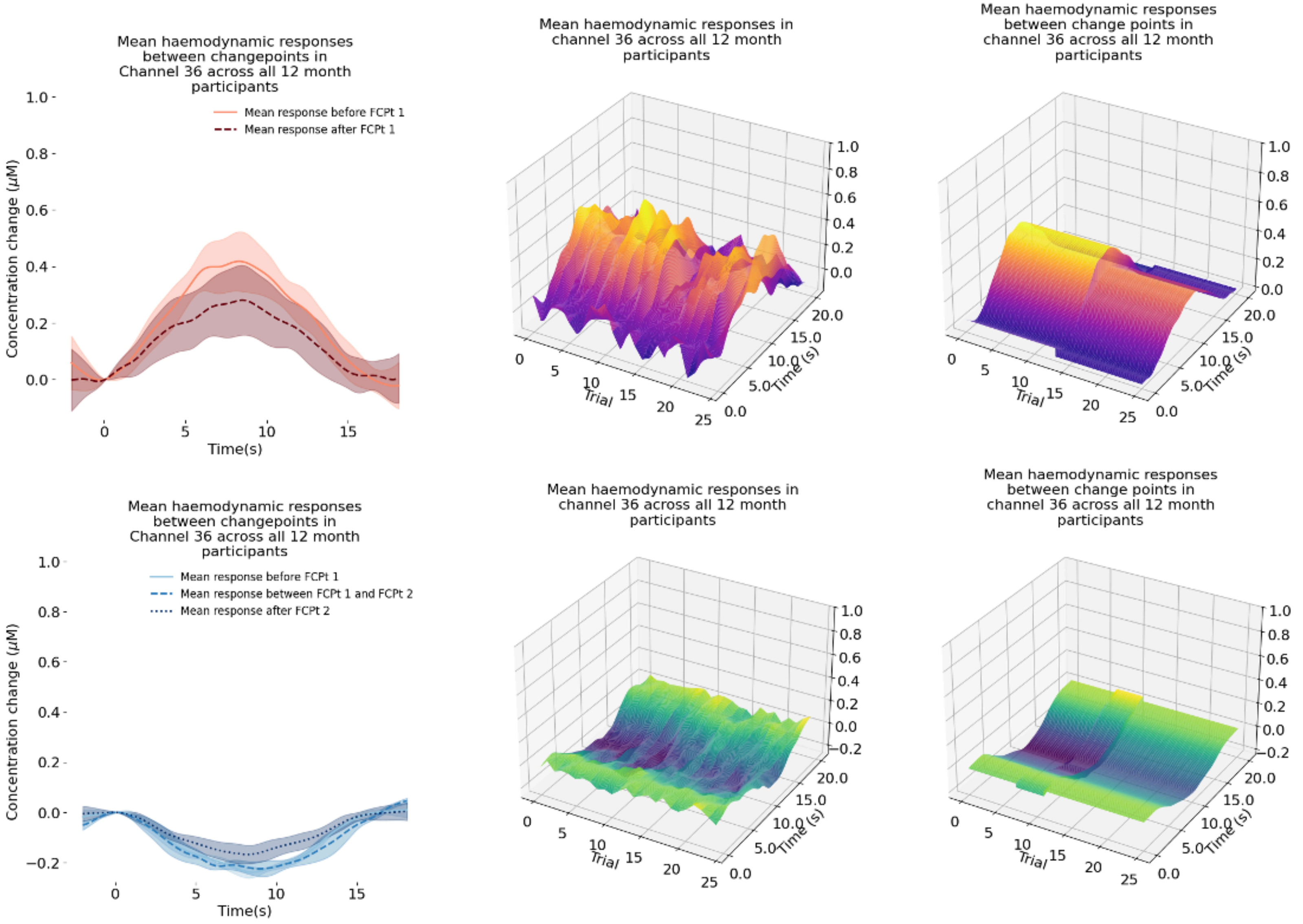
Mean haemodynamic responses between changepoints in Channel 36 across all 12 month participants
Mean response before FCPt 1
Mean response after FCPt 1
Concentration change (μM)
Time(s)
Mean haemodynamic responses in channel 36 across all 12 month participants
Mean haemodynamic responses between change points in channel 36 across all 12 month participants
Trial
Time (s)
Mean haemodynamic responses between changepoints in Channel 36 across all 12 month participants
Mean response before FCPt 1
Mean response between FCPt 1 and FCPt 2
Mean response after FCPt 2
Concentration change (μM)
Time(s)
Mean haemodynamic responses in channel 36 across all 12 month participants
Mean haemodynamic responses between change points in channel 36 across all 12 month participants
Trial
Time (s)

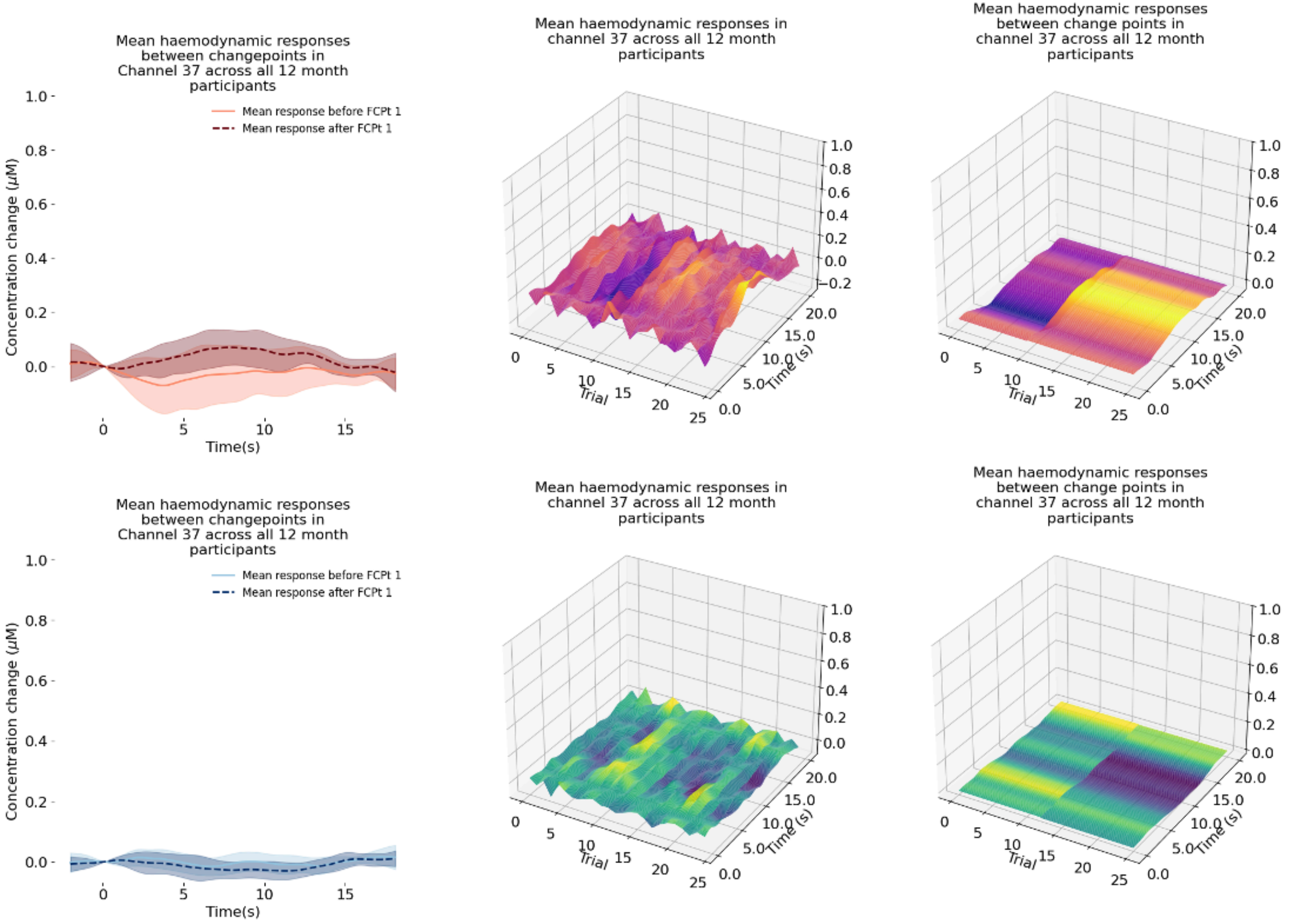
Mean haemodynamic responses between changepoints in Channel 37 across all 12 month participants
Mean response before FCPt 1
Mean response after FCPt 1
Concentration change (μM)
Time(s)
Mean haemodynamic responses in channel 37 across all 12 month participants
Mean haemodynamic responses between change points in channel 37 across all 12 month participants
Trial
Time (s)
Mean haemodynamic responses between changepoints in Channel 37 across all 12 month participants
Mean response before FCPt 1
Mean response after FCPt 1
Concentration change (μM)
Time(s)
Mean haemodynamic responses in channel 37 across all 12 month participants
Mean haemodynamic responses between change points in channel 37 across all 12 month participants
Trial
Time (s)

1998

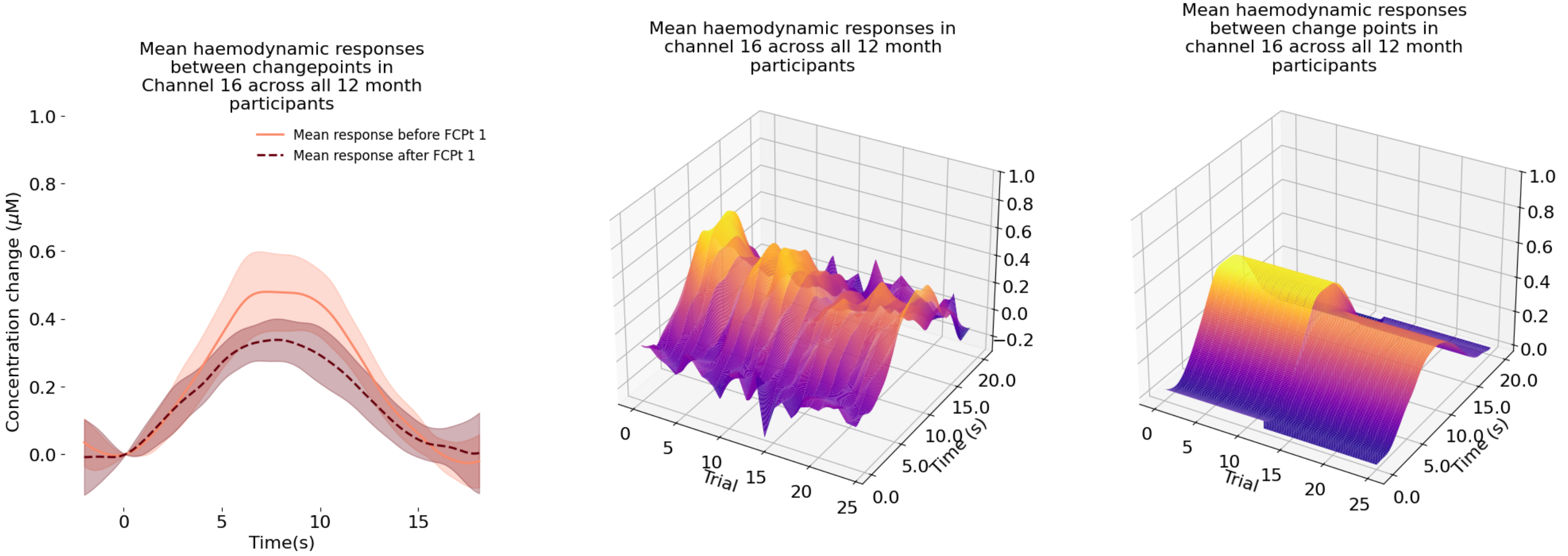
Mean haemodynamic responses between changepoints in Channel 16 across all 12 month participants
Mean response before FCPt 1
Mean response after FCPt 1
Concentration change (μM)
Time(s)
Mean haemodynamic responses in channel 16 across all 12 month participants
Trial
Time (s)
Mean haemodynamic responses between change points in channel 16 across all 12 month participants
Trial
Time (s)

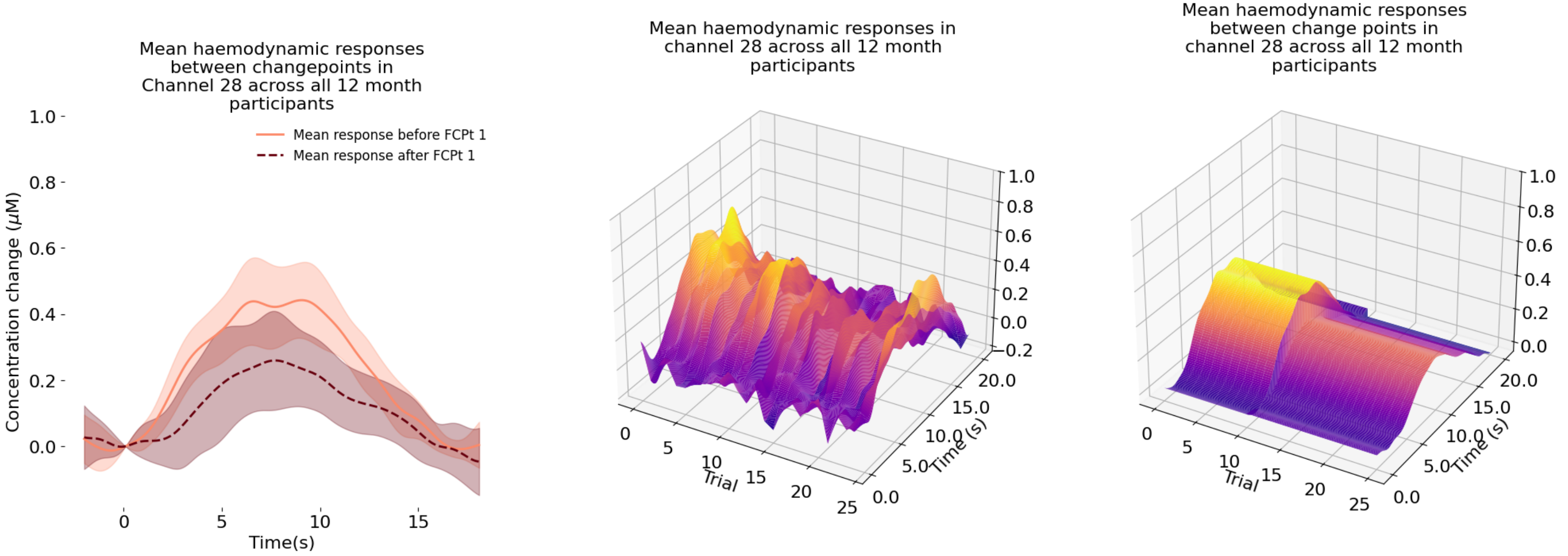
Mean haemodynamic responses between changepoints in Channel 28 across all 12 month participants
Mean response before FCPt 1
Mean response after FCPt 1
Concentration change ($\mu$M)
Time(s)
Mean haemodynamic responses in channel 28 across all 12 month participants
Trial
Time (s)
Mean haemodynamic responses between change points in channel 28 across all 12 month participants
Trial
Time (s)

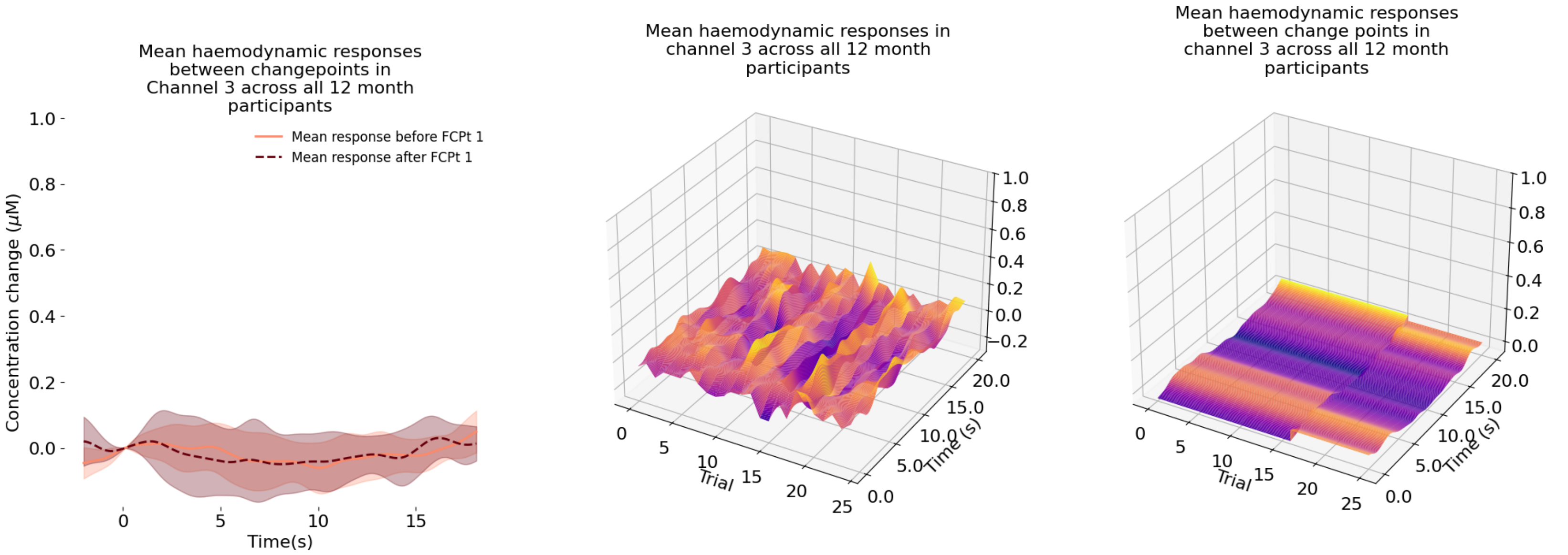
Mean haemodynamic responses between changepoints in Channel 3 across all 12 month participants
Mean response before FCPt 1
Mean response after FCPt 1
Concentration change (μM)
Time(s)
Mean haemodynamic responses in channel 3 across all 12 month participants
Trial
Time (s)
Mean haemodynamic responses between change points in channel 3 across all 12 month participants
Trial
Time (s)

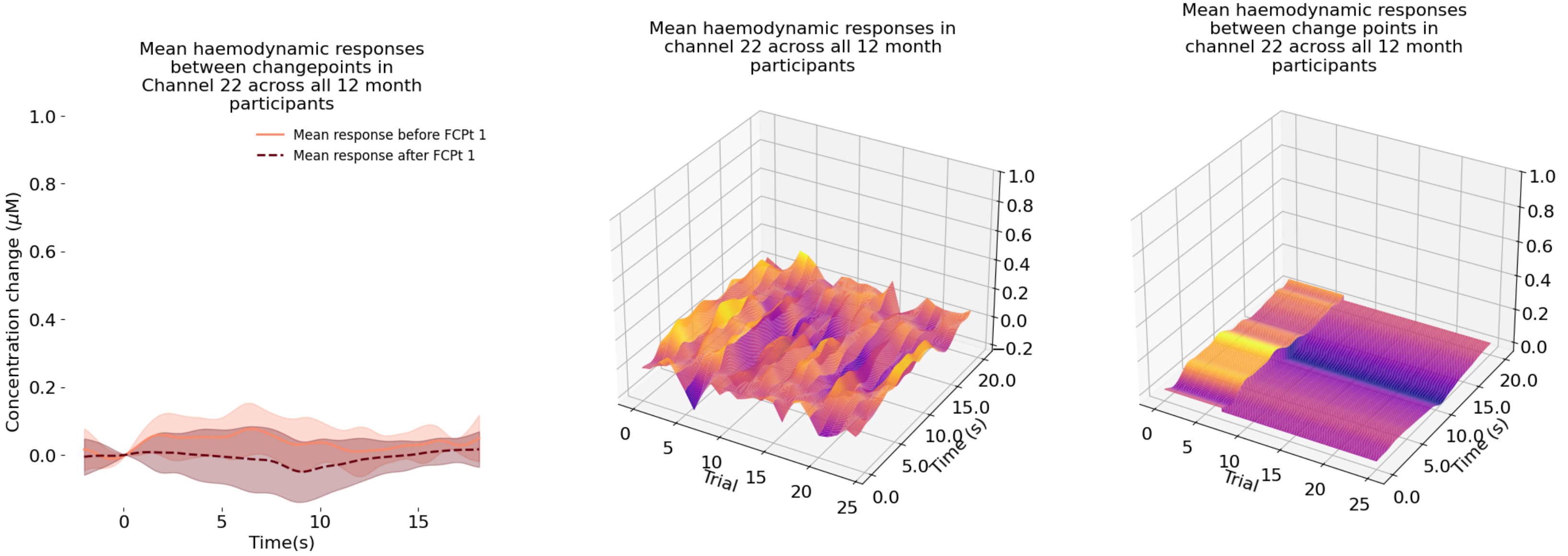
Mean haemodynamic responses between changepoints in Channel 22 across all 12 month participants
Mean response before FCPt 1
Mean response after FCPt 1
Concentration change (μM)
Time(s)
Mean haemodynamic responses in channel 22 across all 12 month participants
Trial
Time (s)
Mean haemodynamic responses between change points in channel 22 across all 12 month participants
Trial
Time (s)

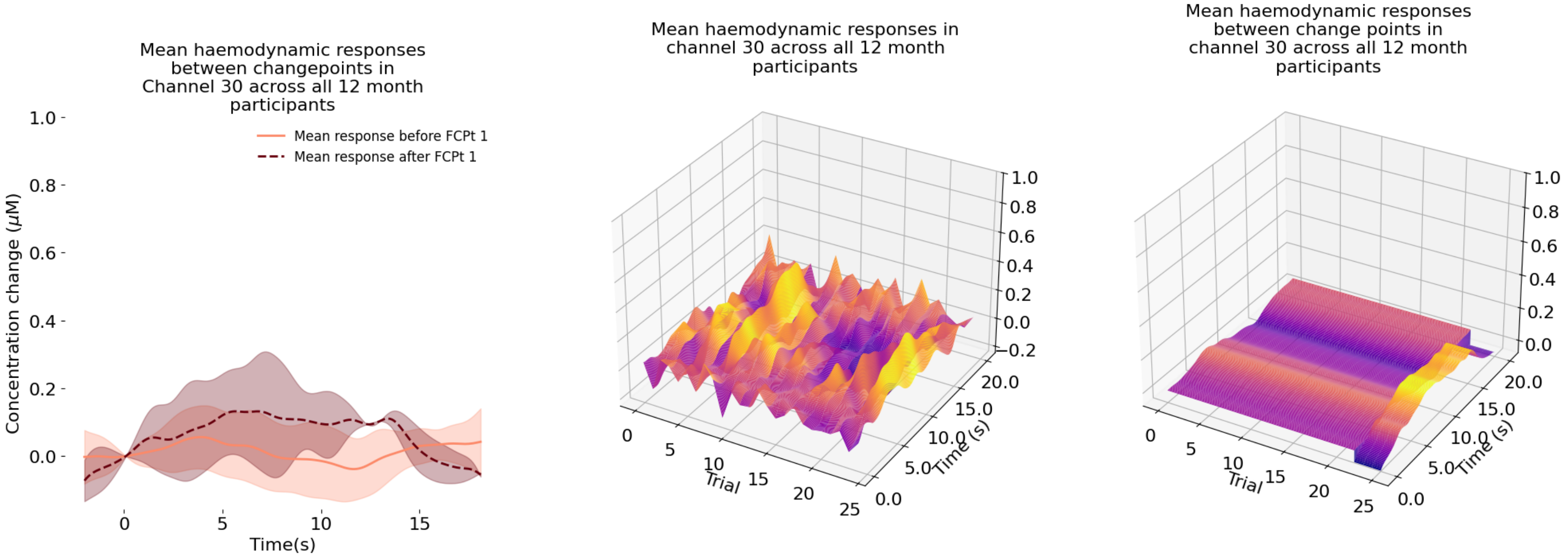
Mean haemodynamic responses between changepoints in Channel 30 across all 12 month participants
Mean response before FCPt 1
Mean response after FCPt 1
Concentration change (μM)
Time(s)
Mean haemodynamic responses in channel 30 across all 12 month participants
Trial
Time (s)
Mean haemodynamic responses between change points in channel 30 across all 12 month participants
Trial
Time (s)

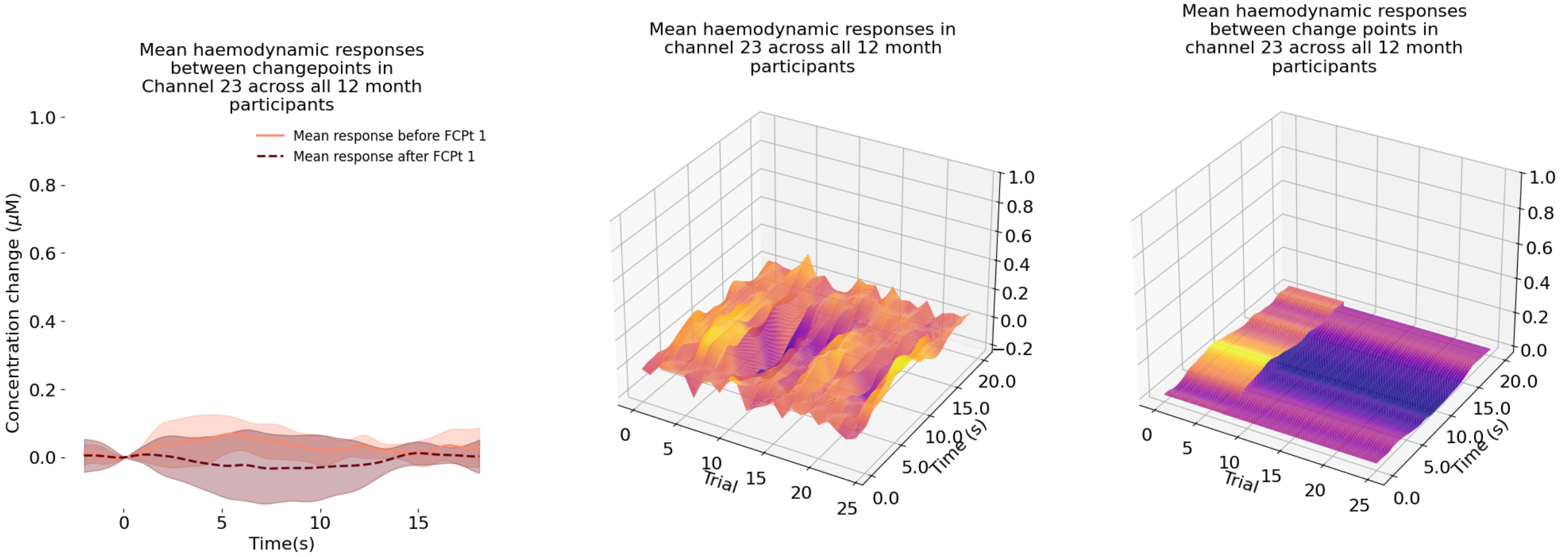
Mean haemodynamic responses between changepoints in Channel 23 across all 12 month participants
Mean response before FCPt 1
Mean response after FCPt 1
Concentration change (μM)
1.0
0.8
0.6
0.4
0.2
0.0
0
5
10
15
Time(s)
Mean haemodynamic responses in channel 23 across all 12 month participants
Trial
Time (s)
-0.2
0.0
5.0
10.0
15.0
20.0
25
Mean haemodynamic responses between change points in channel 23 across all 12 month participants

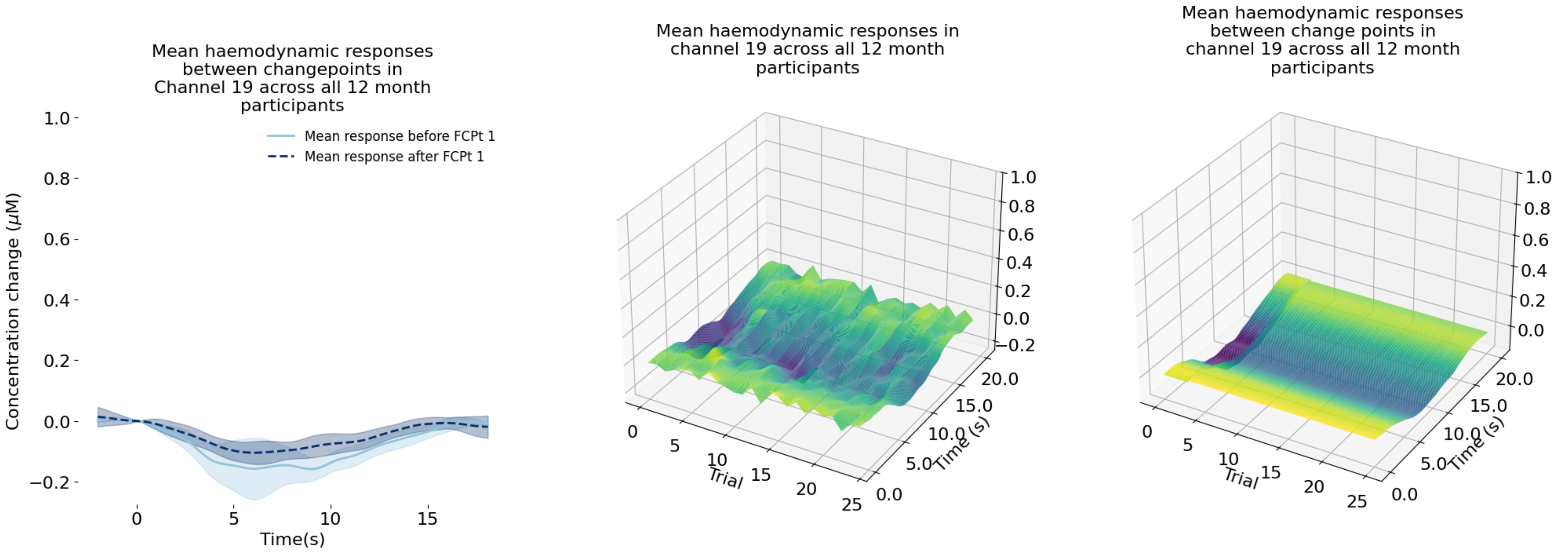
Mean haemodynamic responses between changepoints in Channel 19 across all 12 month participants
Mean response before FCPt 1
Mean response after FCPt 1
Concentration change (μM)
Time(s)
Mean haemodynamic responses in channel 19 across all 12 month participants
Trial
Time (s)
Mean haemodynamic responses between change points in channel 19 across all 12 month participants
Trial
Time (s)

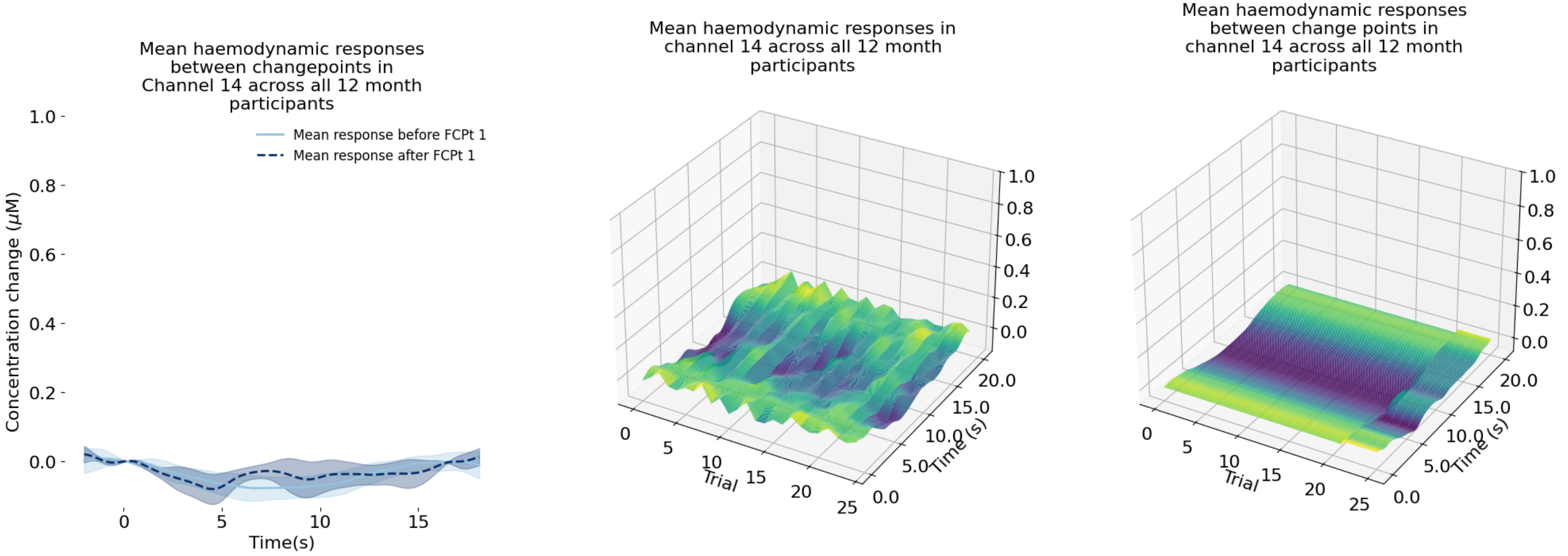
Mean haemodynamic responses between changepoints in Channel 14 across all 12 month participants
Mean response before FCPt 1
Mean response after FCPt 1
Concentration change (μM)
Time(s)
Mean haemodynamic responses in channel 14 across all 12 month participants
Trial
Time (s)
Mean haemodynamic responses between change points in channel 14 across all 12 month participants
Trial
Time (s)

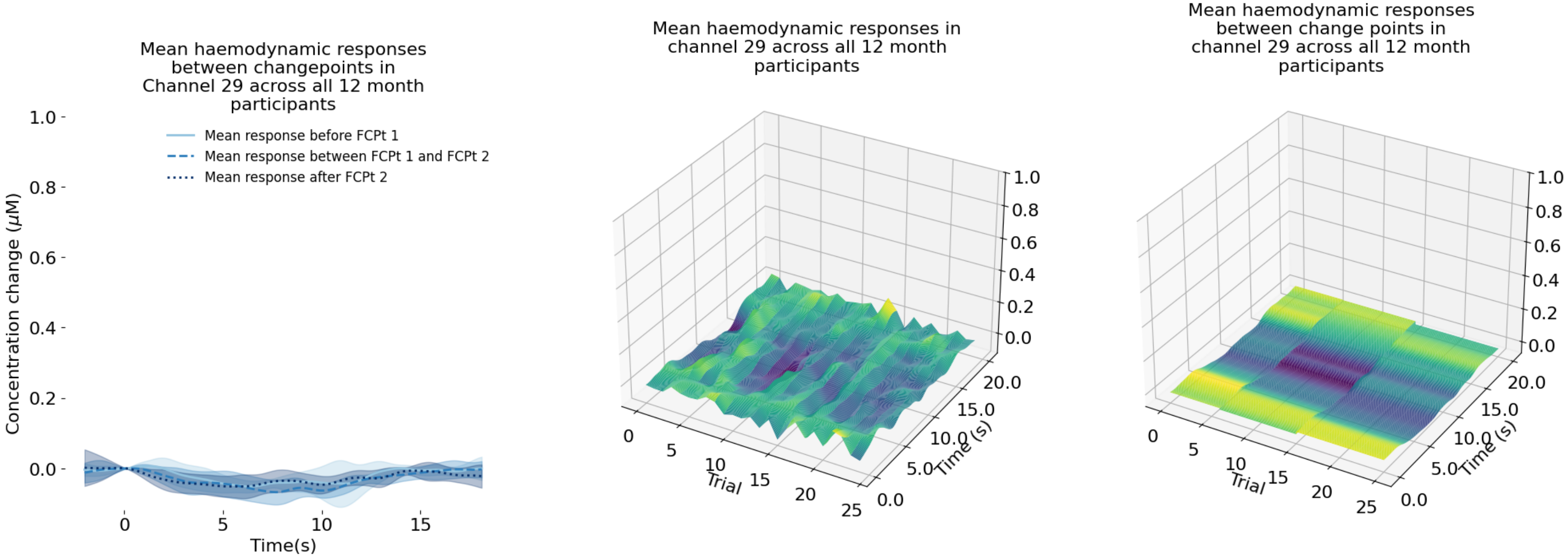
Mean haemodynamic responses between changepoints in Channel 29 across all 12 month participants
Mean response before FCPt 1
Mean response between FCPt 1 and FCPt 2
Mean response after FCPt 2
Concentration change (μM)
Time(s)
Mean haemodynamic responses in channel 29 across all 12 month participants
Trial
Time (s)
Mean haemodynamic responses between change points in channel 29 across all 12 month participants
Trial
Time (s)

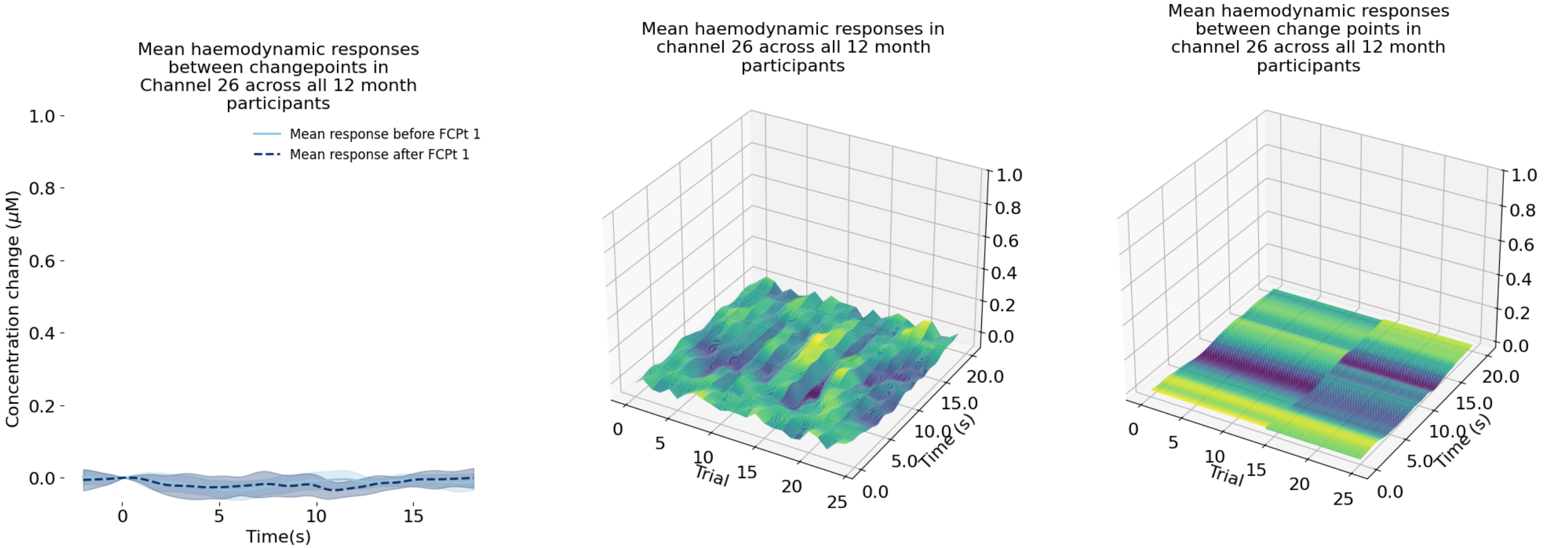
Mean haemodynamic responses between changepoints in Channel 26 across all 12 month participants
Mean response before FCPt 1
Mean response after FCPt 1
Concentration change (μM)
Time(s)
Mean haemodynamic responses in channel 26 across all 12 month participants
Trial
Time (s)
Mean haemodynamic responses between change points in channel 26 across all 12 month participants
Trial
Time (s)

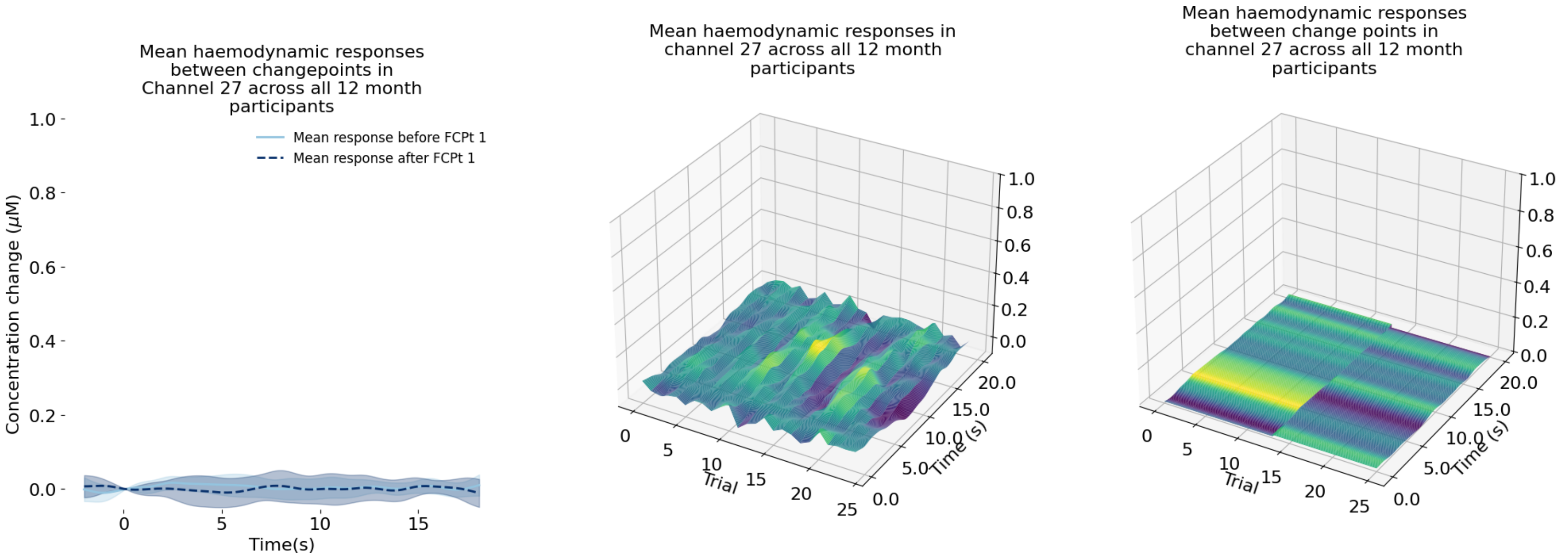
Mean haemodynamic responses between changepoints in Channel 27 across all 12 month participants
Mean response before FCPt 1
Mean response after FCPt 1
Concentration change (μM)
Time(s)
Mean haemodynamic responses in channel 27 across all 12 month participants
Trial
Time (s)
Mean haemodynamic responses between change points in channel 27 across all 12 month participants
Trial
Time (s)

#### Supplementary Materials 9: Channel overlap with univariate methods when using 4 basis functions for the FF method

| Fully functional ($T_N^F$) – 4 basis functions | | | Age-specific ROI | | | Cross-age ROI | | |
|---|---|---|---|---|---|---|---|---|
| Age (months) | Chromophore | $K$ | $n$ | $k$ | $p$-value | $n$ | $k$ | $p$-value |
| 5 | HbO | 11 | *11* | *5* | *0.264* | *11* | *5* | *0.264* |
| | HbR | 8 | *16* | *4* | *0.650* | *16* | *4* | *0.650* |
| | Both | 6 | *7* | *3* | *0.141* | *7* | *3* | *0.141* |
| 8 | HbO | 9 | *14* | *5* | *0.264* | *14* | *5* | *0.264* |
| | HbR | 6 | *11* | *2* | *0.650* | *11* | *2* | *0.650* |
| | Both | 5 | *4* | *2* | *0.141* | *4* | *2* | *0.141* |
| 12 | HbO | 10 | *17* | *7* | *0.264* | *17* | *7* | *0.264* |
| | HbR | 8 | *15* | *4* | *0.650* | *15* | *4* | *0.650* |
| | Both | 6 | *9* | *2* | *0.513* | *9* | *2* | *0.513* |

#### Supplementary Materials 10: The relationship between change point and trial number

Change point trial number plotted against relative change size for BRIGHT data. Relative change size value reported as a proportion of the largest change, with change size itself calculated as the difference between pre- and post-FCPt AUC values.

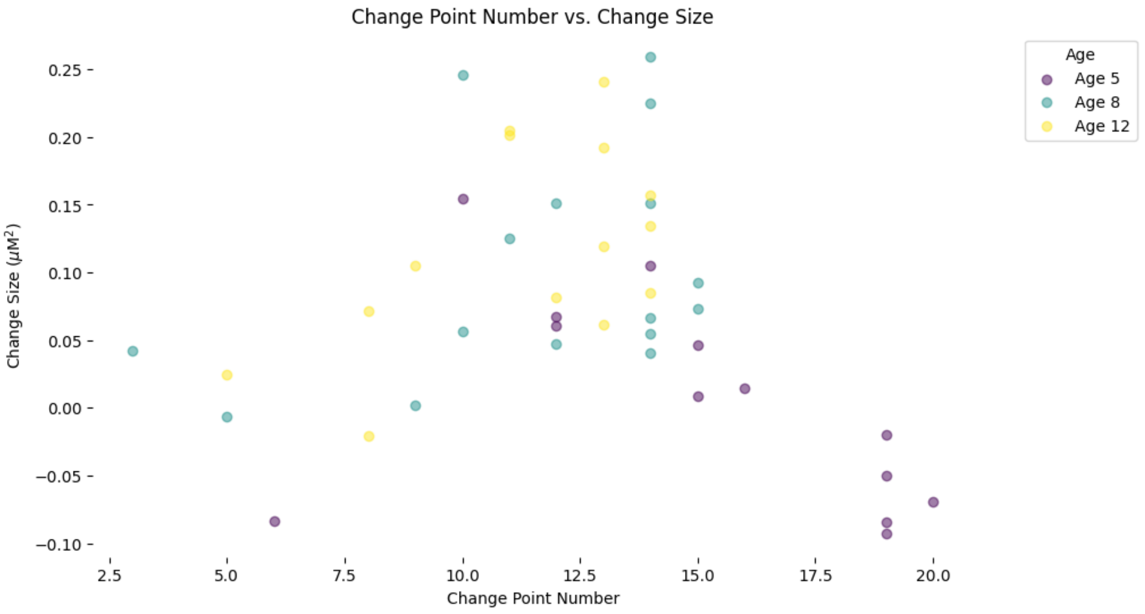

### Supplementary Materials 11: Stability of LR covariance estimations for all channels at 5mo

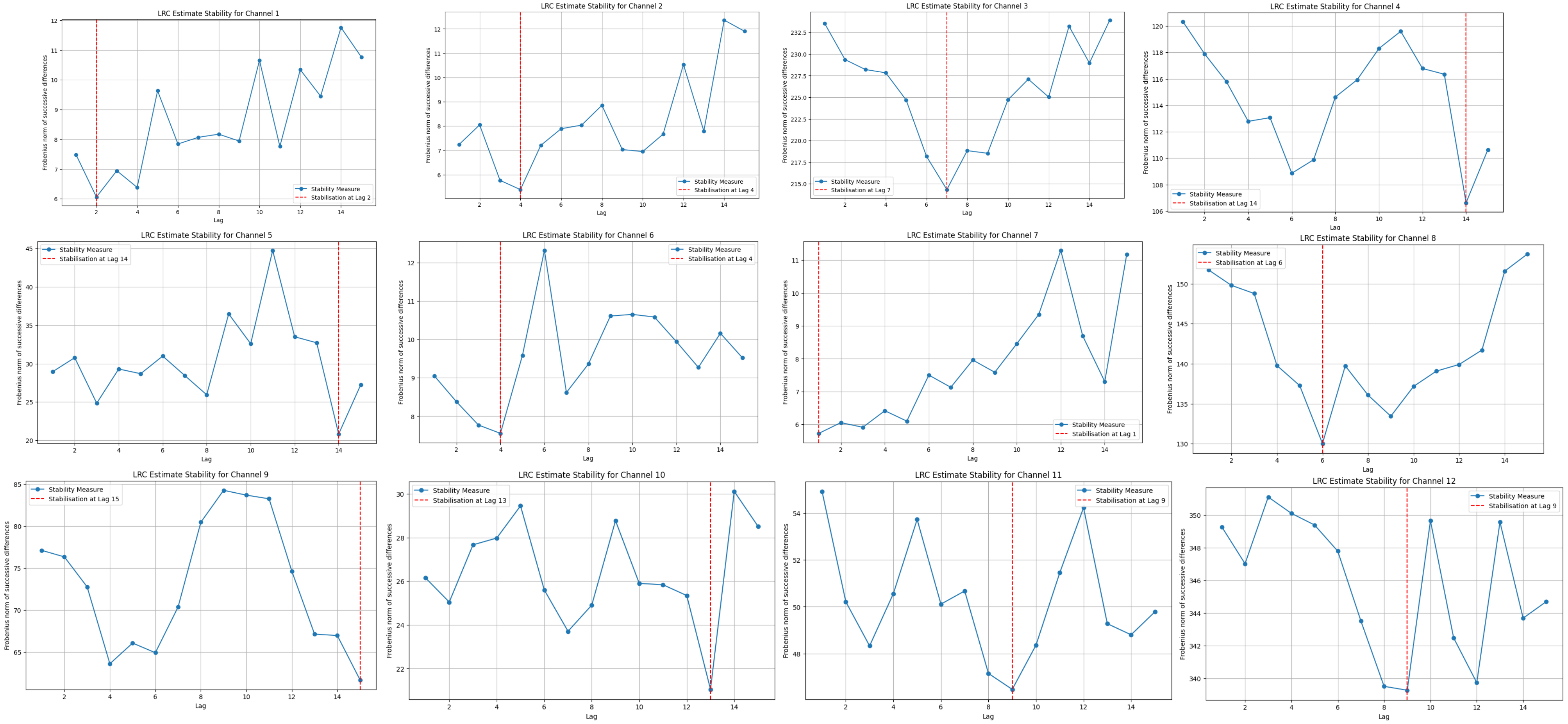

2137

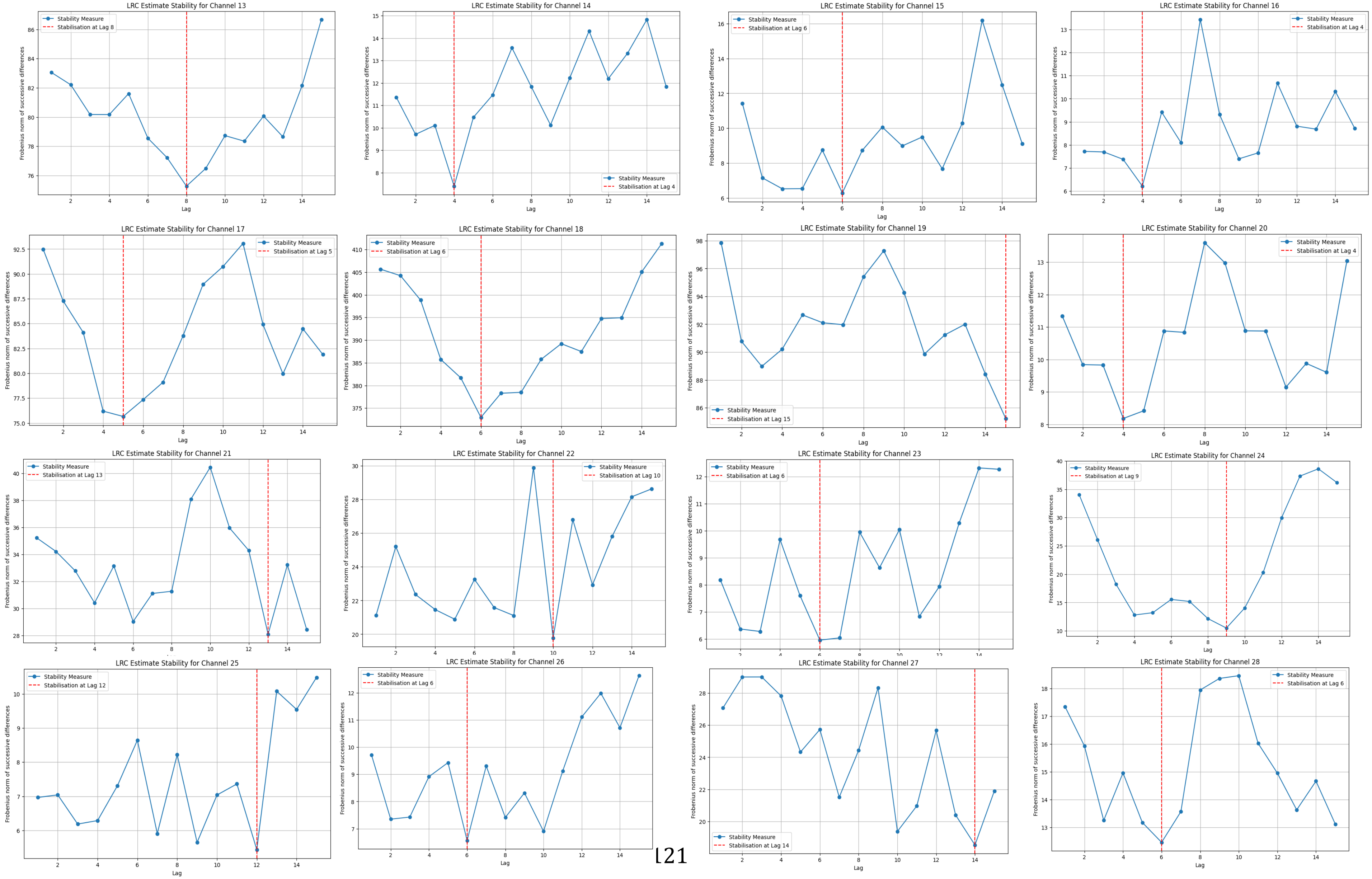
LRC Estimate Stability for Channel 13
Stability Measure
Stabilisation at Lag 8
LRC Estimate Stability for Channel 14
Stability Measure
Stabilisation at Lag 4
LRC Estimate Stability for Channel 15
Stability Measure
Stabilisation at Lag 6
LRC Estimate Stability for Channel 16
Stability Measure
Stabilisation at Lag 4
LRC Estimate Stability for Channel 17
Stability Measure
Stabilisation at Lag 5
LRC Estimate Stability for Channel 18
Stability Measure
Stabilisation at Lag 6
LRC Estimate Stability for Channel 19
Stability Measure
Stabilisation at Lag 15
LRC Estimate Stability for Channel 20
Stability Measure
Stabilisation at Lag 4
LRC Estimate Stability for Channel 21
Stability Measure
Stabilisation at Lag 13
LRC Estimate Stability for Channel 22
Stability Measure
Stabilisation at Lag 10
LRC Estimate Stability for Channel 23
Stability Measure
Stabilisation at Lag 6
LRC Estimate Stability for Channel 24
Stability Measure
Stabilisation at Lag 9
LRC Estimate Stability for Channel 25
Stability Measure
Stabilisation at Lag 12
LRC Estimate Stability for Channel 26
Stability Measure
Stabilisation at Lag 6
LRC Estimate Stability for Channel 27
Stability Measure
Stabilisation at Lag 14
LRC Estimate Stability for Channel 28
Stability Measure
Stabilisation at Lag 6
Lag
Frobenius norm of successive differences

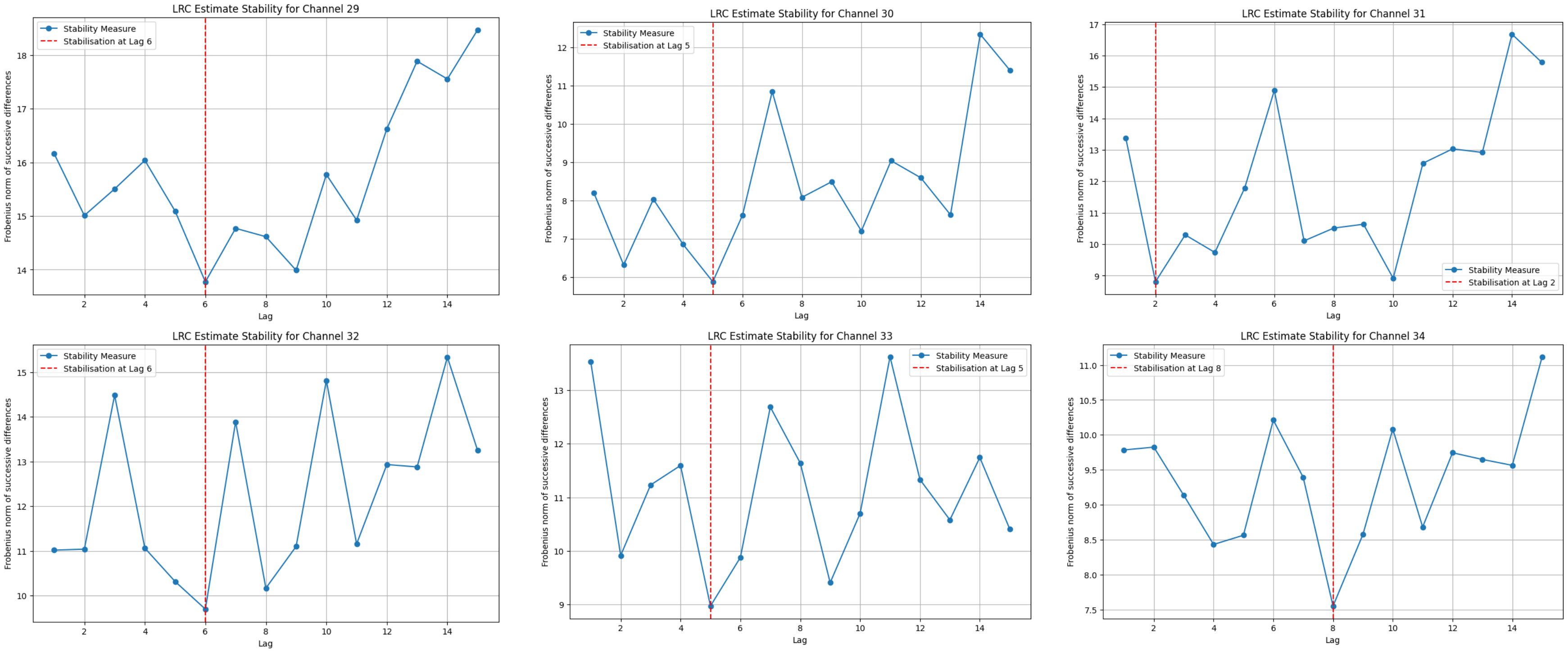

LRC Estimate Stability for Channel 29
Stability Measure
Stabilisation at Lag 6
LRC Estimate Stability for Channel 30
Stability Measure
Stabilisation at Lag 5
LRC Estimate Stability for Channel 31
Stability Measure
Stabilisation at Lag 2
LRC Estimate Stability for Channel 32
Stability Measure
Stabilisation at Lag 6
LRC Estimate Stability for Channel 33
Stability Measure
Stabilisation at Lag 5
LRC Estimate Stability for Channel 34
Stability Measure
Stabilisation at Lag 8
Frobenius norm of successive differences
Lag
